# E^2^M: A Deep Learning Framework for Associating Combinatorial Methylation Patterns with Gene Expression

**DOI:** 10.1101/527044

**Authors:** Jianhao Peng, Idoia Ochoa, Olgica Milenkovic

## Abstract

1

**Motivation:** We focus on the new problem of determining which methylation patterns in gene promoters strongly associate with gene expression in cancer cells of different types. Although a number of results regarding the influence of methylation on expression data have been reported in the literature, our approach is unique in so far that it retrospectively predicts the combinations of methylated sites in promoter regions of genes that are reflected in the expression data. Reversing the traditional prediction order in many cases makes estimation of the model parameters easier, as real-valued data are used to predict categorical data, rather than vice-versa; in addition, our approach allows one to better assess the overall influence of methylation in modulating expression via state-of-the-art learning methods. For this purpose, we developed a novel neural network learning framework termed *E*^2^*M* (Expression-to-Methylation) to predict the status of different methylation sites in promoter regions of several bio-marker genes based on a sufficient statistics of the whole gene expression captured through Landmark genes. We ran our experiments on unquantized and quantized expression sets and neural network weights to illustrate the robustness of the method and reduce the storage footprint of the processing pipeline.

**Results:** We implemented a number of machine learning algorithms to address the new problem of methylation pattern inference, including multiclass regression, canonical correlation analysis (CCA), naive fully connected neural network and inception neural networks. Inception neural networks such as E^2^M learners outperform all other techniques and offer an average prediction accuracy of 82% when tested on 3,671 pan-cancer samples including low grade glioma, glioblastoma, lung adenocarcinoma, lung squamus cell carcinoma, and stomach adenocarcinoma. As an illustrative example, one can increase the prediction accuracy for the methylation pattern in the promoter of gene GATA6 in glioblastoma samples by 20% when using inception rather than simple fully connected neural networks. These performance guarantees remain largely unchanged even when both expression values and network weights are quantized. Our work also provides new insight about the importance of specific methylation site patterns on expression variations for different genes. In this context, we identified genes for which the overwhelming majority of patients exhibit one methylation pattern, and other genes with three or more significant classes of methylation patterns. Inception networks identify such patterns with high accuracy and suggest possible stratification of cancers based on methylation pattern profiles.

**Availability:** The E^2^M code and datasets are freely available at https://github.com/jianhao2016/E2M

## 2 Introduction

Recent studies in computational biology have focused on analyzing multiomics datasets in order to gain a better understanding of the relationships between different dataset components and their unique information content, and to elucidate the relationships between their underlying biological phenomena. This is of particular importance for the case of gene expression data, as there are many genomic and epigenomic factors that influence gene expression (e.g., transcriptional regulation, methylation and histone modification, copy number variation, etc) and as gene expression itself affects almost all aspects of cellular function [7, 30, 1, 22, 5]. One approach to determine to which extent gene expression is determined by or determines other molecular and biochemical modularities is to predict expression values based on associated datasets, such as methylation data [14, 6, 25]. For this task, many learning methods are available, such as logistic regression and deep learning [9, 2]. If the prediction accuracy of the expression values is high, it is reasonable to assume that corresponding data are statistically correlated and that the processes under consideration are biologically interlinked.

Several lines of work in this areas have focused on applying machine learning methods on gene expression data in order to predict clinical outcomes or the dynamics of diseases. In [4], the authors identified a subset of genes whose expression values have strong diagnostic value in cancer staging and survival rate evaluation. The work described in [29] focused on predicting gene expression values based on histone modification data, while taking into account the inherent redundancy present in combined gene expression profiles.

Of particular interest are analyses involving expression and methylation datasets, as methylation is known to be one of the key regulators of expression [15] (see Figure 1). Methylation is a common epigenetic modification [24] that plays an important role in tumorigenesis and cancer progression. The methylation process alters the chemical structure of Cytosine or Guanine at *CpG sites*, which often cluster within *CpG islands* in promoter regions of genes. Although a CpG island may contain tens of CpG sites, it has been a standard practice to only report the *thresholded cumulative methylation effect* of the sites and declare a binary methylation state of a gene (methylated or unmethylated). In order to associate DNA methylation with gene expression changes, the authors of [25] proposed a supervised learning method termed ME-Class (Methylation-based Expression Classification) for predicting expression changes based on soft methylation features. The goal of the aforementioned study was to assess the raw predictive power of methylation data, rather than to determine which *combinations of methylation sites* truly contribute to the observed expression profiles. On the other hand, authors of [18] proposed an attention model which utilized both the expression data and CpG sites distance information to predict methylation states of one CpG site. Our work hence centers on a higher-order and in-depth analysis of the mutual relationship between expression and *methylation site patterns* in the context of pan cancer data analysis. The natural approach to pursue within this framework is retrospective analysis, which amounts to inferring methylation patterns (i.e., discrete states) from expression values (i.e., continuous observations).

**Fig. 1:**
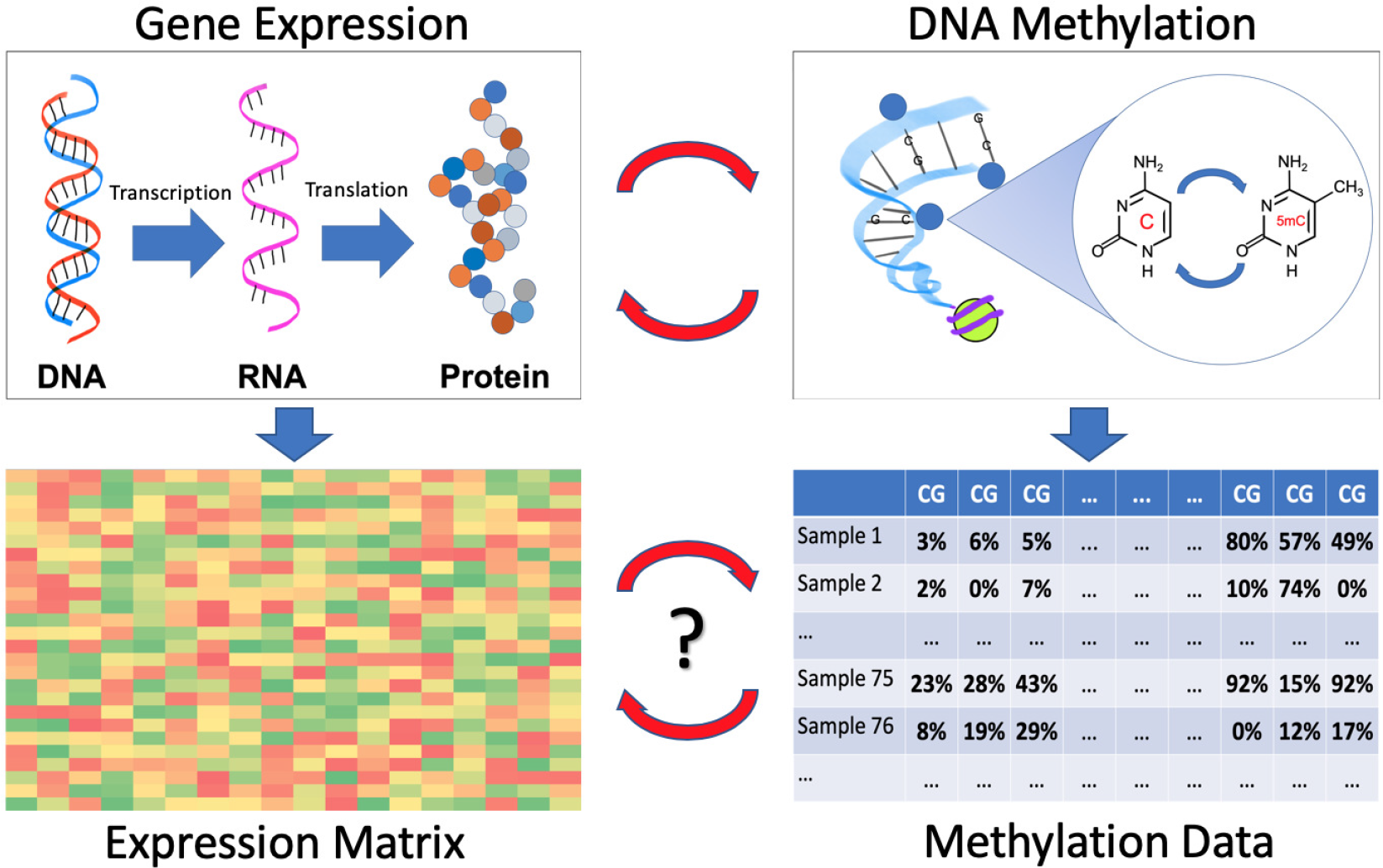
Associations between gene expression and DNA methylation and their corresponding datasets.

Our technical contributions are three-fold. First, we introduce the problem of correlating binary methylation patterns with the expressions of Landmark genes [8]. This significantly reduces the complexity of the problem and simultaneously performs denoising of expression values, as the set of Landmark genes is rather small (≈ 1000) and selected for its predictive power for whole-genome expressions. Second, we propose a new *inception network* [31] architecture for deep learning, termed E^2^M, which performs retrospective classification. The prediction accuracy of the E^2^M network is up to 20% better than that offered by traditional multi-class logistics regression and three-layer fully connected networks. Third, we demonstrate that our learning framework can operate with quantized parameter sets and significantly compressed datasets. Large-depth neural networks are known to be more robust to noise [3, 11] than shallow networks, but their practical application is limited by their large storage footprint. To show that this problem may be easily mitigated via quantization, we demonstrate that 8-bit uniformly quantized expression values and 16-bit quantized network weights cause negligible degradations in recovering underlying methylation patterns for almost all tested examples.

Our analysis also reveals that methylation patterns are gene-dependent and that they influence the expression dynamics differently for different types of cancers. In some cases, such as lung cancer, the methylation patterns in genes such as MGMT, ATM, GATA6 and KRAS differ significantly, while the methylation patterns in the MLH1 and CASP8 genes show little variation. For some other cases, such as brain cancer, most genes show very similar methylation patterns, except for GATA6. Furthermore, some genes, such as TP53, have a unique methylation pattern for a specific cancer type, and the patterns vary little across cancer types.

## 3 Methods

We start our exposition by describing the datasets used in our analysis, and then proceed with a discussion of existing and new methods suitable for addressing the prediction problem at hand.

### 3.1 Data Description

The problem of associating different types of multiomics data has received significant attention in the computational biology community [17, 10]. To assess the performance of the proposed framework E^2^M, we restrict our attention to human cancer cell expression and methylation data. Our goal is to predict methylation patterns from gene expression values.

There are over 20,000 genes in the human genome, and using their gene expression values directly in any machine learning task would lead to undetermined problems and overfitting issues due to redundancy and small sample set sizes. Hence, the first step in our approach is to perform dimensionality reduction. To this end, we focus on expression levels of so-called L1000 Landmark genes, comprising 978 genes. This subset of genes has been carefully selected by the NIH LINCS project (http://lincsportal.ccs.miami.edu/dcic-portal/) for its good predictive capabilities for the whole genome expression profile. It has also been demonstrated in [8] that deep networks can accurately recover the whole genome expression profile using only L1000 expression information. An additional advantage of using L1000 genes is that LINCS provides efficient and low-cost assays for measuring the expression of these genes.

Gene expression data is available in multiple formats. High throughput (HT) sequence counts (i.e., raw counts of gene transcripts) are the most frequently used measurements for describing expression, and all other data representations are derived from these counts. However, since DNA transcripts have different lengths and concentrations, the raw counts may not accurately reflect the relative expression level. To mitigate this problem, raw counts are transformed into Fragments Per Kilobase of Transcript per Million (FPKM) mapped reads values, computed as:

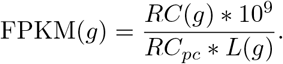

Here, *RC*(*g*) and *L*(*g*) represent the raw counts of reads covered by gene *g* and the length of gene *g*, respectively, while *RC*_*pc*_ denotes the total raw count of reads mapped to all protein coding genes.

In all our experiments, we use FPKM readings of the 978 Landmark genes as inputs. The actual FPKM data was retrieved from The Cancer Genome Atlas (TCGA) project (https://cancergenome.nih.gov/). We selected five different cancer (sub)types: lower grade glioma (LGG), glioblastoma (GBM), lung adenocarcinoma (LUAD), lung squamous cell carcinoma (LUSC) and stomach adenocarcinoma (STAD), and extracted all available Landmark gene expression datasets (https://portal.gdc.cancer.gov/, downloaded March 2018).

Since most CpG sites are naturally methylated inside a gene, we focus on methylation measurements of individual CpG sites within promoter regions. Genes of interest include well-known cancer biomarkers, MGMT, MLH1, ATM, GATA6, CASP8, KRAS and TP53 (see the Supplementary Material, Section 1). Information available at TCGA includes methylation microarray values for 7, 6, 4, 2, 3, 6 and 3 CpG sites in the promoter region of these genes, respectively. Since genes in different cancer types have nonuniform methylation levels as measured in terms of coverage of the methylated sites, the recorded readings only capture the percentage of methylated CpG sites (bottom, right-hand format in Figure 1). We convert these percentages into binary values by thresholding at 10%, as suggested in multiple prior works [19]. The output of this preprocessing step is an *m*-dimensional binary vector, where *m* is the number of CpG sites in the promoter region of the underlying gene.

As one needs to ensure that both methylation and expression data are available for the same sample, the test data included 511 samples from LGG, 126 from GBM, 511 from LUAD, 500 from LUSC, and 375 from STAD. This amounts to a total of 637 samples for brain cancer (LGG and GBM), and 1011 samples for lung cancer (LUAD and LUSC). In all subsequent analyses, these datasets were split into training and test sets in a 80%-20% proportion.

In summary, we used 978-length positive real-valued vectors containing the FPKM counts of Landmark genes as inputs of a learner tasked with predicting binary methylation patterns with *m* entries, corresponding to our preselected biomarker genes.

### 3.2 Mathematical Approaches

There exists many methods that may be used for associating different types of multiomics data. Among these, the most frequently used approaches include canonical correlation analysis (CCA) [13], multiclass regression (MR) and fully connected neural networks (FCNN) [20]. However, these techniques have limitations that make them unsuitable for the problem at hand, as described in what follows. Note that to demonstrate the drawbacks of CCA, MR, and FCNN, we actually applied these methods on the curated datasets and reported their performance.

#### 3.2.1 Canonical Correlation Analysis

Canonical correlation analysis (CCA) is widely used to infer linear relationships between two correlated random measurements (e.g, random vectors) *X* ∈ *R*^*n*^ and *Y* ∈ *R*^*m*^. In our setting, *n* = 978, and each *X* corresponds to the gene expression profile of a cancer patient, while *m* ≤ 7 and each *Y* corresponds to a binary DNA methylation pattern of the same cancer patient. The CCA objective formally reads as:

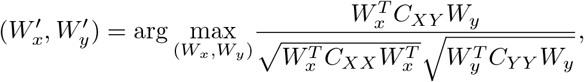

where *C*_*XY*_ is the empirical covariance matrix of *X* and *Y* computed using all *s* ≤ 1011 available samples for each individual cancer type. Then, *W*_*x*_ ∈ *R*^*s*^, *W*_*y*_ ∈ *R*^*s*^. Intuitively, CCA aims to find a subspace such that the projections *W*_*x*_ and *W*_*y*_ of the random vectors *X* and *Y*, respectively, have the largest possible correlation. This optimization process can be repeated sequentially to obtain a sequence of pairs of random vectors (*W*_*x*_, *W*_*y*_) that are mutually uncorrelated, akin to what is standardly done in eigendecomposition problems.

CCA may be directly applied to our data, but it does not provide a constructive means for inferring methylation patterns based on expression; furthermore, it can only identify *linear dependencies* between two data sample matrices. In addition, given that we have more features (978 genes) than samples (less than 600 for each cancer subtype), highly-correlated projections arise naturally and are easy to identify through the described optimization process. Table 1 illustrates this point for the case of CCA analysis on the MGMT gene, and all cancer subtypes. As expected, canonical correlation values in this case exceed 0.94.

**Tab. 1:** The first three canonical correlation values for the MGMT gene according to cancer subtype.

|  | LGG | GBM | BRAIN | LUAD | LUSC | LUNG | STAD |
| --- | --- | --- | --- | --- | --- | --- | --- |
| 1st | 0.99 | 1.00 | 0.99 | 0.99 | 0.99 | 0.96 | 0.99 |
| 2nd | 0.99 | 1.00 | 0.98 | 0.99 | 0.99 | 0.96 | 0.99 |
| 3rd | 0.99 | 1.00 | 0.98 | 0.99 | 0.99 | 0.94 | 0.99 |

#### 3.2.2 Multiclass Logistic Regression

A methylation pattern is represented by a binary vector of length *m*, where we recall that *m* denotes the number of CpG sites in the promoter region of a gene of interest. Hence, a methylation pattern corresponds to one of 2^*m*^ possible binary vectors (classes). Since in our case *m* ≤ 7, multiclass logistic regression is a natural candidate for prediction.

Let *ω*_*y*_ be the weight of the class label *y* ∈ [0 : 2^*m*^ − 1]. The posterior probability of the class label given a particular expression profile *X* = *x* may be written as

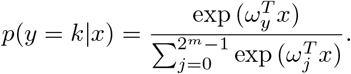

Under the assumption that all samples are drawn independently from each other, the goal is to maximize the product of *p*(*y*|*x*) over all pairs (*x*, *y*).

Multiclass logistic regression is only guaranteed to perform well for linearly separable data [20]. Due to the large dimension of gene expression vectors, it is computationally difficult to determine if the datasets of interest are linearly separable or not. To address this problem, we first performed dimensionality reduction via principal component analysis (PCA) and then visually inspected the data. Figures 2 and 3 depict the obtained results for two principle components of training and test samples, for the LGG and LUNG (e.g., the combination of LUAD and LUSC) cancer subtype(s), respectively. Only results for the GATA6 gene are shown; the results for other cancer types and genes are available in section 2 of the Supplementary Material. It can be observed that the chosen methylation patterns are not linearly separable. For LGG, we observed two classes of methylation patterns (light and dark blue points) that exhibit a small degree of separability, whereas for LUNG, the two patterns are superimposed onto each other. Hence, multiclass logistic regression is not expected to perform well on most of the data involving multiple labels (see Table 2, and in particular, the values corresponding to cancer types LGG and LUNG).

**Tab. 2:** Accuracy of Multiclass Regression (MR) and Fully Connected Neural Network (FCNN) methods according to cancer data type, for gene GATA6.

|  | LGG | GBM | BRAIN | LUAD | LUSC | LUNG | STAD |
| --- | --- | --- | --- | --- | --- | --- | --- |
| MR | 0.83 | 0.38 | 0.79 | 0.71 | 0.98 | 0.70 | 0.96 |
| FCNN | 0.84 | 0.35 | 0.82 | 0.71 | 0.99 | 0.85 | 0.96 |

**Fig. 2:**
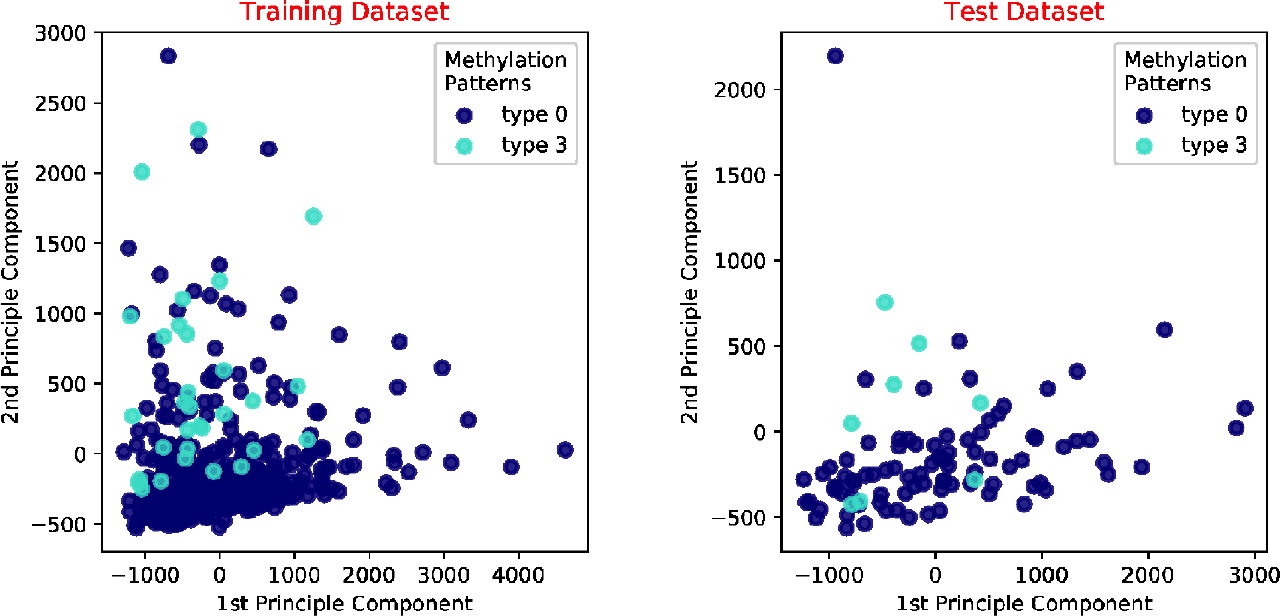
Visualization of the first two principal components of LGG cancer data for the GATA6 gene.

**Fig. 3:**
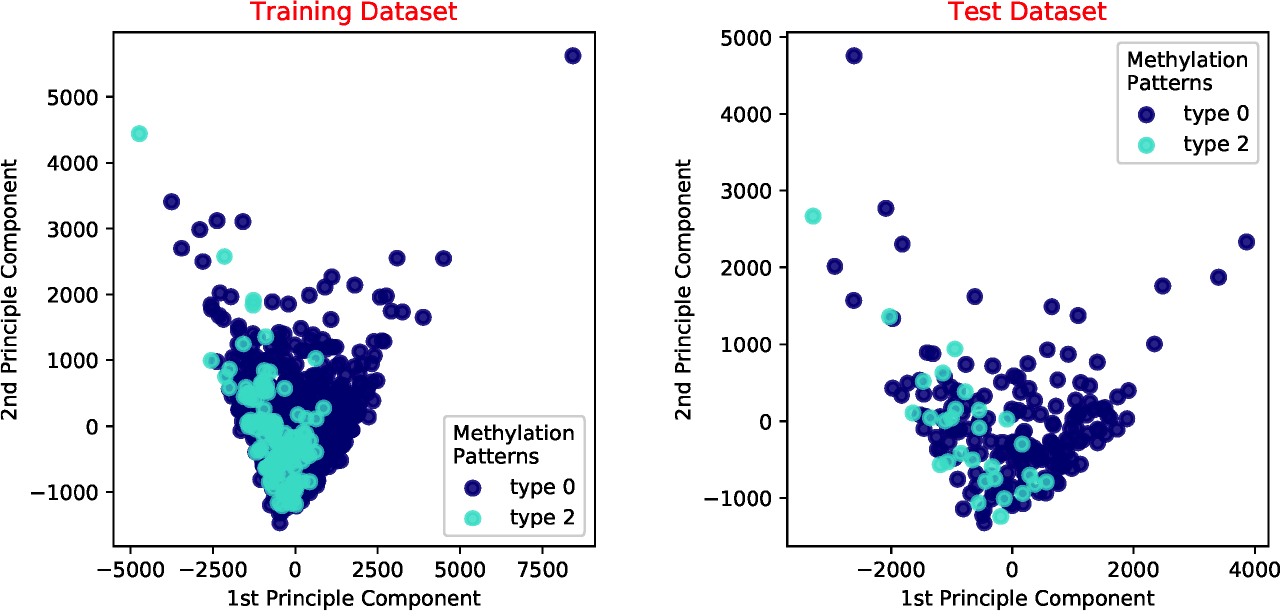
Visualization of the first two principal components of LUNG cancer data for the GATA6 gene.

#### 3.2.3 Fully Connected Neural Network

FCNNs are a method of choice for many classification tasks [23] as they do not require data to be linearly separable for practically good performance. Figure 4 depicts the structure of a classic three-layer FCNN. Each neuron in the network uses a nonlinear activation function on a linear combination of the outputs from the previous layer. The activation function introduces nonlinearities into the network model and increases its expressive power compared to linear models such as logistics regression or support vector machines (SVM) [20]. There are many choices for the nonlinear function, but we restrict our attention to Rectified Linear Units (ReLUs) as they have constant gradients during training and are commonly used in practice.

**Fig. 4:**
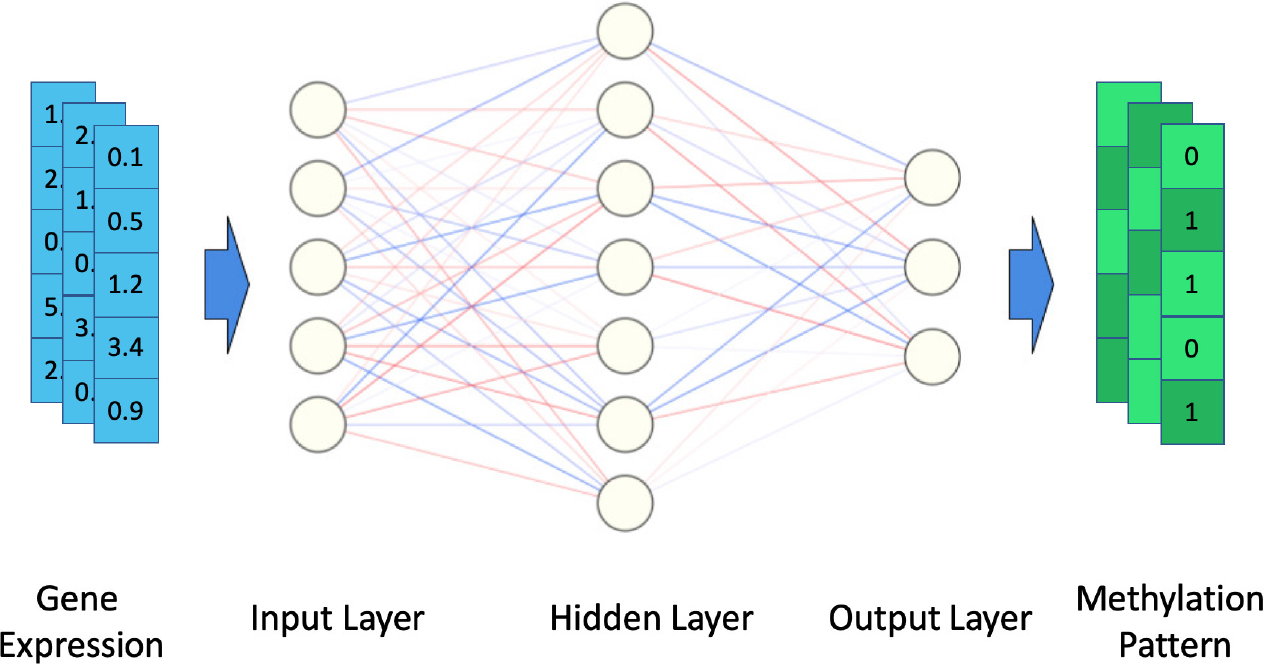
Architecture of a 3-layer fully connected neural network.

When the activation function equals the identity function and the loss is chosen appropriately, the resulting shallow network is equivalent to a linear model (e.g., a linear model such as logistic regression). Hence, one expects fully connected neural networks to be more accurate in finding the correct association between the gene expression profiles and the methylation patterns. Nevertheless, fully connected neural networks may fail to take into account possible correlations between genes, which considerably compromises their performance (see Table 2).

### 3.3 Inception-Based Deep Network E^2^M

To address the issues present in the previously described methods, we propose a new method for mining associations between methylation patterns and Landmark gene expressions, termed E^2^M. The approach is based on a novel neural network learning framework centered around so-called inception neural network learners [31].

Inception networks include modules that mitigate certain problems encountered in simple fully connected networks (Figure 5). One such problem pertains to capturing long-distance interactions between genes and correlations between their expressions, which can be addressed in part by adding convolutions. However, since the interaction distance is not known a priori, it cannot be used to inform the choice of the depth of convolution. Inception modules therefore include multiple convolutions of different depths at the same layer. Another component is the filter stack, which aggregates the outputs of the convolutions and pooling layers, and feeds them to the next network layer. These features make inception networks more robust and allow them to converge faster than traditional convolutional neural networks.

**Fig. 5:**
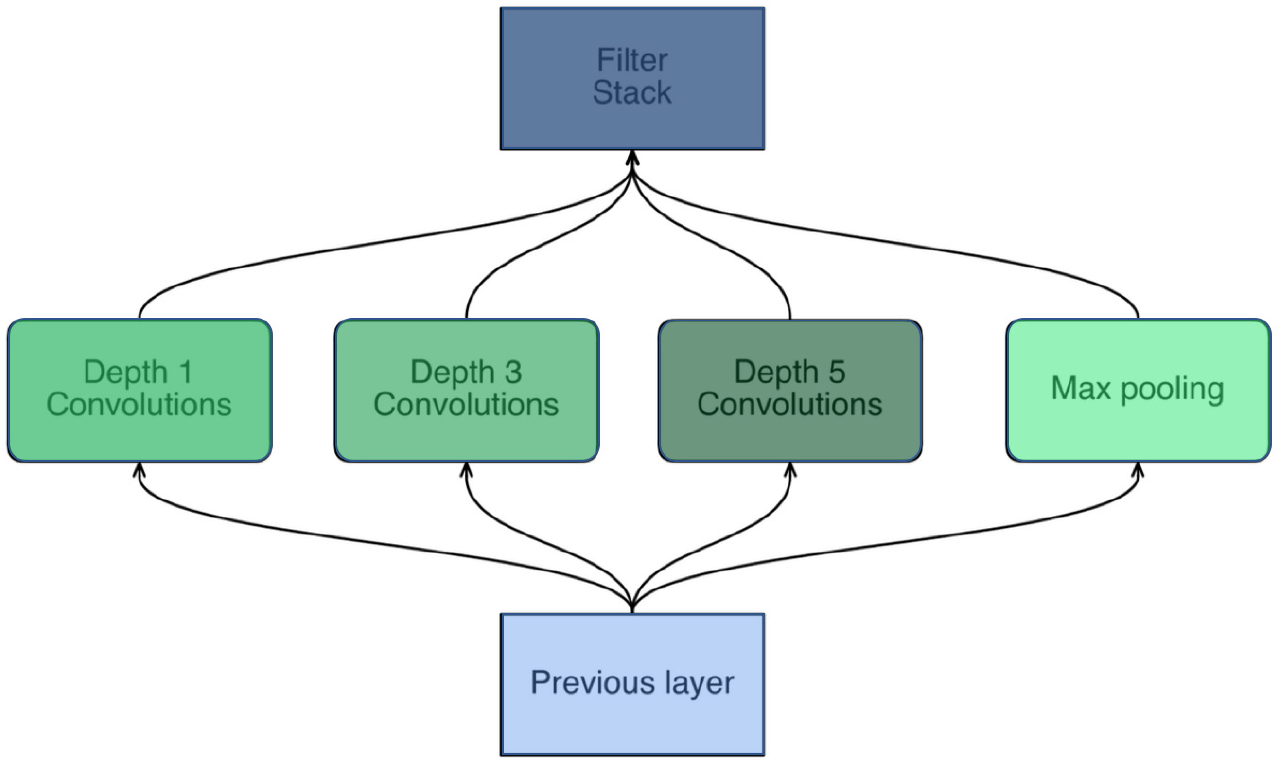
Architecture of an inception module.

**Fig. 6:**
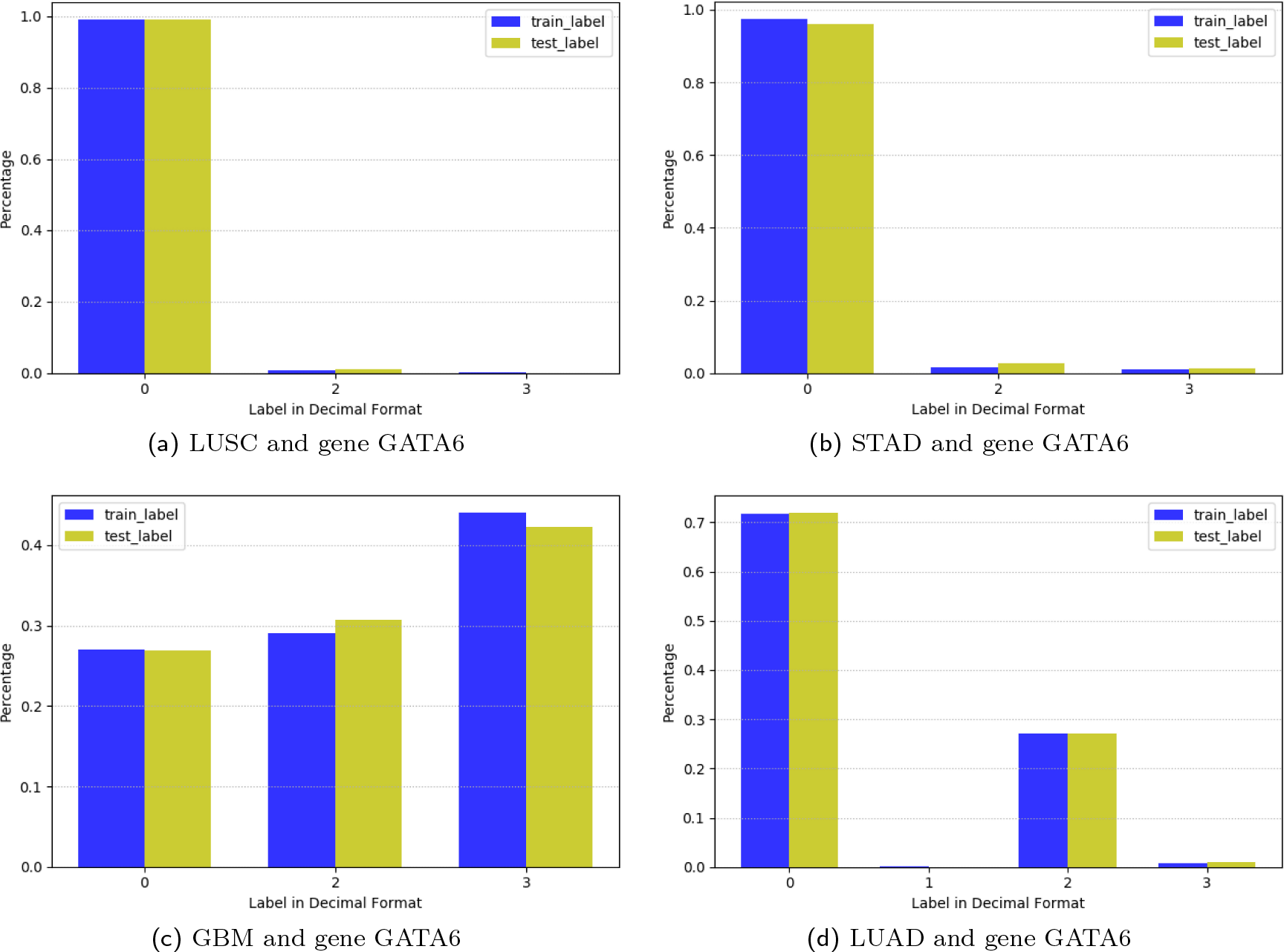
Histograms of methylation patterns present in the promoter of gene GATA6 for cancer types LUSC, b) STAD, c) GBM, and d) LUAD.

E^2^M includes two 1D convolution and maxpooling layers that increase the number of filters and reduce the dimension of the feature space. These layers are followed by two inception modules of the form shown in Figure 5. This structure is terminated by a fully connected layer that “flattens” the output (i.e., converts a matrix into a vector via column concatenation) and a softmax layer that is used to make the predictions. We point out that the largest contribution to the storage footprint of neural network learners comes from the fully connected layer that is densely connected. As a result, even though the E^2^M structure may appear to have higher description complexity than a fully connected network, it actually uses fewer parameters than a 3-layer fully connected network, and is consequently faster, more compact and less prone to overfitting.

### 3.4 Quantized E^2^M and Gene Expression Values

As the depth of neural networks increases, their storage and running costs become prohibitively high for many practical applications. While training a neural network model, it is important to maintain high data precision in order to propagate accurate gradient information. Nevertheless, it has been shown that quantizing the weights of an already trained neural network, if done properly, only slightly degrades its performance and occasionally even improves it [12, 21]. In addition, since the FPKM gene expression readings are normalized to convey relative expression differences among genes, it is unnecessary to force them to be of high precision. Furthermore, original counts can always be recomputed from their corresponding FASTQ files, which are standardly stored in a lossless manner. Hence, in our experiments, we also consider quantizing the input gene expression data.

For network quantization, we identify the largest and smallest weights in each layer, and bin all other weights according to uniform quantization rules, with 16–, 8–, or 4–bit level representations. Note that full precision floating point data in our setting are represented with 64 bits. To perform network weight quantization, we use a built-in function of TensorFlow that allows for performing quantized weights multiplication and addition, without mapping them back to the floating point format. To quantize gene expression values, we select a cut-off threshold for the top 5% highest-reading genes; readings between 0 and the cut-off value are uniformly quantized at the 16–, 8–, or 4–bit level.

## 4 Results

In what follows, we present our findings regarding associations between methylation patterns and Landmark expression profiles for different cancer types and genes. In the process, we first identify through extensive data analysis the most suitable inception network parameters, module and layer numbers. Then, we proceed to compare the performance of the proposed E^2^M framework to that of multiclass logistic regression and a 3-layer fully connected neural network. We then describe the effects of quantization of network weights and input expression data under the E^2^M approach. Our discussion concludes with an interpretation of the uncovered biological phenomenon.

### 4.1 E^2^M Parameter Selection

A detailed description of the structure of the newly proposed E^2^M learner can be found in Table 3. The reported parameters were chosen using cross-validation methods on the entirety of the training data described in the previous section, by splitting it into a 90 : 10 proportion. Subsequently, for 36 sets of parameters, we trained the network on the 90% training set and tested it on the remaining 10% dataset. The selected parameters were the best-performing ones under the validation setting.

**Tab. 3:**
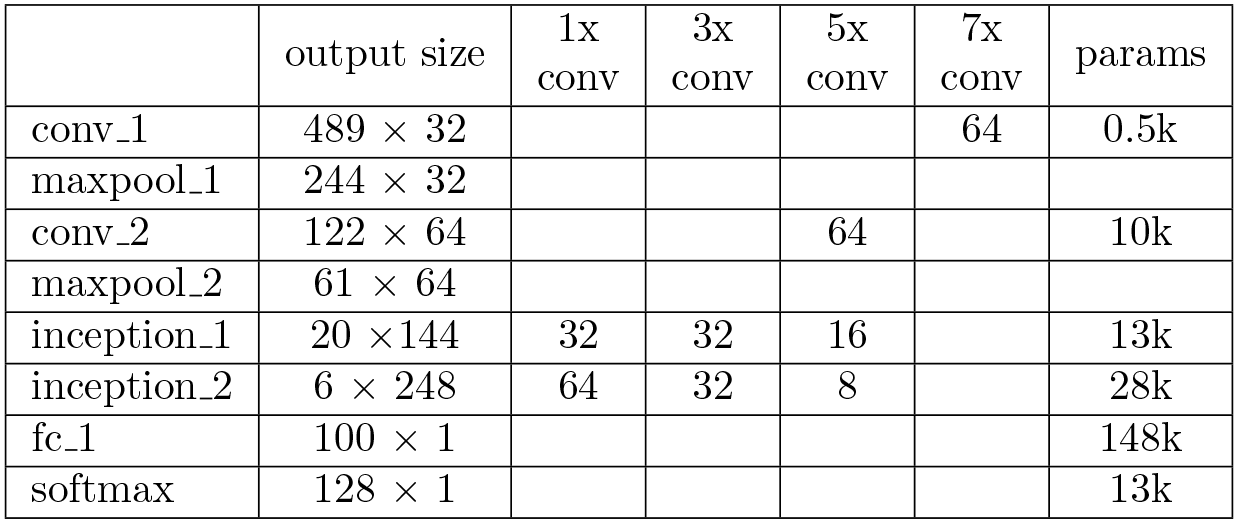
Detailed structure of the proposed E^2^M framework with 128 classes. The entries in column numbers 3 6 correspond to the number of parallel convolutions (filters). The last column lists the number of parameters in each layer of the learner.

|  | output size | 1x<br>conv | 3x<br>conv | 5x<br>conv | 7x<br>conv | params |
| --- | --- | --- | --- | --- | --- | --- |
| conv_1 | $489 \times 32$ | | | | 64 | 0.5k |
| maxpool_1 | $244 \times 32$ | | | | | |
| conv_2 | $122 \times 64$ | | | 64 | | 10k |
| maxpool_2 | $61 \times 64$ | | | | | |
| inception_1 | $20 \times 144$ | 32 | 32 | 16 | | 13k |
| inception_2 | $6 \times 248$ | 64 | 32 | 8 | | 28k |
| fc_1 | $100 \times 1$ | | | | | 148k |
| softmax | $128 \times 1$ | | | | | 13k |

### 4.2 Performance Analysis of E^2^M

We compared the performance of the chosen E^2^M framework with MRs and 3-layer FCNNs, and summarized the results in Table 4. The results correspond to all chosen cancer types and gene GATA6; the results pertaining to the remaining selected genes may be found in Section 5 of Supplementary Material (Table 3 and 4).

**Tab. 4:** Comparison of accuracies of different prediction methods applied to gene GATA6 and all considered cancer types. The best results are highlighted in bold font. Note that the accuracy of random guessing is 0.25, as the promoter of GATA6 contains only two methylation sites.

|  | MR | 3-layer<br>FCNN | E<sup>2</sup>M |
| --- | --- | --- | --- |
| LGG | 0.83 | 0.84 | **0.90** |
| GBM | 0.38 | 0.35 | **0.58** |
| BRAIN | 0.79 | **0.82** | 0.81 |
| LUAD | 0.71 | **0.71** | 0.69 |
| LUSC | 0.98 | 0.99 | **0.99** |
| LUNG | 0.70 | **0.85** | 0.84 |
| STAD | 0.96 | 0.96 | **0.96** |
| Average | 0.77 | 0.79 | **0.82** |

As it may be observed from the table, at least one of the two nonlinear network models always outperforms logistics regression for all tested cases. The reason, as explained in the previous section, is that the data used may not be linearly separable. Once again, we point out the results for LGG and LUNG cancers. From Figures 2 and 3, it is apparent that LGG data is easier to separate in the principal component space than LUNG data. In particular, for the LGG dataset, MR and FCNN perform very similarly (0.83 and 0.84, respectively), while E^2^M offers the best performance (0.90). On the other hand, for LUNG data, the non-linear models outperform the logistic regression model by 15%. From the table, we also observe that in most cases our proposed inception network E^2^M has higher prediction accuracy than FCNN. For example, for LGG and GBM, E^2^M exhibits a 6% and 23% improvement over FCNN, respectively. Whenever FCNN outperforms E^2^M, the difference in performance is only about 1 − 2%. Hence, E^2^M offers the best average performance among all the methods under consideration (additional results along the same line may be found in section 5 of the Supplementary Material).

Given that E^2^M offers the best average performance of all tested methods, we henceforth restrict our attention to this method only. Table 5 provides the performance results of E^2^M for all cancer types and all selected genes (the biological relevance of the bold font and italic values will be described in the Discussion section).

**Tab. 5:** Prediction accuracy of E^2^M for all considered cancer types and genes. The row RG lists the accuracy of random guessing.

| Gene | MGMT | MLH1 | ATM | GATA6 | CASP8 | KRAS | TP53 |
| --- | --- | --- | --- | --- | --- | --- | --- |
| RG | (0.008) | (0.016) | (0.0625) | (0.25) | (0.125) | (0.016) | (0.125) |
| LGG | 0.56 | 0.75 | *0.99* | *0.90* | *0.99* | *0.84* | *0.96* |
| GBM | **0.31** | 0.61 | *0.92* | **0.58** | *0.88* | *0.80* | *1.00* |
| BRAIN | 0.53 | 0.69 | *0.98* | **0.81** | *0.98* | *0.84* | *0.98* |
| LUAD | *0.83* | 0.42 | 0.73 | 0.69 | **0.62** | 0.38 | *0.84* |
| LUSC | **0.64** | 0.65 | *0.94* | *0.99* | **0.63** | *0.92* | *0.99* |
| LUNG | 0.71 | 0.53 | *0.85* | *0.84* | **0.58** | 0.68 | *0.90* |
| STAD | 0.65 | **0.44** | *0.84* | *0.96* | **0.63** | *0.79* | *0.95* |

### 4.3 Performance of the Quantized E^2^M Method

Table 6 shows an example of how quantization of network parameters and input data influences the prediction accuracy of E^2^M for all considered cancer types and genes. We only report on the results obtained using 16-bit uniform quantization of all network weights and 8-bit uniform quantization on the expression input data (see section 6 in Supplementary Material for results with other quantization levels). For ease of data interpretation, the numbers in parenthesis correspond to the prediction accuracy values without quantization. As it may be observed, there is almost no degradation in the performance of the quantized E^2^M method, and in some cases, quantization even improves the prediction results. The only degradations observed are for GBM – gene GATA6, LUAD – gene ATM, and LUNG – gene CASP8. An explanation for this finding is described in the Discussion section.

**Tab. 6:** Prediction accuracy of E^2^M with network weights quantized to 16 bits, and expression inputs quantized to 8 bits.

|  | MGMT | MLH1 | ATM | GATA6 | CASP8 | KRAS | TP53 |
| --- | --- | --- | --- | --- | --- | --- | --- |
| LGG | 0.55<br>(0.56) | 0.71<br>(0.75) | 0.99<br>(0.99) | 0.90<br>(0.90) | 0.99<br>(0.99) | 0.80<br>(0.84) | 0.97<br>(0.96) |
| GBM | 0.24<br>(0.31) | 0.52<br>(0.61) | 0.96<br>(0.92) | **0.36**<br>(0.58) | 0.92<br>(0.88) | 0.86<br>(0.80) | 1.00<br>(1.00) |
| BRAIN | 0.47<br>(0.53) | 0.69<br>(0.69) | 0.98<br>(0.98) | 0.76<br>(0.81) | 0.98<br>(0.98) | 0.85<br>(0.84) | 0.97<br>(0.98) |
| LUAD | 0.82<br>(0.83) | 0.43<br>(0.42) | **0.64**<br>(0.73) | 0.69<br>(0.69) | 0.55<br>(0.62) | 0.41<br>(0.38) | 0.81<br>(0.81) |
| LUSC | 0.61<br>(0.64) | 0.61<br>(0.65) | 0.96<br>(0.94) | 0.99<br>(0.99) | 0.61<br>(0.63) | 0.92<br>(0.92) | 0.99<br>(0.99) |
| LUNG | 0.73<br>(0.71) | 0.54<br>(0.53) | 0.84<br>(0.85) | 0.81<br>(0.84) | **0.44**<br>(0.58) | 0.67<br>(0.68) | 0.91<br>(0.90) |
| STAD | 0.61<br>(0.65) | 0.37<br>(0.44) | 0.85<br>(0.84) | 0.95<br>(0.96) | 0.59<br>(0.63) | 0.76<br>(0.79) | 0.96<br>(0.95) |

We also remark that for the three cases with compromised performance under quantization, the predicted patterns are at small Hamming distance from the correct one. In other cases, like GBM and gene GATA6, the Hamming distance between predicted patterns is at least two, and hence E^2^M may be quantized with even fewer bits while preserving prediction accuracy.

In conclusion, aggressive quantization in most cases leads to small prediction performance degradation, while providing significant storage savings: in the example provided, input data is reduced to 1/8 of its size and the quantized E^2^M network can be stored using only 1/4 of the space needed for its unquantized counterpart.

## 5 Discussion

We start with a discussion that highlights the reason behind the variations in the performance of various prediction methods for different genes and cancer types. We then proceed to interpret the sources of variation in a biological context.

The results previously presented in Table 5 reveal that the prediction accuracy of E^2^M varies widely for fixed genes and different cancer types. For example, the prediction accuracy for the methylation pattern of gene GATA6 in LUSC and STAD cancers is 0.99 and 0.96, respectively. On the other hand, for the same gene, the prediction accuracy for cancer types such as GBM and LUAD is significantly lower, around 0.6. To gain more insight as of why these variations in accuracy prediction arise, we plot the histograms for different methylation patterns of gene GATA6 in Figure 5. For the GATA6 we observe only one dominant methylation pattern in LUSC and STAD. As a consequence, it is unsurprising that the prediction accuracy of the methylation pattern for cancer types LUSC and STAD is close to one for all methods tested and presented in Table 4.

Interestingly, for the same gene GATA6 we observe 3 and 2 different methylation patterns in GBM and LUAD, respectively. The most likely methylation pattern for GBM has a frequency of about 40%, and this matches the performance of the logistic regression and the fully connected network methods. Indeed, a quick check of the results reveals that the two aforementioned methods almost always predict the dominant methylation pattern. On the other hand, E^2^M is able to capture and predict some of the additional, non-dominant patterns, which is one of the reasons behind its significant performance improvement.

Similar results may be observed in Table 5 for other genes and methylation patterns. The blue and italic entries correspond to cases for which there is a unique dominant methylation pattern in the data, and hence the prediction accuracy of E^2^M is high. The significantly more interesting results are listed in red and bold font as they correspond to settings in which there is more than one dominant methylation pattern, and E^2^M is able to capture at least one more pattern than the other investigated methods. The histograms for all other cancer types and genes considered in the study may be found in Section 3 of the Supplementary Material.

The previous discussion reveals that for different combinations of genes and cancer types one either observes a single dominant or multiple methylation patterns (as many as 12, for the case of gene MGMT and all cancers considered). Let us turn our attention back to Table 5. For example, the promoter regions of gene TP53 and ATM exhibit one dominant methylation pattern (0, unmethylated) across all considered cancer types, while the observed Landmark gene expression profiles differ significantly. This suggests that methylation in TP53 and ATM is most often not the cause of characteristic changes in expression values, and that other regulatory phenomena and point and copy-number mutations may be at work. On the other hand, the promoter regions of genes MLH1, MGMT, CASP8 and KRAS exhibits multiple methylation patterns across all cancer types, with no clear dominant pattern; and, the methylation patterns in MLH1, GATA6 and MGMT associate strongly with the corresponding Landmark gene expressions.

For a more in-depth explanation of these events, we consider BRAIN cancer and gene GATA6 as an illustrative example. Figure 7 shows the heatmap of the expression data of the Landmark genes (left column), as well as the top 15 varying genes (right column), across four types of methylation patterns found in the promoter region of gene GATA6. The horizontal lines in each plot separates the different methylation patterns, sorted by their decimal representation, and the color of each grid represents the magnitude of FPKM readings of the corresponding gene. The right column reveals that the expression level of genes ALDOC, GAPDH, SPP1, APOE, and HLA-DRA change jointly in response to different methylation patterns.

**Fig. 7:**
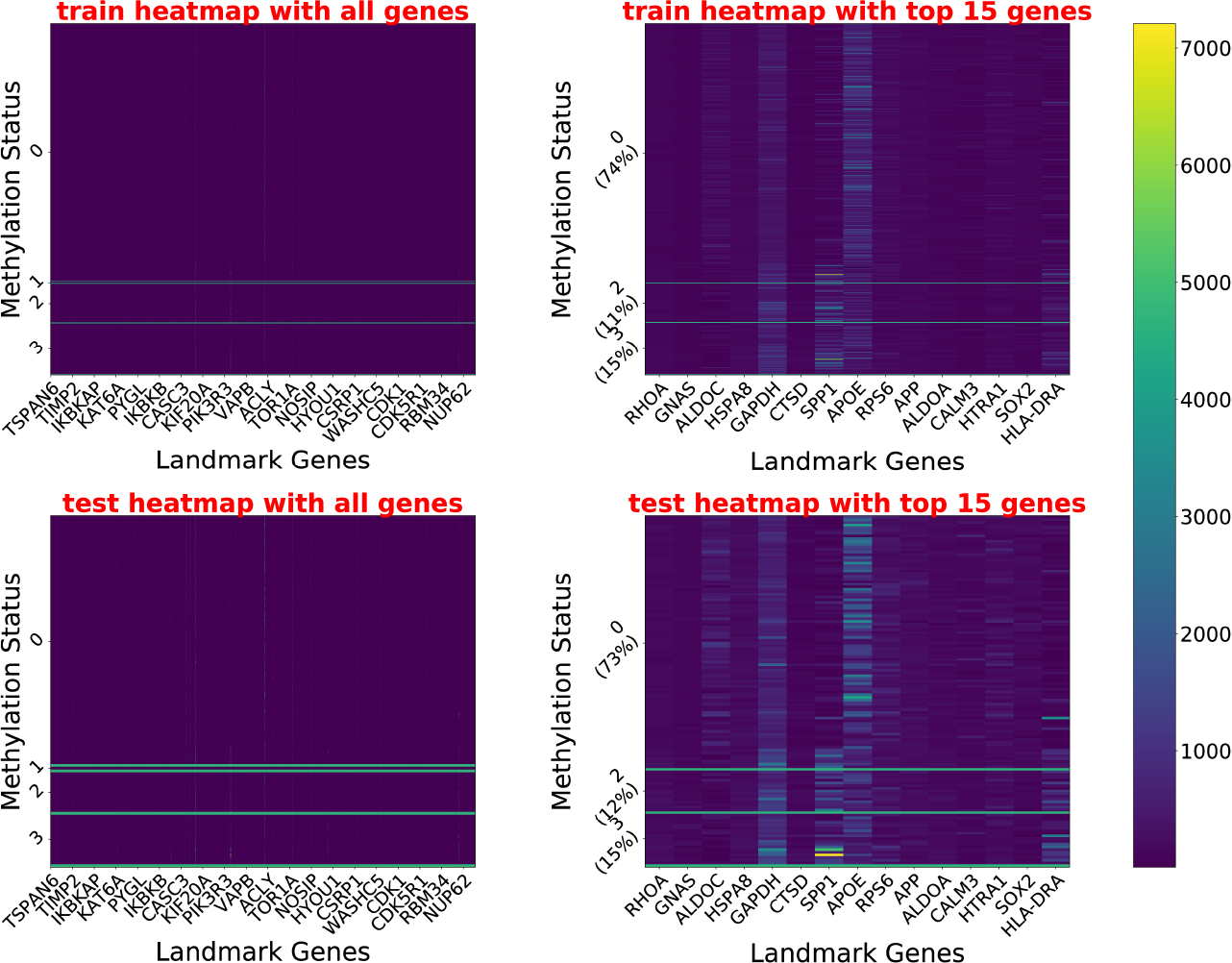
Heatmap of cancer type BRAIN and gene GATA6.

To test if the interactions amongst these genes are due to chance, we performed an enrichment analysis for the selected genes using the pathway datasets KEGG, Biocarta, GeneSigDB, and Reactome. For each pathway, we performed a Fisher exact test with a Null hypothesis assuming mutual independence of the gene variables in the query set. We computed the *p*-values after multiple testing correction, and only selected those with a False Discovery Rate (FDR) less than 0.05. The pathways related to BRAIN cancer (identified through rejection of Null) are the KEGG ALZHEIMERS DISEASE (from KEGG), the REACTOME GLUCOSE METABOLISM, the METABOLISM OF CARBOHYDRATES (from Reactome) and the Mouse Brain Johansson and genes UpRegulatedbyHypoxia (from GeneSigDB). It is known from previously reported studies that these pathways are indeed involved in the progression of brain cancer [28, 16, 26, 27].

In summary, our most important biological finding is that *patterns of methylation sites*, rather than the global methylation status of a gene (methylated or unmethylated) alone, govern Landmark and global gene expressions. This observation is strongly supported by the good predictive performance of E^2^M on the CASP8 gene for LUAD and STAD, and the MLH1 gene for STAD. In both cases, at least two patterns which are *both* deemed globally methylated can be accurately distinguished from each other thought their expression profile.

## 6 Conclusion

We proposed an inception based deep learning framework, termed E^2^M, capable of associating specific methylation patterns in gene promoter regions with Landmark and consequently global gene expression. We tested the proposed framework on TCGA data including five cancer types, and the promoter regions of seven genes. Our findings were two-fold. First, we showed that the proposed E^2^M frame-work outperforms multiclass logistics regression and 3-layer fully connected network in prediction accuracy. Furthermore, the performance of E^2^M was shown not to be affected by quantization of both the input data and the weights of the inception network. Second, we found that methylation of some tumor suppressor genes does not bear a detectable influence on the expression profiles; at the same time, different methylation patterns in the promoter regions of the same gene can lead to observable changes in the gene expressions, even when the patterns result in the same binary methylation status.

As a final remark, we point out that E^2^M is a general learning framework that may be successfully applied to other multiomics data association studies and single cell researches [2].

## Acknowledgements

The authors are grateful to Mikel Hernaez from the University of Illinois for many useful discussions and for providing the gene enrichment analysis results.

