## Supplementary for "E^2^M: A Deep Learning Framework for Associating Combinatorial Methylation Patterns with Gene Expression"

### Supplementary Material
### Predicting Combinatorial Methylation Patterns Based on Gene Expression via E$^2$M Deep Learning

Jianhao Peng, Olgica Milenkovic and Idoia Ochoa

Department of Electrical and Computer Engineering,
University of Illinois at Urbana Champaign

The Supplementary Material provides more comprehensive findings for both the comparative performance of E$^2$M and regarding the biological significance of methylation patterns. In particular, we report results for all chosen genes (MGMT, MLH1, ATM, GATA6, KRAS, CASP8 and TP53) and cancer types, including results for MR (Multiclass Regression), FCNN (Fully Connected Neural Network) and E$^2$M, PCA results, histograms of methylation patterns and expression heatmaps for all datasets.

### 1 Gene Description

We selected 7 tumor suppressor/repair genes for our study of methylation pattern prediction based on their relevance in tumorigenesis. The genes listed below, along with a description of their function.

- **MGMT**: **repair gene**. MGMT gene produces the O6-alkylguanine DNA alkyltransferase. It repairs the naturally occurring mutagenic DNA lesion O6-methylguanine back to guanine and prevents mismatch and errors during DNA replication and transcription [2].
- **MLH1**: **tumor suppressor**. Its product, MutL homolog 1, is a DNA mismatch repair protein. According to previous studies [3, 5], methylation of MLH1 inhibits expression and is commonly mutated or methylated in stomach and lung cancers.
- **ATM**: **repair gene**. One of ATM's protein product is involved in controlling cell division. It also aids other genes in finding and repairing damaged DNA.
- **GATA6**: **tumor suppressor**. It produces one of the GATA Transcription factor, GATA-6, which binds (A/T/C)GAT(A/T)(A) of the consensus binding sequence. Such transcription factors play a crucial role in early cell differentiation and development. The gene is strongly expressed in many metastatic cancers [4].
- **CASP8**: **tumor suppressor**. CASP8 is in the cysteine-aspartic acid protease (caspase) family, and plays an important role in controlling cell apoptosis. It is involved in different cancer types and a biomarker for cancer [6].
- **KRAS**: **tumor suppressor**. The disruption of gene KRAS causes continuous cell proliferation and is an early cancer driver gene. It is also a proto-oncogene and was observed to have heightened activity in many malignancies, including lung adenocarcinoma (LUAD) [1].
- **TP53**: **tumor suppressor**. TP53 produces the tumor protein p53. It also controls cell proliferation.

### 2 PCA Results

The presented results include PCA visualizations of expressions of all genes under consideration, for all cancer types.

#### 2.1 GBM

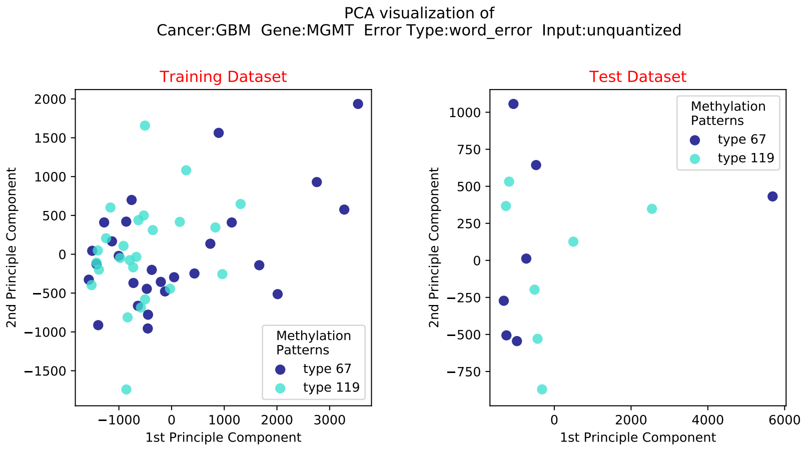
PCA visualization of
Cancer:GBM Gene:MGMT Error Type:word_error Input:unquantized
Training Dataset
Test Dataset
1st Principle Component
2nd Principle Component
Methylation
Patterns
type 67
type 119

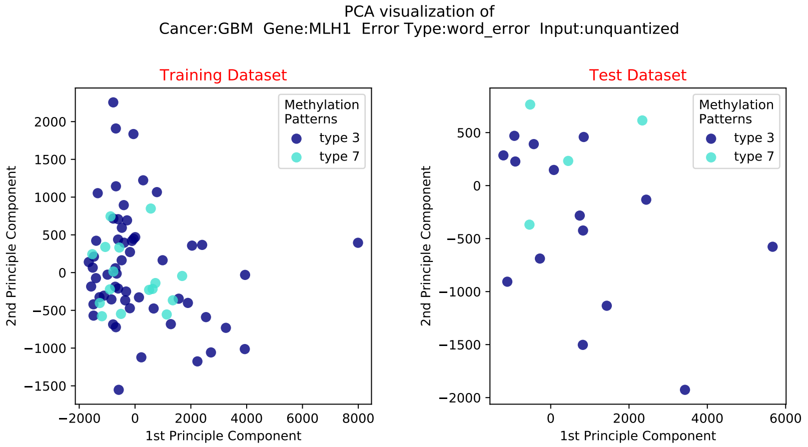
PCA visualization of
Cancer:GBM Gene:MLH1 Error Type:word_error Input:unquantized
Training Dataset
Test Dataset
1st Principle Component
2nd Principle Component
Methylation
Patterns
type 3
type 7

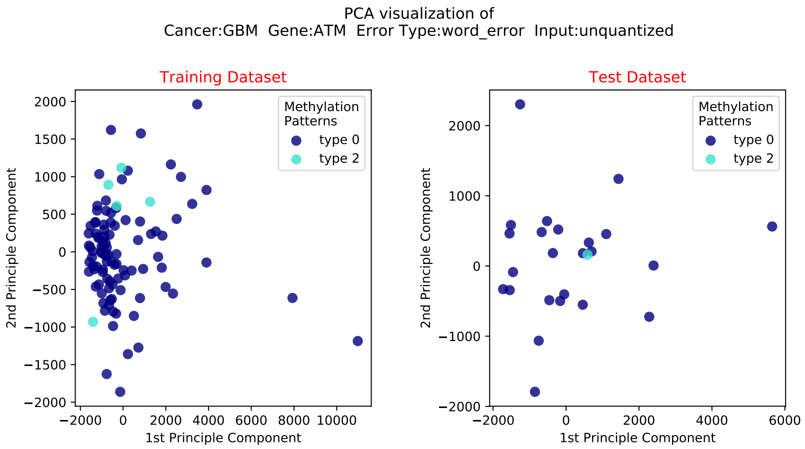
PCA visualization of
Cancer:GBM Gene:ATM Error Type:word_error Input:unquantized
Training Dataset
Test Dataset
Methylation Patterns
type 0
type 2
2nd Principle Component
1st Principle Component

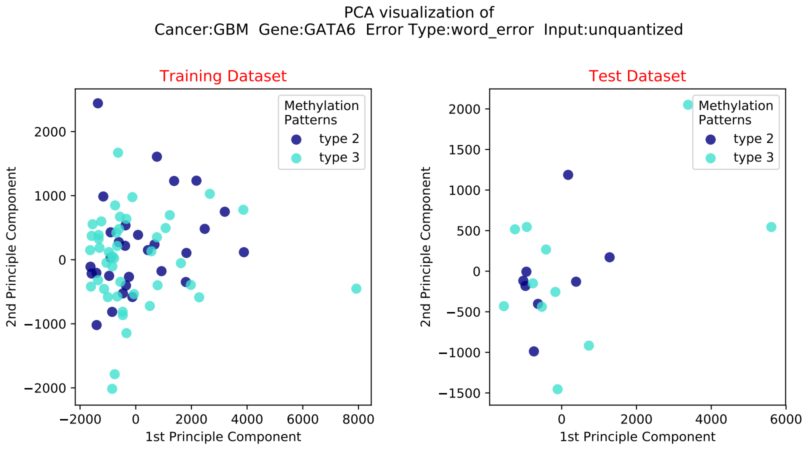
PCA visualization of
Cancer:GBM Gene:GATA6 Error Type:word_error Input:unquantized
Training Dataset
Test Dataset
Methylation Patterns
type 2
type 3
2nd Principle Component
1st Principle Component

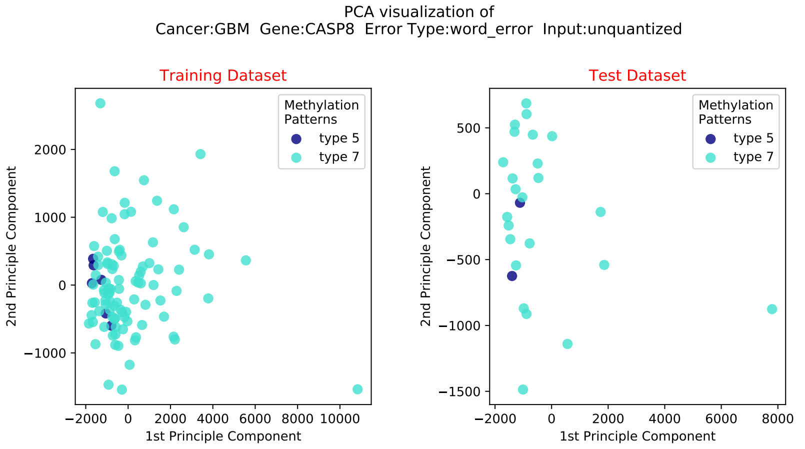
PCA visualization of
Cancer:GBM Gene:CASP8 Error Type:word_error Input:unquantized
Training Dataset
Test Dataset
Methylation Patterns
type 5
type 7
2nd Principle Component
1st Principle Component

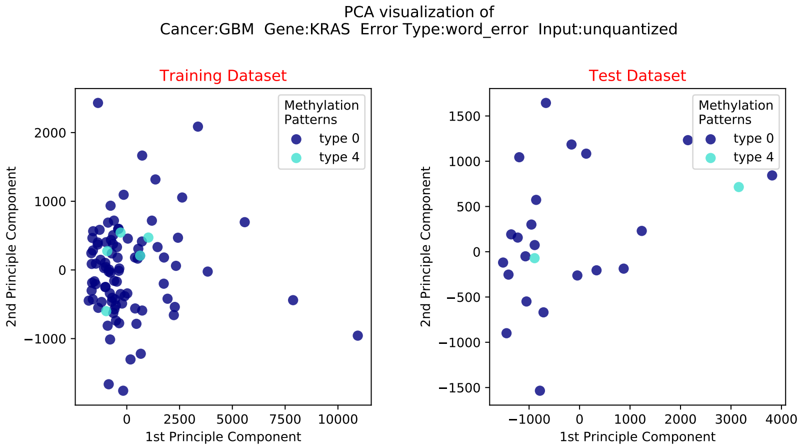
PCA visualization of
Cancer:GBM Gene:KRAS Error Type:word_error Input:unquantized
Training Dataset
Test Dataset
Methylation Patterns
type 0
type 4
2nd Principle Component
1st Principle Component

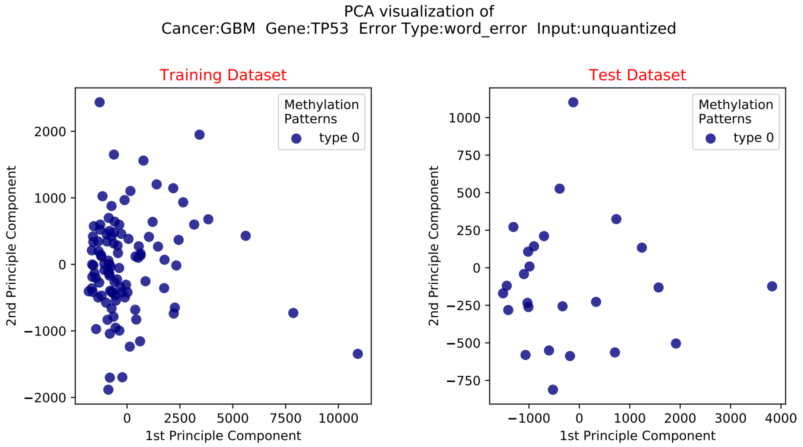
PCA visualization of
Cancer:GBM Gene:TP53 Error Type:word_error Input:unquantized
Training Dataset
Test Dataset
Methylation Patterns
type 0
2nd Principle Component
1st Principle Component

#### 2.2 LGG

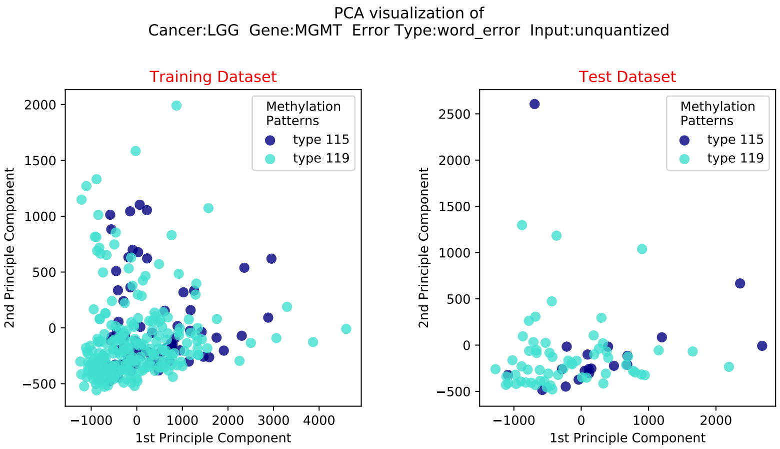
PCA visualization of
Cancer:LGG Gene:MGMT Error Type:word_error Input:unquantized
Training Dataset
Test Dataset
Methylation Patterns
type 115
type 119
1st Principle Component
2nd Principle Component

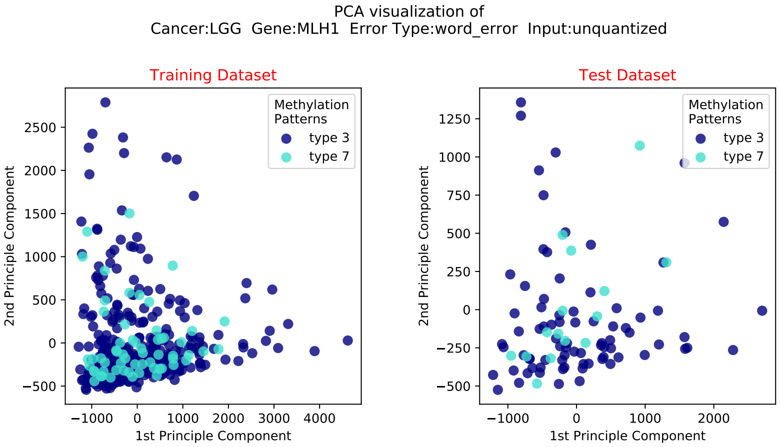
PCA visualization of
Cancer:LGG Gene:MLH1 Error Type:word_error Input:unquantized
Training Dataset
Test Dataset
Methylation Patterns
type 3
type 7
1st Principle Component
2nd Principle Component

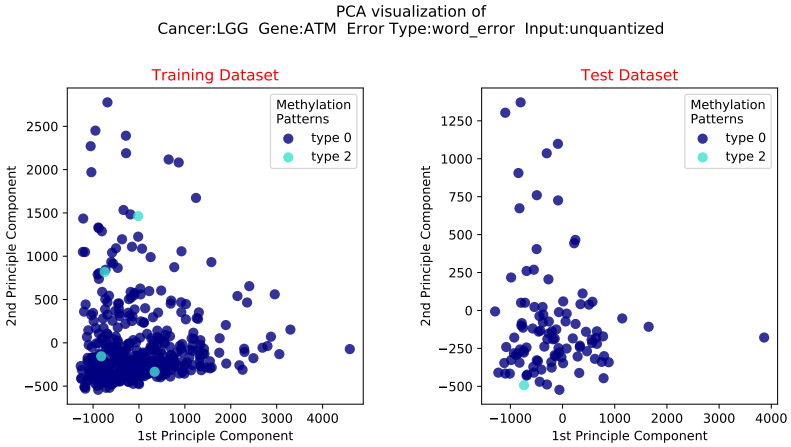
PCA visualization of
Cancer:LGG Gene:ATM Error Type:word_error Input:unquantized
Training Dataset
Test Dataset
Methylation Patterns
type 0
type 2
2nd Principle Component
1st Principle Component

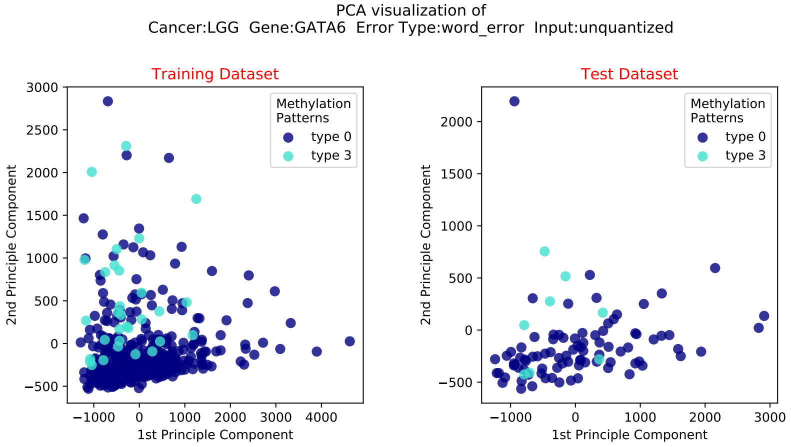
PCA visualization of
Cancer:LGG Gene:GATA6 Error Type:word_error Input:unquantized
Training Dataset
Test Dataset
Methylation Patterns
type 0
type 3
2nd Principle Component
1st Principle Component

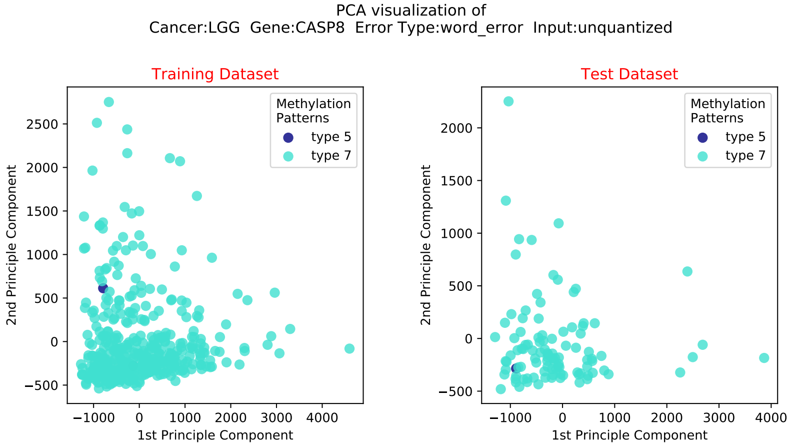
PCA visualization of
Cancer:LGG Gene:CASP8 Error Type:word_error Input:unquantized
Training Dataset
Test Dataset
Methylation Patterns
type 5
type 7
2nd Principle Component
1st Principle Component

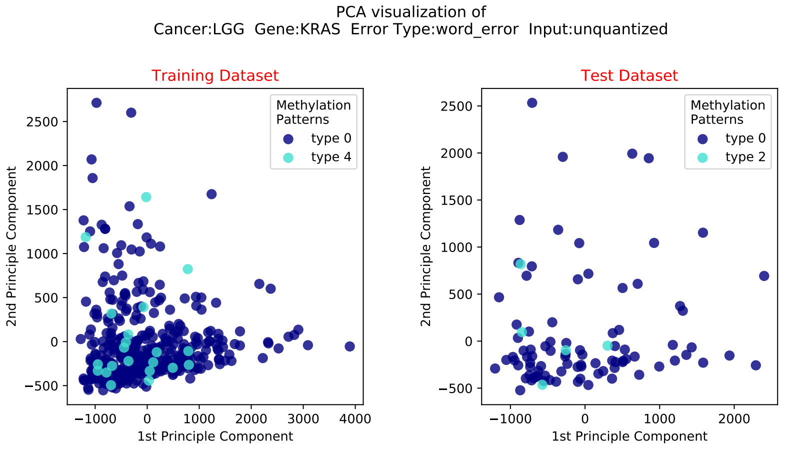
PCA visualization of
Cancer:LGG Gene:KRAS Error Type:word_error Input:unquantized
Training Dataset
Test Dataset
Methylation Patterns
type 0
type 4
type 2
2nd Principle Component
1st Principle Component

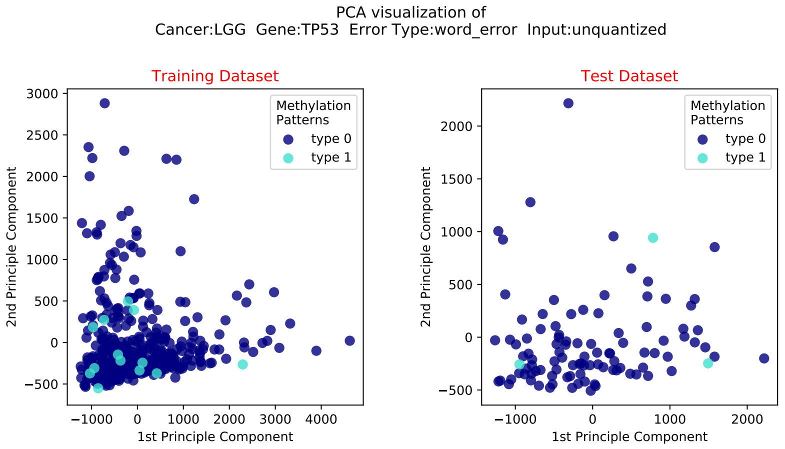
PCA visualization of
Cancer:LGG Gene:TP53 Error Type:word_error Input:unquantized
Training Dataset
Test Dataset
Methylation Patterns
type 0
type 1
2nd Principle Component
1st Principle Component

#### 2.3 Brain Cancer

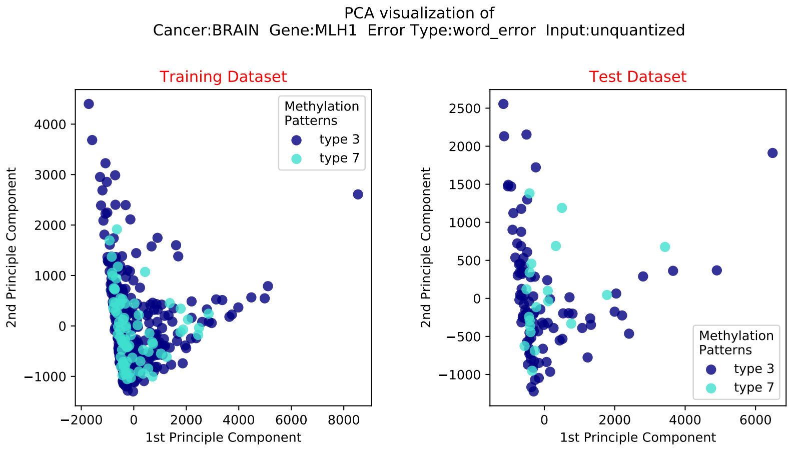

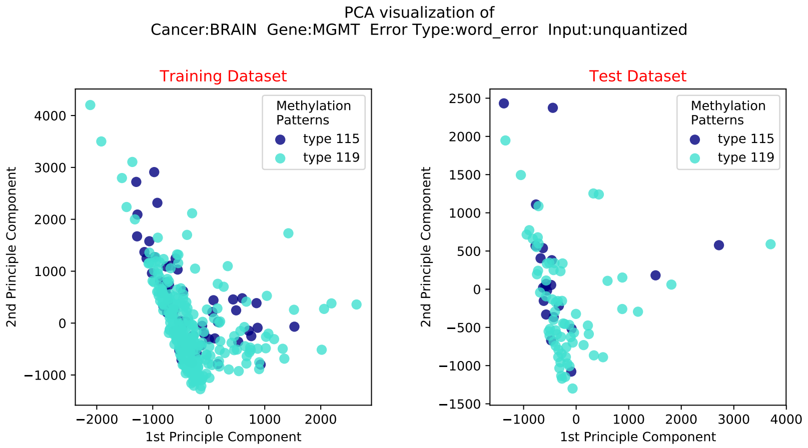

PCA visualization of
Cancer:BRAIN Gene:ATM Error Type:word_error Input:unquantized

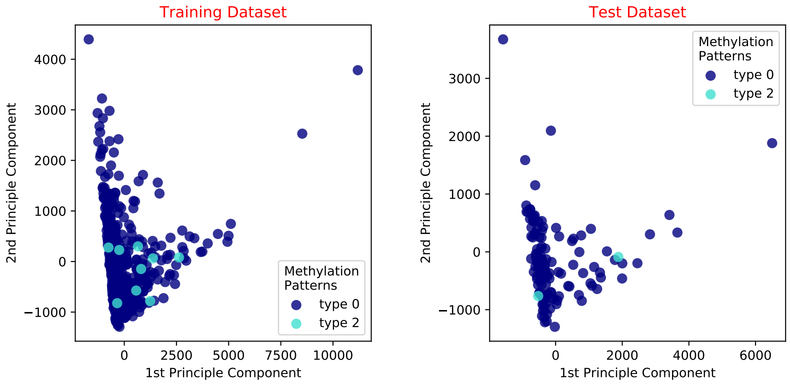

PCA visualization of
Cancer:BRAIN Gene:GATA6 Error Type:word_error Input:unquantized

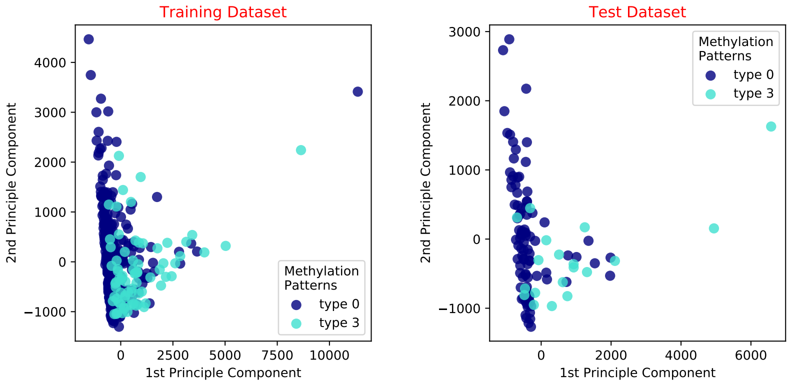

PCA visualization of
Cancer:BRAIN Gene:CASP8 Error Type:word_error Input:unquantized

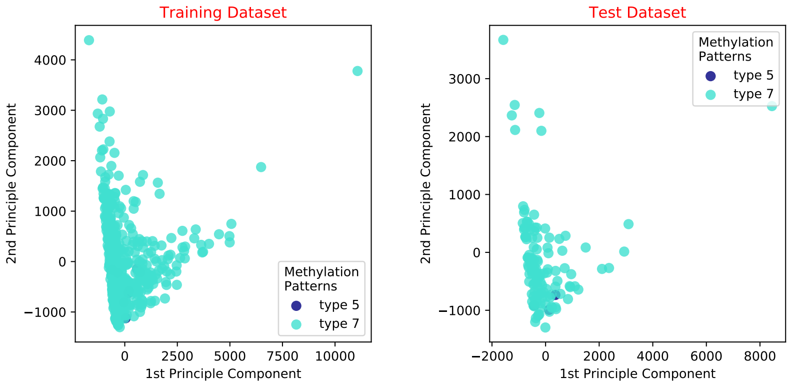

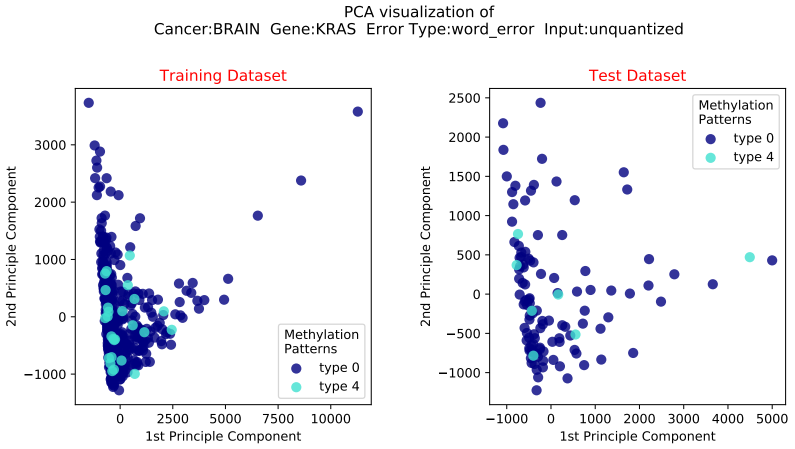
PCA visualization of
Cancer:BRAIN Gene:KRAS Error Type:word_error Input:unquantized
Training Dataset
Test Dataset
2nd Principle Component
1st Principle Component
Methylation Patterns
type 0
type 4

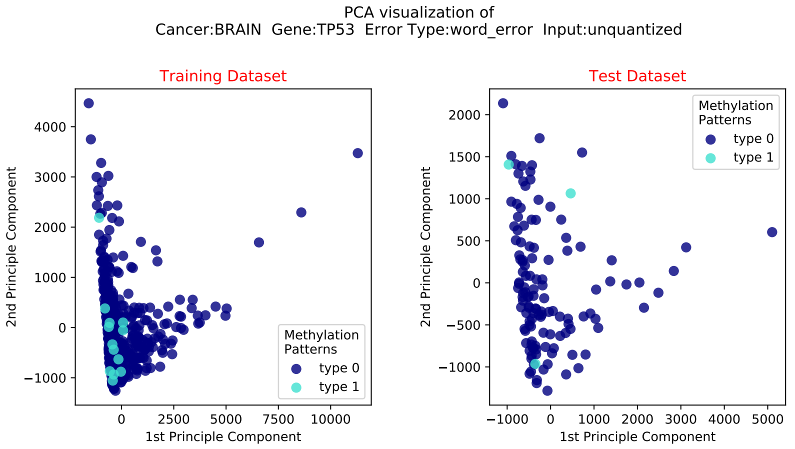
PCA visualization of
Cancer:BRAIN Gene:TP53 Error Type:word_error Input:unquantized
Training Dataset
Test Dataset
2nd Principle Component
1st Principle Component
Methylation Patterns
type 0
type 1

#### 2.4 LUAD

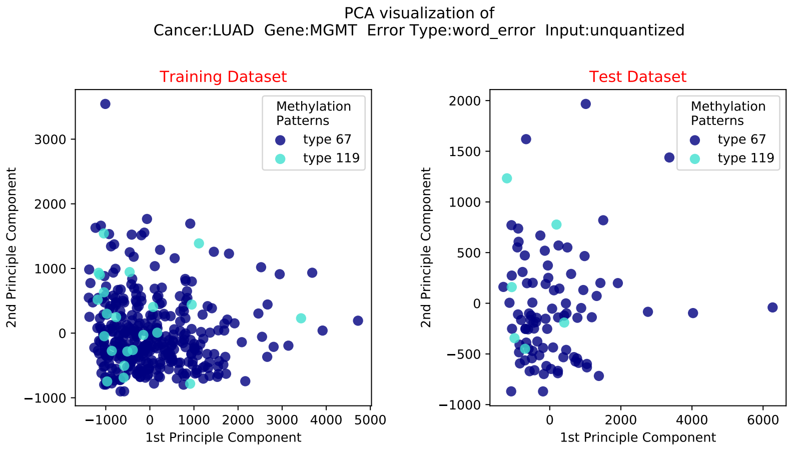
PCA visualization of
Cancer:LUAD Gene:MGMT Error Type:word_error Input:unquantized
Training Dataset
Test Dataset
Methylation Patterns
type 67
type 119
2nd Principle Component
1st Principle Component

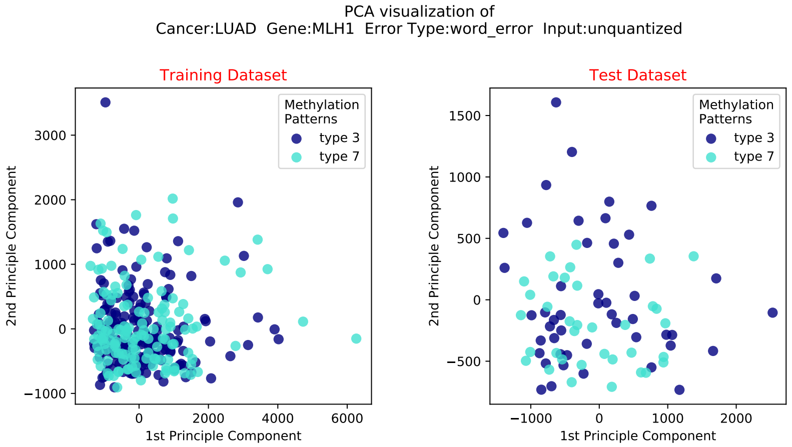
PCA visualization of
Cancer:LUAD Gene:MLH1 Error Type:word_error Input:unquantized
Training Dataset
Test Dataset
Methylation Patterns
type 3
type 7
2nd Principle Component
1st Principle Component

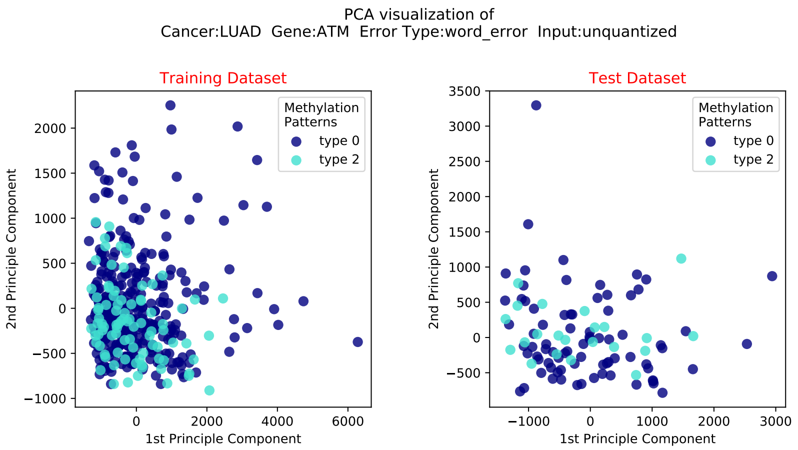

PCA visualization of
Cancer:LUAD Gene:ATM Error Type:word_error Input:unquantized
Training Dataset
Test Dataset
Methylation Patterns
type 0
type 2
2nd Principle Component
1st Principle Component

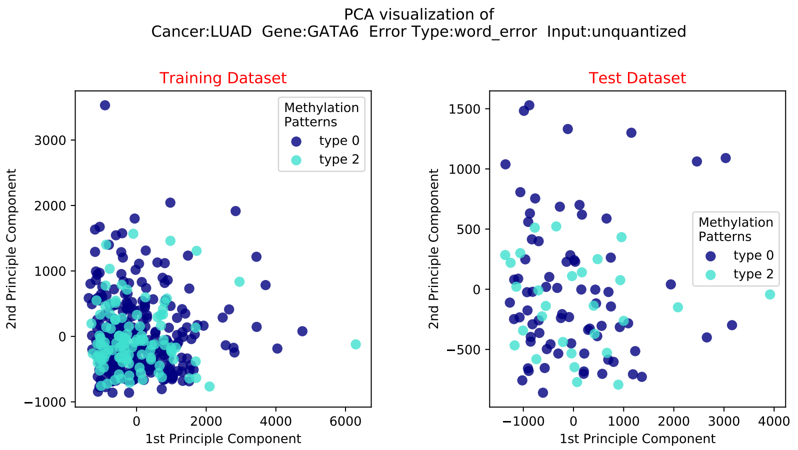

PCA visualization of
Cancer:LUAD Gene:GATA6 Error Type:word_error Input:unquantized
Training Dataset
Test Dataset
Methylation Patterns
type 0
type 2
2nd Principle Component
1st Principle Component

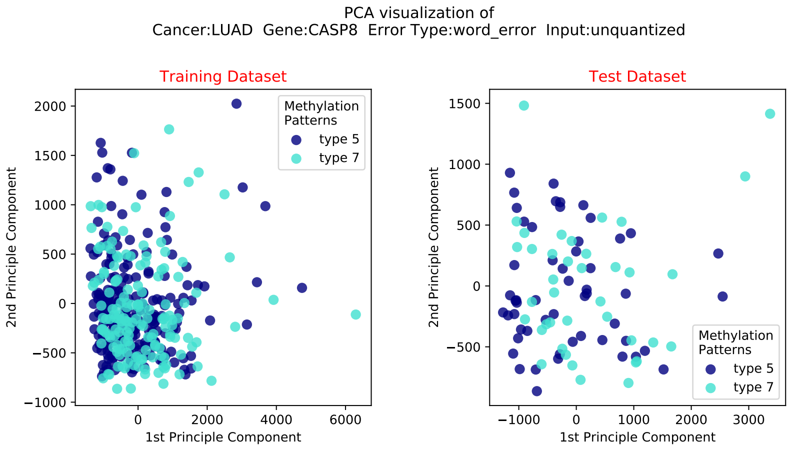

PCA visualization of
Cancer:LUAD Gene:CASP8 Error Type:word_error Input:unquantized
Training Dataset
Test Dataset
Methylation Patterns
type 5
type 7
2nd Principle Component
1st Principle Component

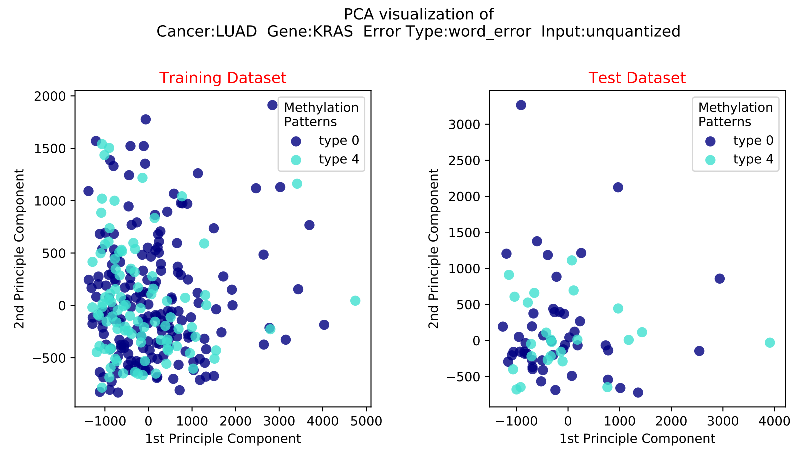
PCA visualization of
Cancer:LUAD Gene:KRAS Error Type:word_error Input:unquantized
Training Dataset
Test Dataset
Methylation Patterns
type 0
type 4
1st Principle Component
2nd Principle Component

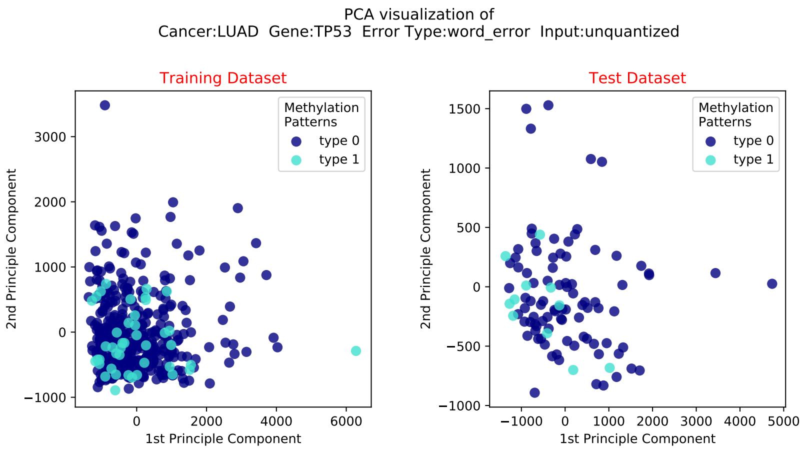
PCA visualization of
Cancer:LUAD Gene:TP53 Error Type:word_error Input:unquantized
Training Dataset
Test Dataset
Methylation Patterns
type 0
type 1
1st Principle Component
2nd Principle Component

#### 2.5 LUSC

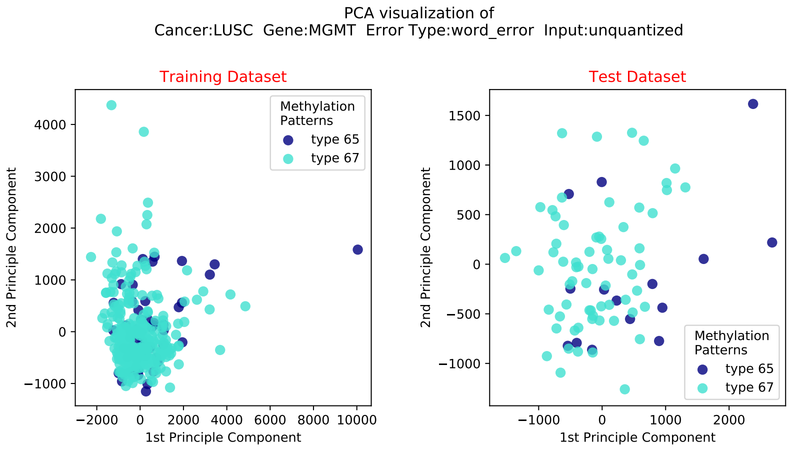
PCA visualization of
Cancer:LUSC Gene:MGMT Error Type:word_error Input:unquantized
Training Dataset
Test Dataset
Methylation Patterns
type 65
type 67
1st Principle Component
2nd Principle Component

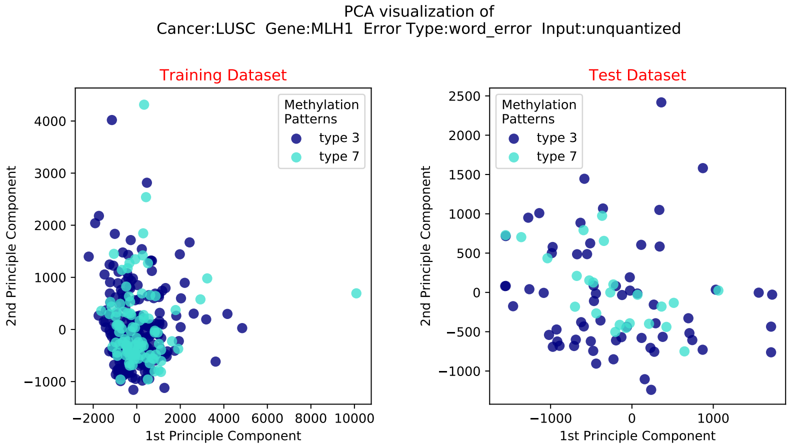
PCA visualization of
Cancer:LUSC Gene:MLH1 Error Type:word_error Input:unquantized
Training Dataset
Test Dataset
Methylation Patterns
type 3
type 7
1st Principle Component
2nd Principle Component

PCA visualization of
Cancer:LUSC  Gene:ATM  Error Type:word_error  Input:unquantized

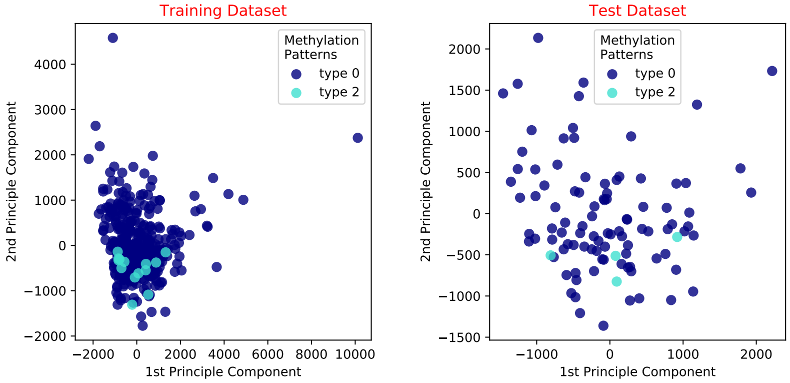

PCA visualization of
Cancer:LUSC  Gene:GATA6  Error Type:word_error  Input:unquantized

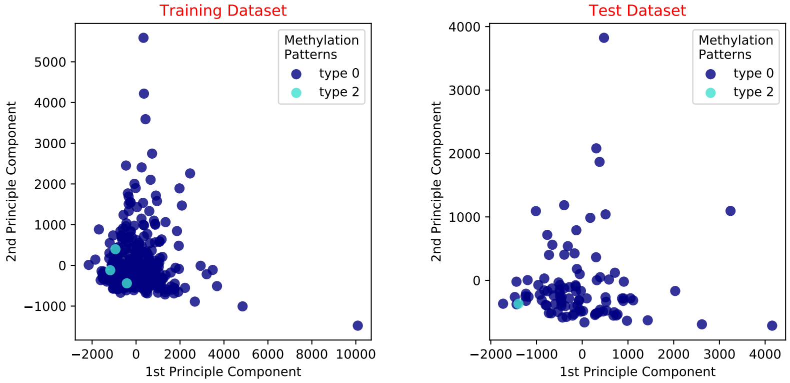

PCA visualization of
Cancer:LUSC  Gene:CASP8  Error Type:word_error  Input:unquantized

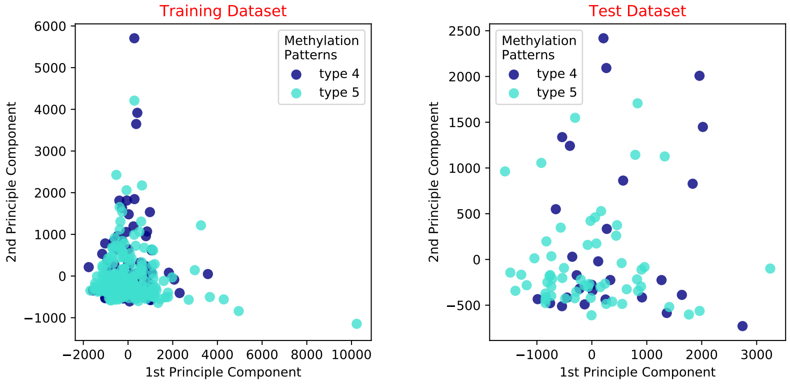

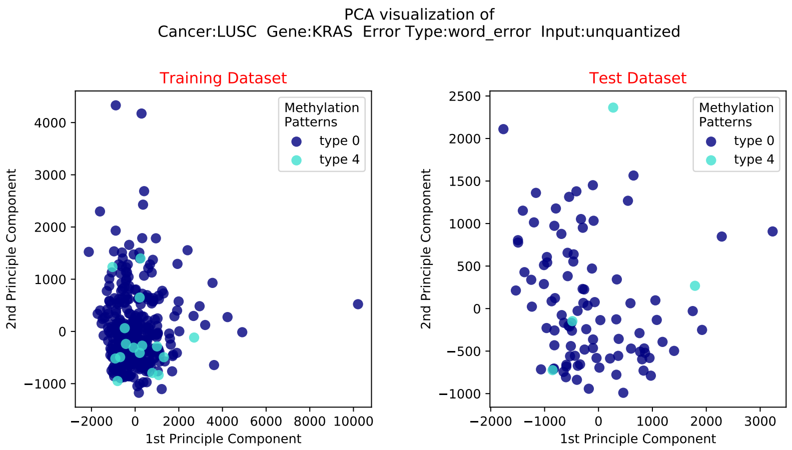
PCA visualization of
Cancer:LUSC Gene:KRAS Error Type:word_error Input:unquantized
Training Dataset
Test Dataset
Methylation Patterns
type 0
type 4
1st Principle Component
2nd Principle Component

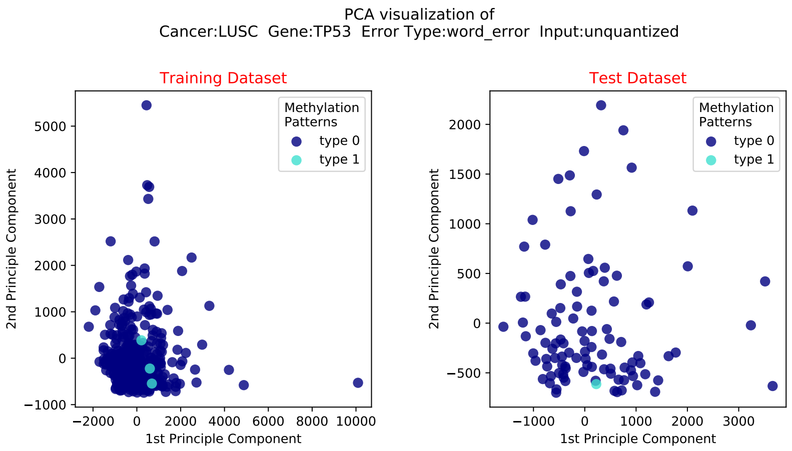
PCA visualization of
Cancer:LUSC Gene:TP53 Error Type:word_error Input:unquantized
Training Dataset
Test Dataset
Methylation Patterns
type 0
type 1
1st Principle Component
2nd Principle Component

#### 2.6 Lung Cancer

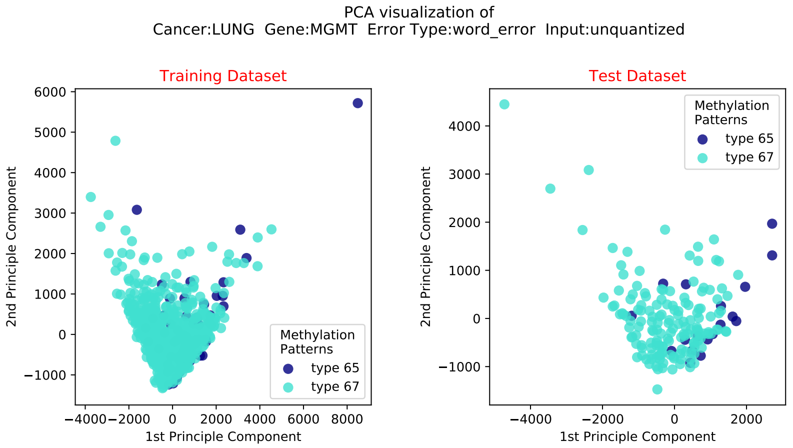

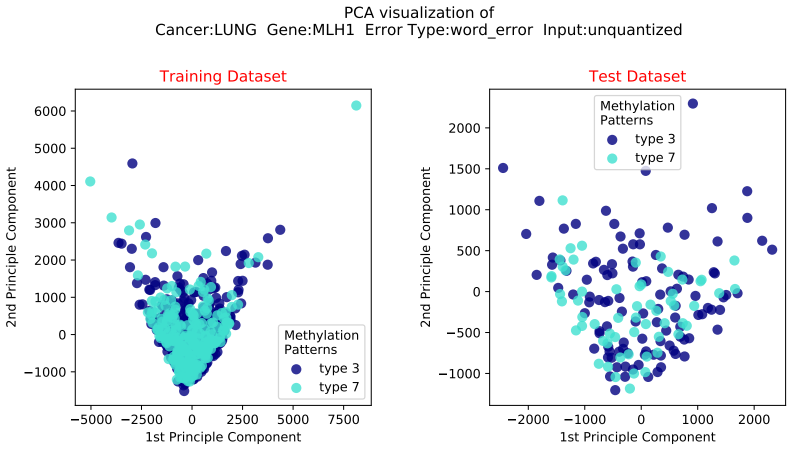

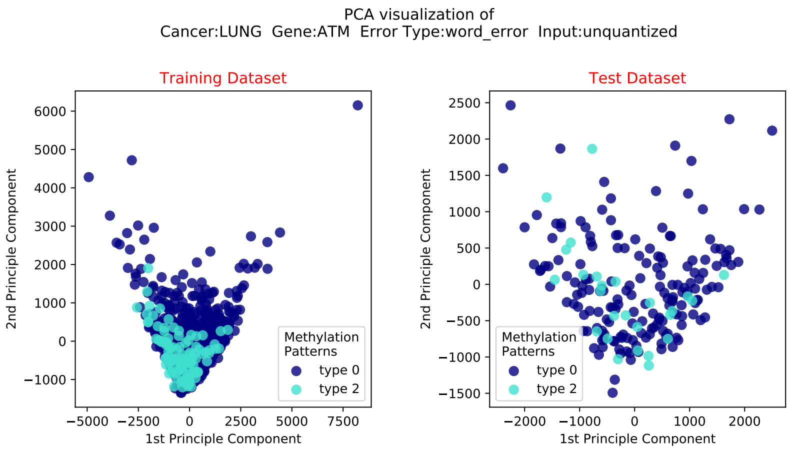
PCA visualization of
Cancer:LUNG Gene:ATM Error Type:word_error Input:unquantized
Training Dataset
Test Dataset
2nd Principle Component
1st Principle Component
Methylation Patterns
type 0
type 2

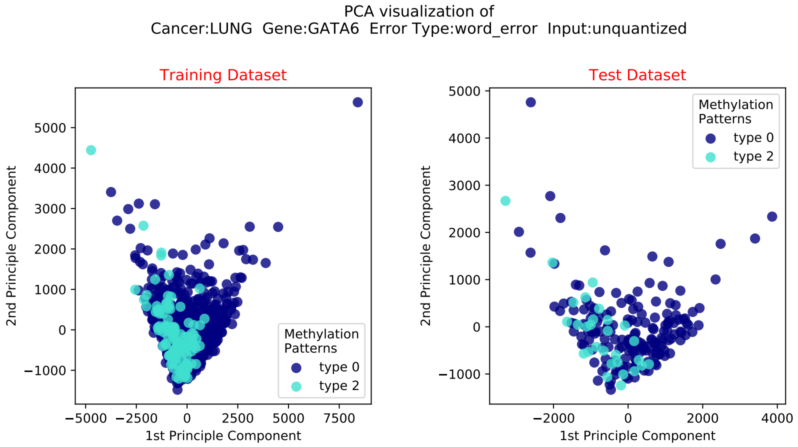
PCA visualization of
Cancer:LUNG Gene:GATA6 Error Type:word_error Input:unquantized
Training Dataset
Test Dataset
2nd Principle Component
1st Principle Component
Methylation Patterns
type 0
type 2

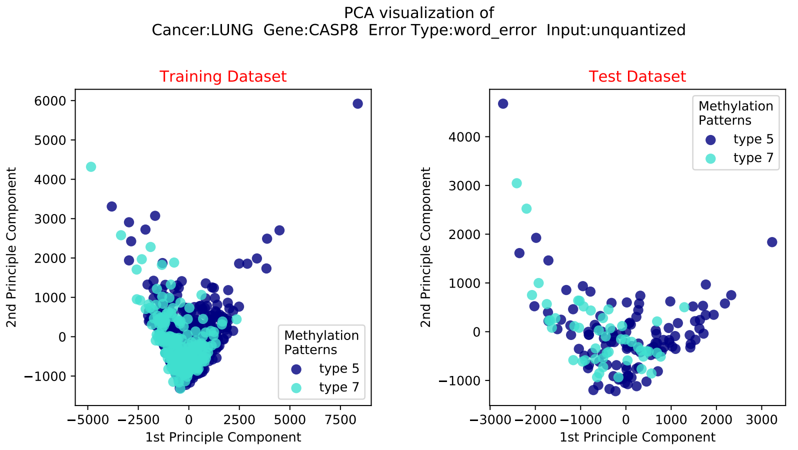
PCA visualization of
Cancer:LUNG Gene:CASP8 Error Type:word_error Input:unquantized
Training Dataset
Test Dataset
2nd Principle Component
1st Principle Component
Methylation Patterns
type 5
type 7

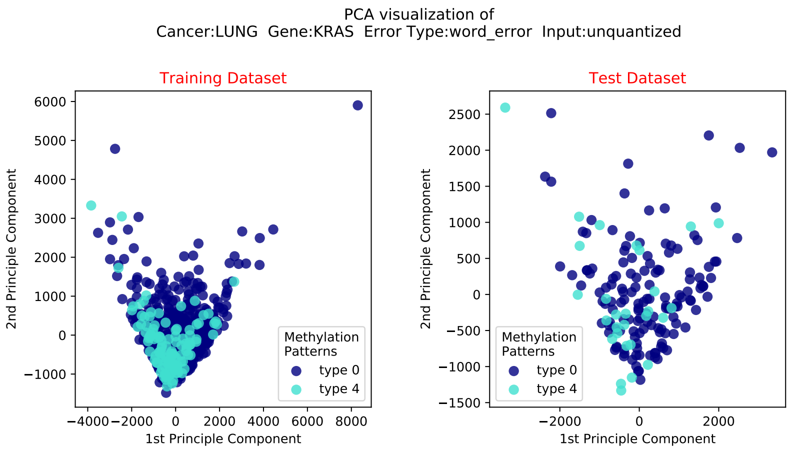
PCA visualization of
Cancer:LUNG Gene:KRAS Error Type:word_error Input:unquantized
Training Dataset
Test Dataset
2nd Principle Component
1st Principle Component
Methylation Patterns
type 0
type 4

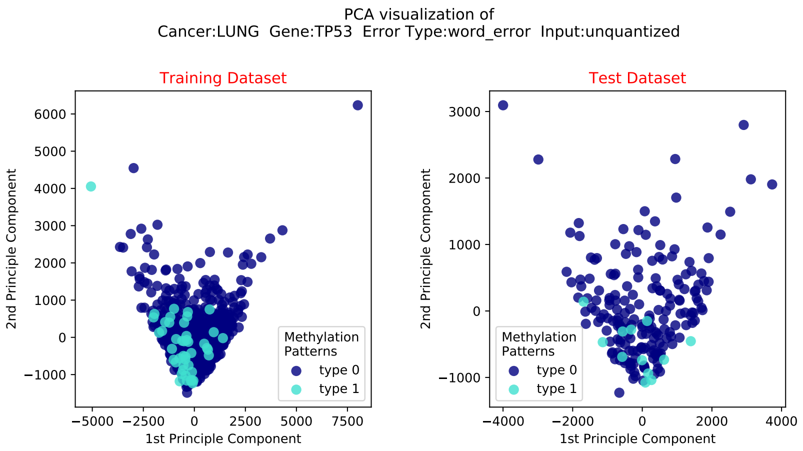
PCA visualization of
Cancer:LUNG Gene:TP53 Error Type:word_error Input:unquantized
Training Dataset
Test Dataset
2nd Principle Component
1st Principle Component
Methylation Patterns
type 0
type 1

#### 2.7 STAD

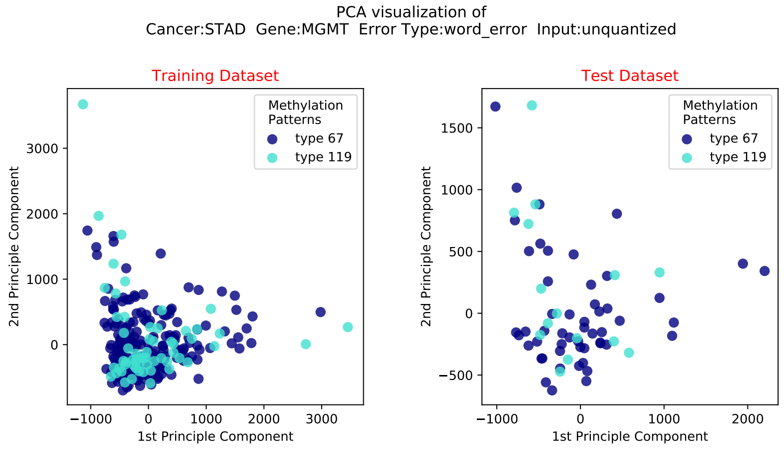
PCA visualization of
Cancer:STAD Gene:MGMT Error Type:word_error Input:unquantized
Training Dataset
Test Dataset
Methylation Patterns
type 67
type 119
1st Principle Component
2nd Principle Component

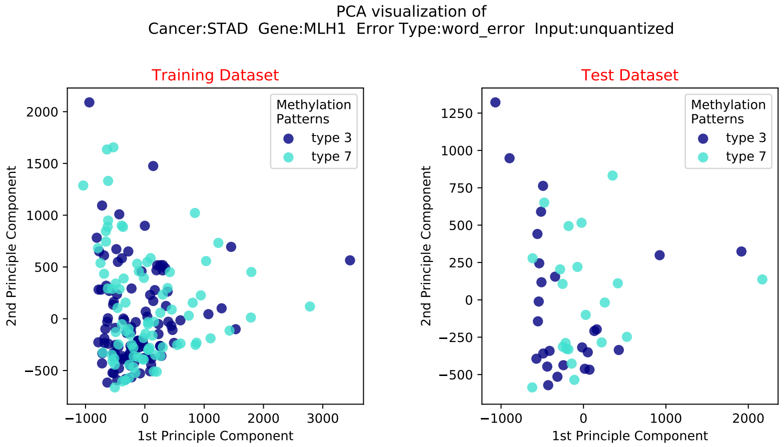
PCA visualization of
Cancer:STAD Gene:MLH1 Error Type:word_error Input:unquantized
Training Dataset
Test Dataset
Methylation Patterns
type 3
type 7
1st Principle Component
2nd Principle Component

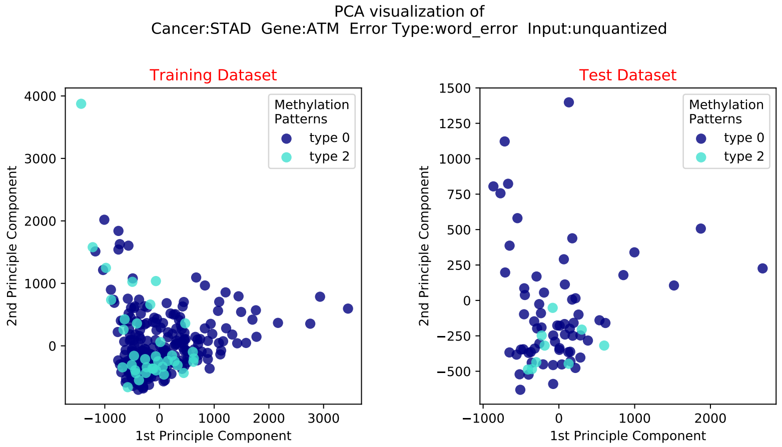
PCA visualization of
Cancer:STAD Gene:ATM Error Type:word_error Input:unquantized
Training Dataset
Test Dataset
Methylation Patterns
type 0
type 2
1st Principle Component
2nd Principle Component

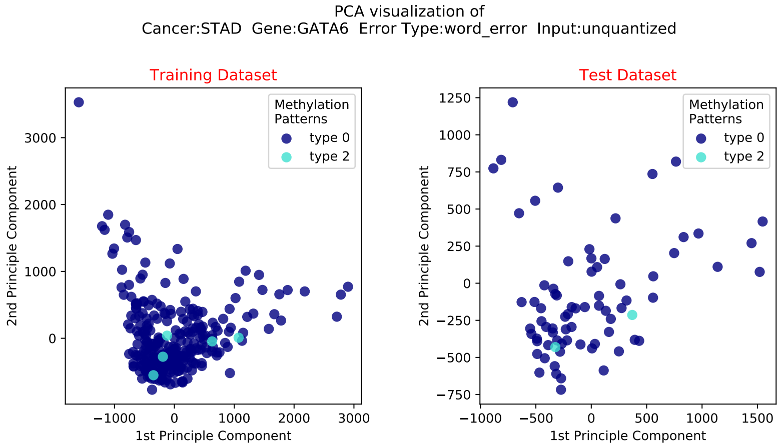
PCA visualization of
Cancer:STAD Gene:GATA6 Error Type:word_error Input:unquantized
Training Dataset
Test Dataset
Methylation Patterns
type 0
type 2
1st Principle Component
2nd Principle Component

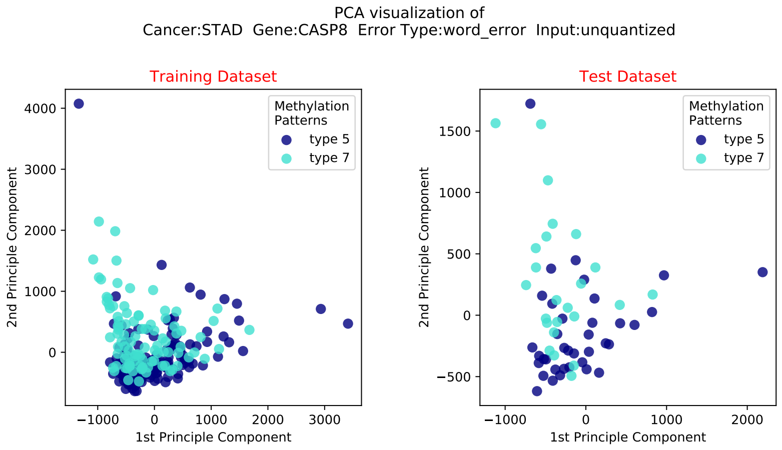
PCA visualization of
Cancer:STAD Gene:CASP8 Error Type:word_error Input:unquantized
Training Dataset
Test Dataset
Methylation Patterns
type 5
type 7
1st Principle Component
2nd Principle Component

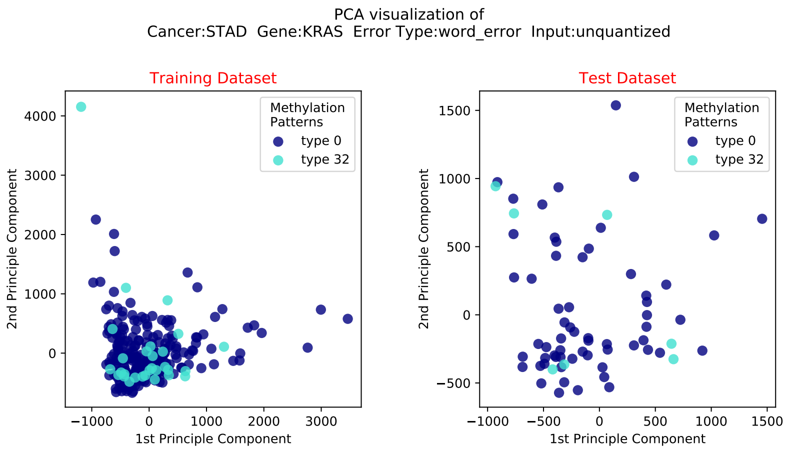
PCA visualization of
Cancer:STAD Gene:KRAS Error Type:word_error Input:unquantized
Training Dataset
Test Dataset
Methylation Patterns
type 0
type 32
1st Principle Component
2nd Principle Component

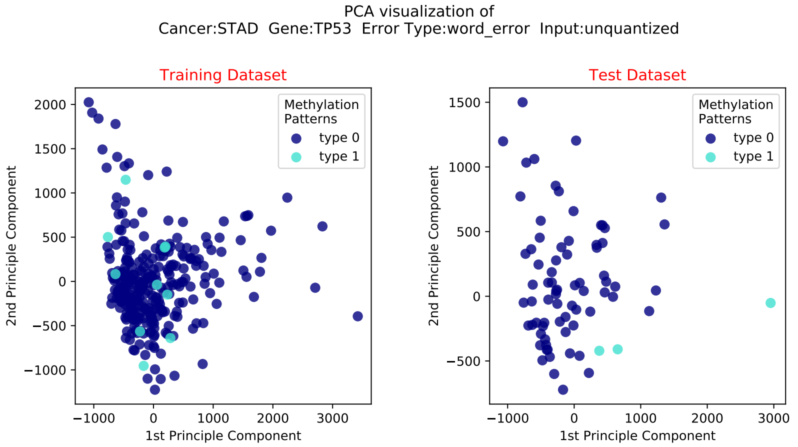
PCA visualization of
Cancer:STAD Gene:TP53 Error Type:word_error Input:unquantized
Training Dataset
Test Dataset
Methylation Patterns
type 0
type 1
1st Principle Component
2nd Principle Component

### 3 Histogram of Methylation Pattern

The presented results include histograms of methylation patterns for all genes under consideration, for all cancer types.

#### 3.1 GBM

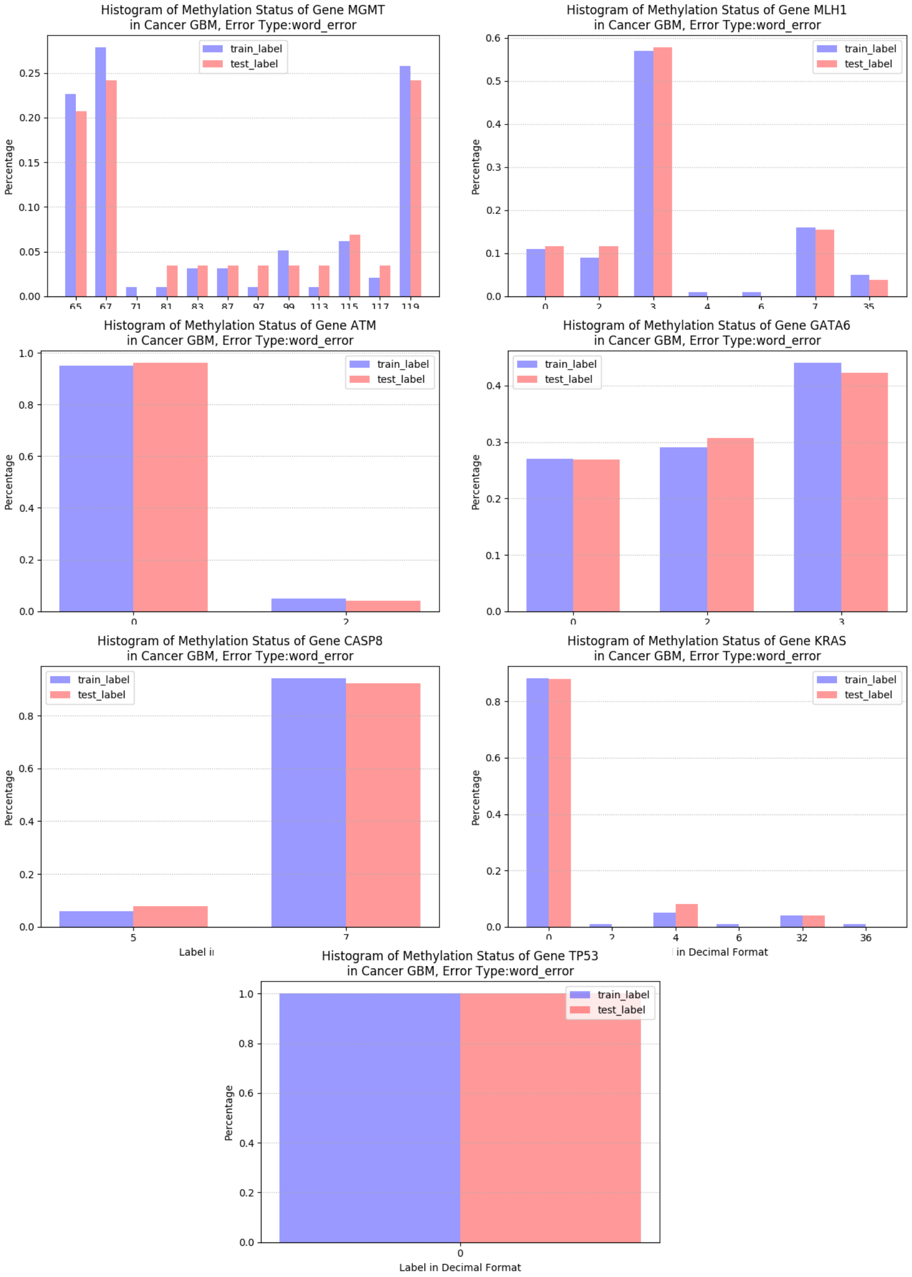

#### 3.2 LGG

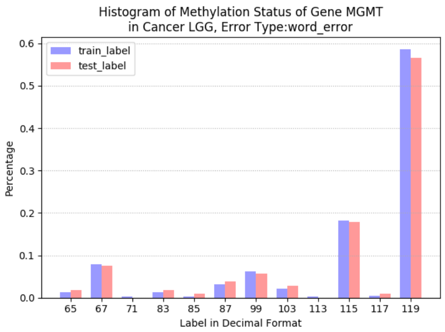
Histogram of Methylation Status of Gene MGMT
in Cancer LGG, Error Type:word_error
train_label
test_label
Percentage
0.0 0.1 0.2 0.3 0.4 0.5 0.6
65 67 71 83 85 87 99 103 113 115 117 119
Label in Decimal Format

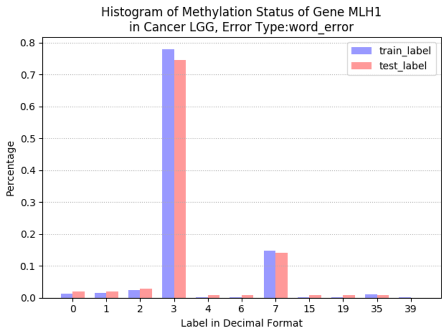
Histogram of Methylation Status of Gene MLH1
in Cancer LGG, Error Type:word_error
train_label
test_label
Percentage
0.0 0.1 0.2 0.3 0.4 0.5 0.6 0.7 0.8
0 1 2 3 4 6 7 15 19 35 39
Label in Decimal Format

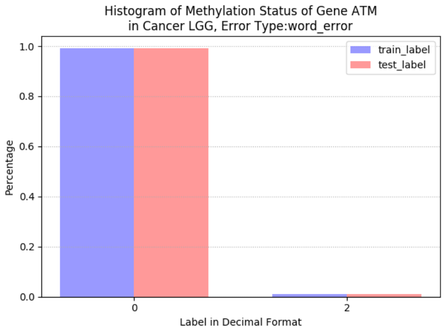
Histogram of Methylation Status of Gene ATM
in Cancer LGG, Error Type:word_error
train_label
test_label
Percentage
0.0 0.2 0.4 0.6 0.8 1.0
0 2
Label in Decimal Format

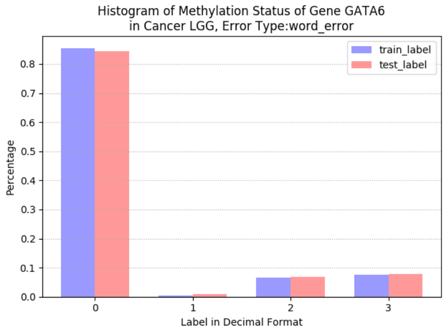
Histogram of Methylation Status of Gene GATA6
in Cancer LGG, Error Type:word_error
train_label
test_label
Percentage
0.0 0.1 0.2 0.3 0.4 0.5 0.6 0.7 0.8
0 1 2 3
Label in Decimal Format

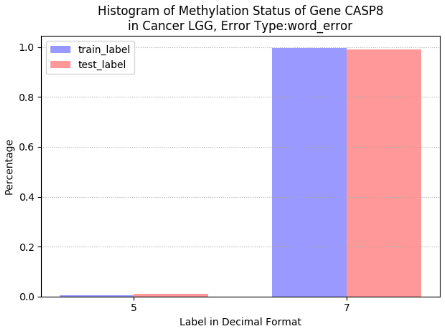
Histogram of Methylation Status of Gene CASP8
in Cancer LGG, Error Type:word_error
train_label
test_label
Percentage
0.0 0.2 0.4 0.6 0.8 1.0
5 7
Label in Decimal Format

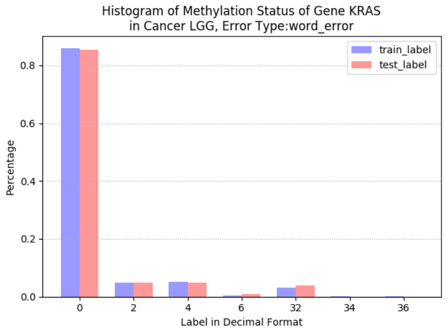
Histogram of Methylation Status of Gene KRAS
in Cancer LGG, Error Type:word_error
train_label
test_label
Percentage
0.0 0.2 0.4 0.6 0.8
0 2 4 6 32 34 36
Label in Decimal Format

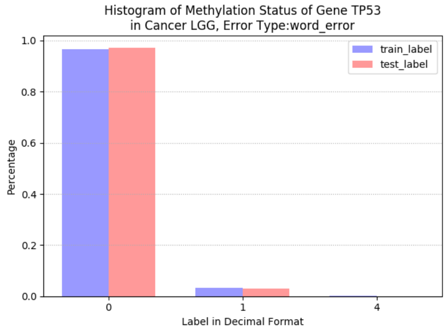
Histogram of Methylation Status of Gene TP53
in Cancer LGG, Error Type:word_error
train_label
test_label
Percentage
0.0 0.2 0.4 0.6 0.8 1.0
0 1 4
Label in Decimal Format

#### 3.3 Brain Cancer

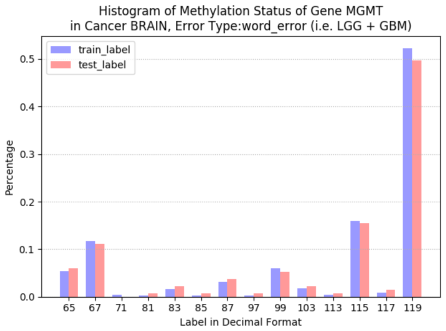

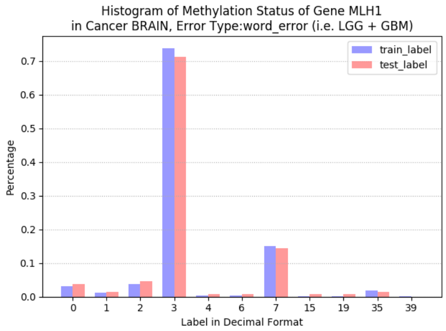

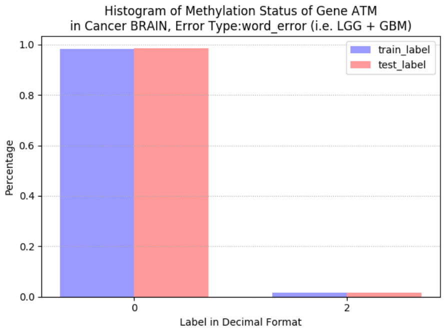

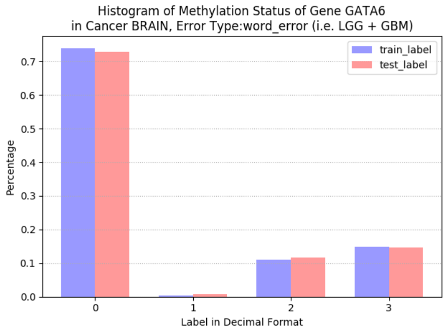

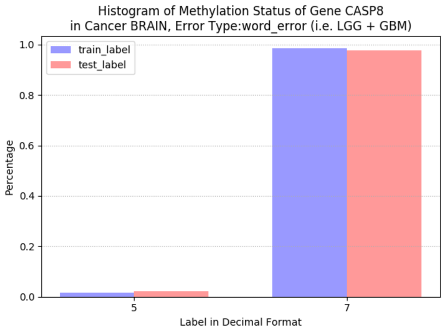

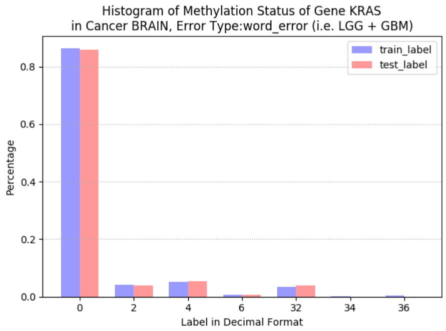

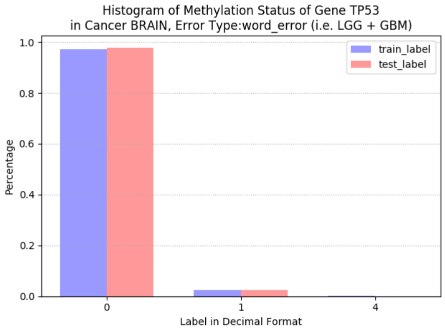

#### 3.4 LUAD

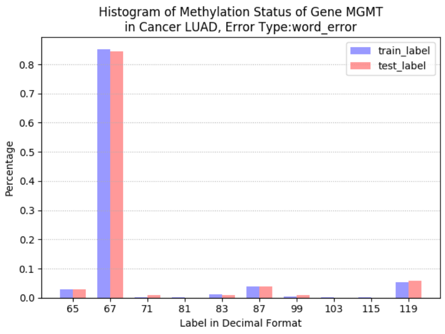
Histogram of Methylation Status of Gene MGMT
in Cancer LUAD, Error Type:word_error
train_label
test_label
Percentage
0.0
0.1
0.2
0.3
0.4
0.5
0.6
0.7
0.8
65
67
71
81
83
87
99
103
115
119
Label in Decimal Format

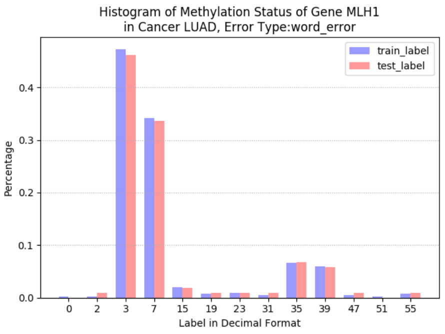
Histogram of Methylation Status of Gene MLH1
in Cancer LUAD, Error Type:word_error
train_label
test_label
Percentage
0.0
0.1
0.2
0.3
0.4
0
2
3
7
15
19
23
31
35
39
47
51
55
Label in Decimal Format

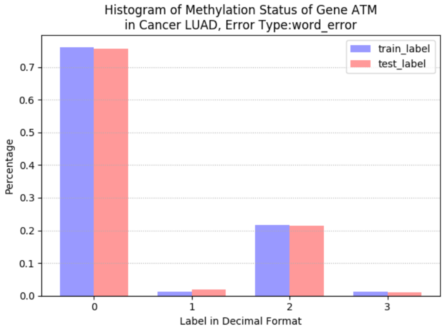
Histogram of Methylation Status of Gene ATM
in Cancer LUAD, Error Type:word_error
train_label
test_label
Percentage
0.0
0.1
0.2
0.3
0.4
0.5
0.6
0.7
0
1
2
3
Label in Decimal Format

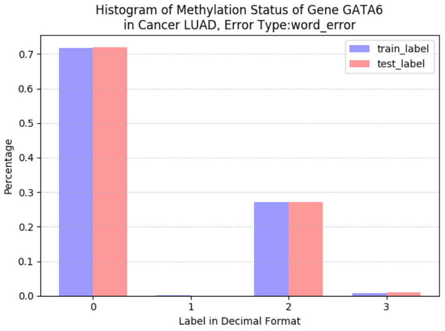
Histogram of Methylation Status of Gene GATA6
in Cancer LUAD, Error Type:word_error
train_label
test_label
Percentage
0.0
0.1
0.2
0.3
0.4
0.5
0.6
0.7
0
1
2
3
Label in Decimal Format

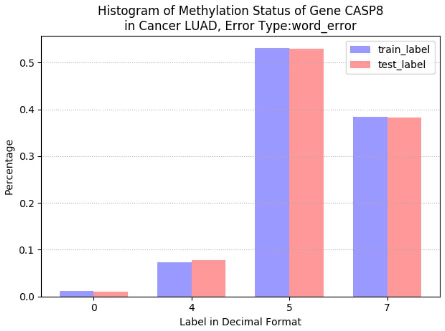
Histogram of Methylation Status of Gene CASP8
in Cancer LUAD, Error Type:word_error
train_label
test_label
Percentage
0.0
0.1
0.2
0.3
0.4
0.5
0
4
5
7
Label in Decimal Format

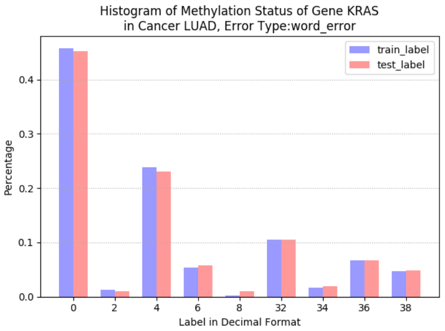
Histogram of Methylation Status of Gene KRAS
in Cancer LUAD, Error Type:word_error
train_label
test_label
Percentage
0.0
0.1
0.2
0.3
0.4
0
2
4
6
8
32
34
36
38
Label in Decimal Format

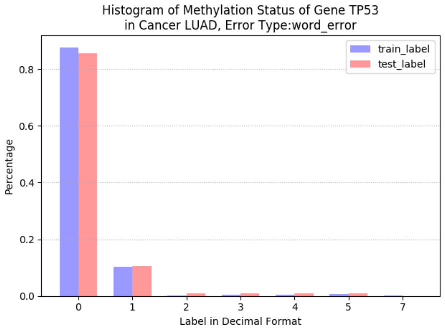
Histogram of Methylation Status of Gene TP53
in Cancer LUAD, Error Type:word_error
train_label
test_label
Percentage
0.0
0.2
0.4
0.6
0.8
0
1
2
3
4
5
7
Label in Decimal Format

#### 3.5 LUSC

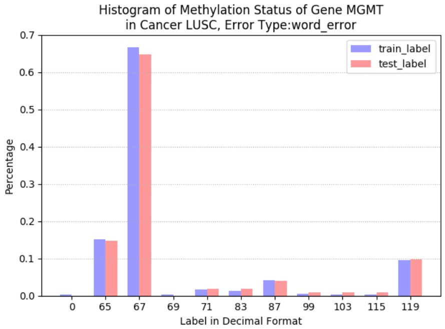
Histogram of Methylation Status of Gene MGMT
in Cancer LUSC, Error Type:word_error
train_label
test_label
Percentage
0.0
0.1
0.2
0.3
0.4
0.5
0.6
0.7
0
65
67
69
71
83
87
99
103
115
119
Label in Decimal Format

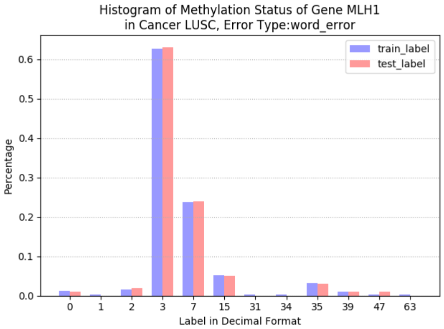
Histogram of Methylation Status of Gene MLH1
in Cancer LUSC, Error Type:word_error
train_label
test_label
Percentage
0.0
0.1
0.2
0.3
0.4
0.5
0.6
0
1
2
3
7
15
31
34
35
39
47
63
Label in Decimal Format

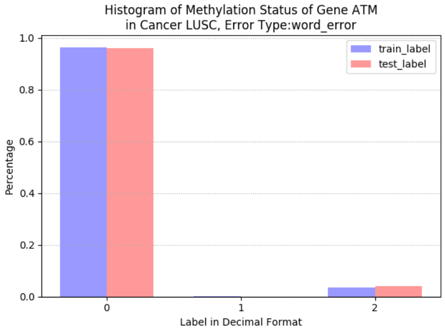
Histogram of Methylation Status of Gene ATM
in Cancer LUSC, Error Type:word_error
train_label
test_label
Percentage
0.0
0.2
0.4
0.6
0.8
1.0
0
1
2
Label in Decimal Format

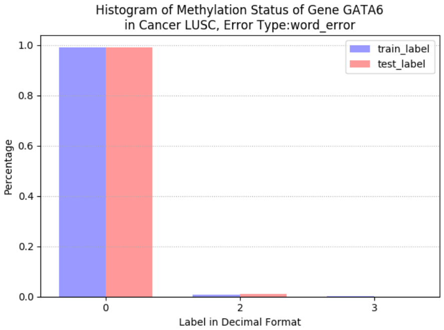
Histogram of Methylation Status of Gene GATA6
in Cancer LUSC, Error Type:word_error
train_label
test_label
Percentage
0.0
0.2
0.4
0.6
0.8
1.0
0
2
3
Label in Decimal Format

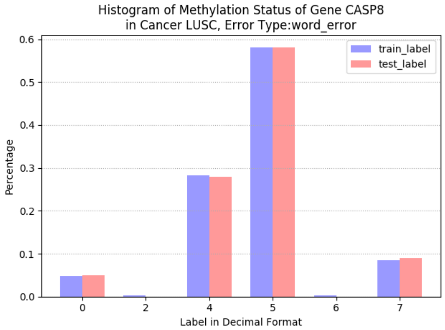
Histogram of Methylation Status of Gene CASP8
in Cancer LUSC, Error Type:word_error
train_label
test_label
Percentage
0.0
0.1
0.2
0.3
0.4
0.5
0.6
0
2
4
5
6
7
Label in Decimal Format

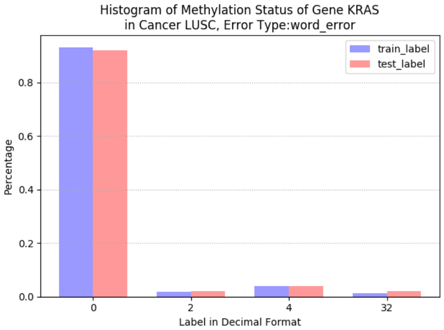
Histogram of Methylation Status of Gene KRAS
in Cancer LUSC, Error Type:word_error
train_label
test_label
Percentage
0.0
0.2
0.4
0.6
0.8
0
2
4
32
Label in Decimal Format

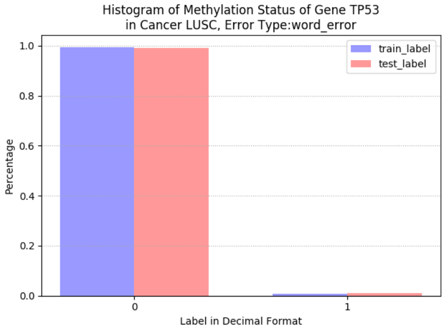
Histogram of Methylation Status of Gene TP53
in Cancer LUSC, Error Type:word_error
train_label
test_label
Percentage
0.0
0.2
0.4
0.6
0.8
1.0
0
1
Label in Decimal Format

#### 3.6 Lung Cancer

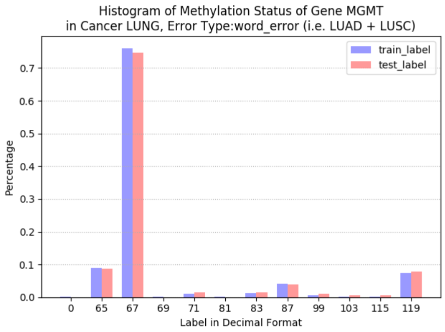

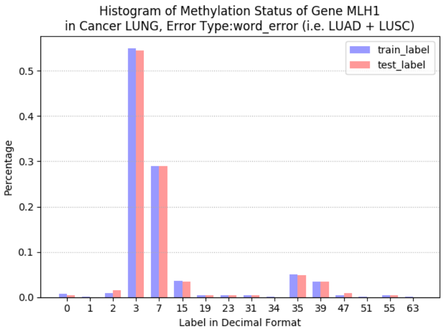

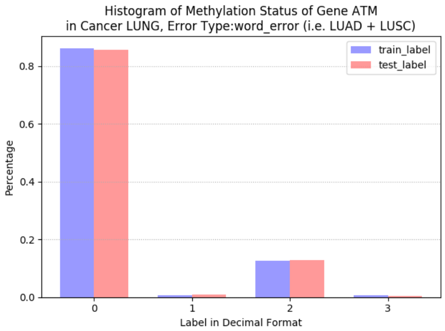

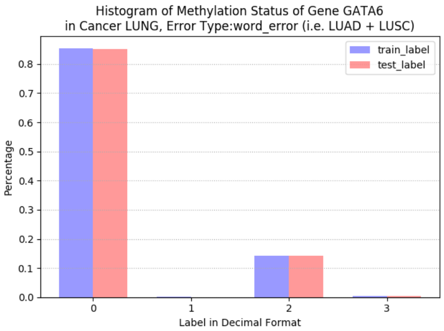

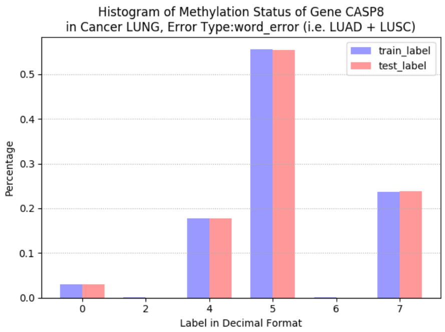

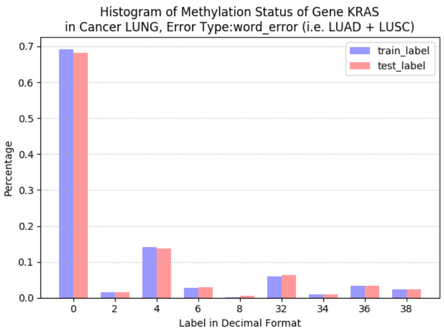

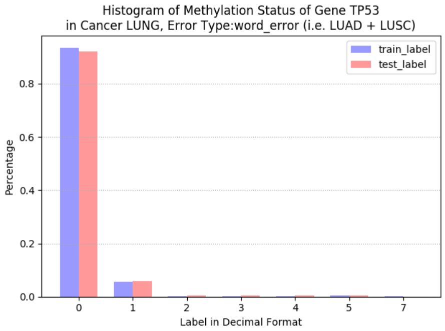

#### 3.7 STAD

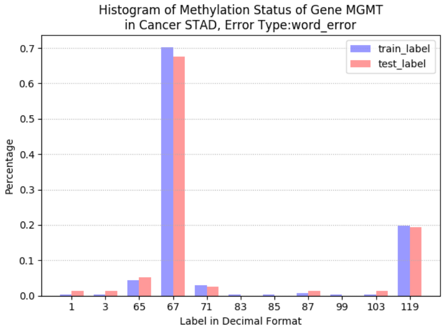
Histogram of Methylation Status of Gene MGMT
in Cancer STAD, Error Type:word_error
train_label
test_label
Percentage
0.0
0.1
0.2
0.3
0.4
0.5
0.6
0.7
1
3
65
67
71
83
85
87
99
103
119
Label in Decimal Format

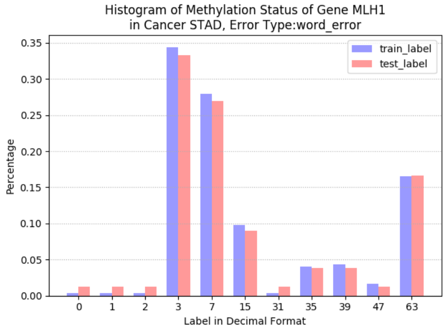
Histogram of Methylation Status of Gene MLH1
in Cancer STAD, Error Type:word_error
train_label
test_label
Percentage
0.00
0.05
0.10
0.15
0.20
0.25
0.30
0.35
0
1
2
3
7
15
31
35
39
47
63
Label in Decimal Format

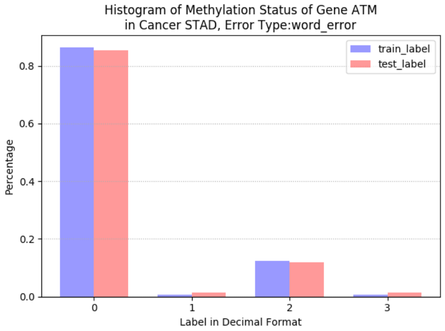
Histogram of Methylation Status of Gene ATM
in Cancer STAD, Error Type:word_error
train_label
test_label
Percentage
0.0
0.2
0.4
0.6
0.8
0
1
2
3
Label in Decimal Format

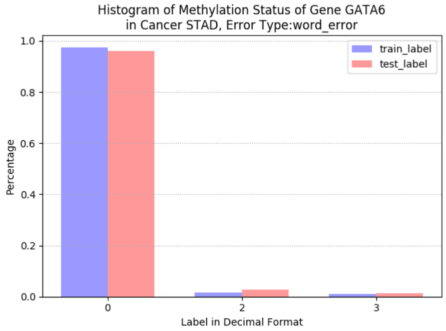
Histogram of Methylation Status of Gene GATA6
in Cancer STAD, Error Type:word_error
train_label
test_label
Percentage
0.0
0.2
0.4
0.6
0.8
1.0
0
2
3
Label in Decimal Format

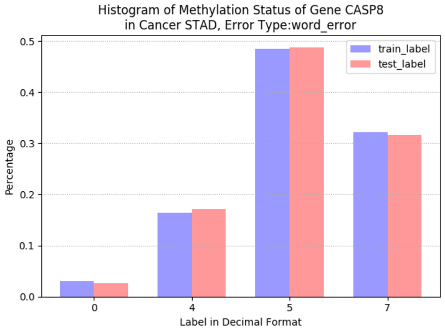
Histogram of Methylation Status of Gene CASP8
in Cancer STAD, Error Type:word_error
train_label
test_label
Percentage
0.0
0.1
0.2
0.3
0.4
0.5
0
4
5
7
Label in Decimal Format

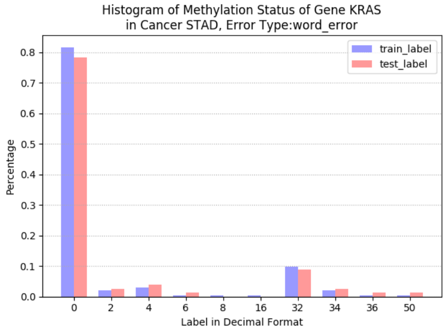
Histogram of Methylation Status of Gene KRAS
in Cancer STAD, Error Type:word_error
train_label
test_label
Percentage
0.0
0.1
0.2
0.3
0.4
0.5
0.6
0.7
0.8
0
2
4
6
8
16
32
34
36
50
Label in Decimal Format

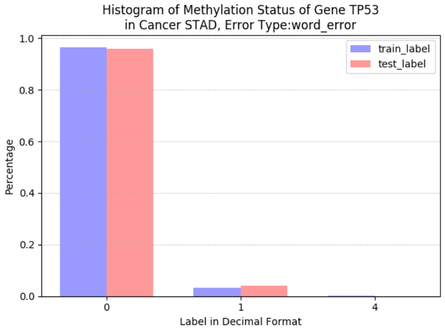
Histogram of Methylation Status of Gene TP53
in Cancer STAD, Error Type:word_error
train_label
test_label
Percentage
0.0
0.2
0.4
0.6
0.8
1.0
0
1
4
Label in Decimal Format

### 4 Heatmaps of Gene Expressions

The presented results include gene expression heatmaps for all genes under consideration, and for all cancer types.

#### 4.1 GBM

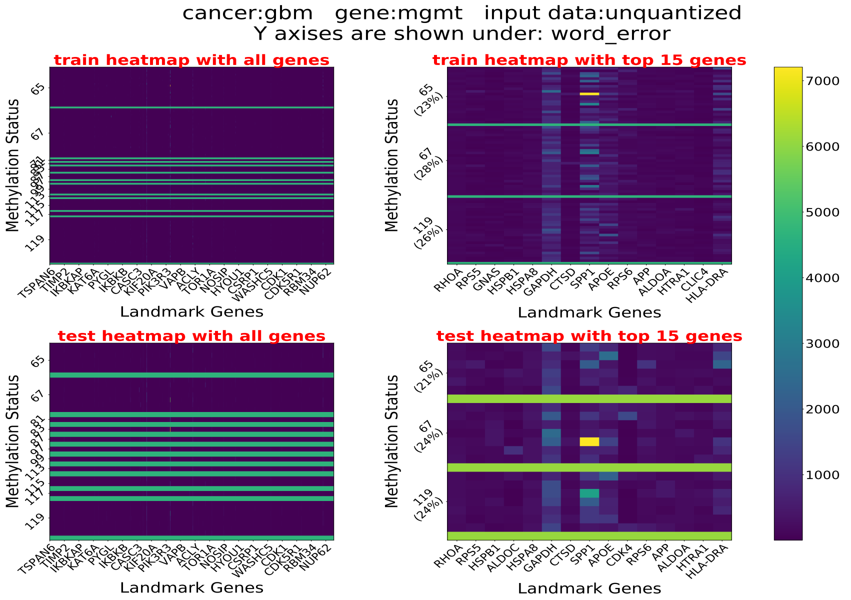

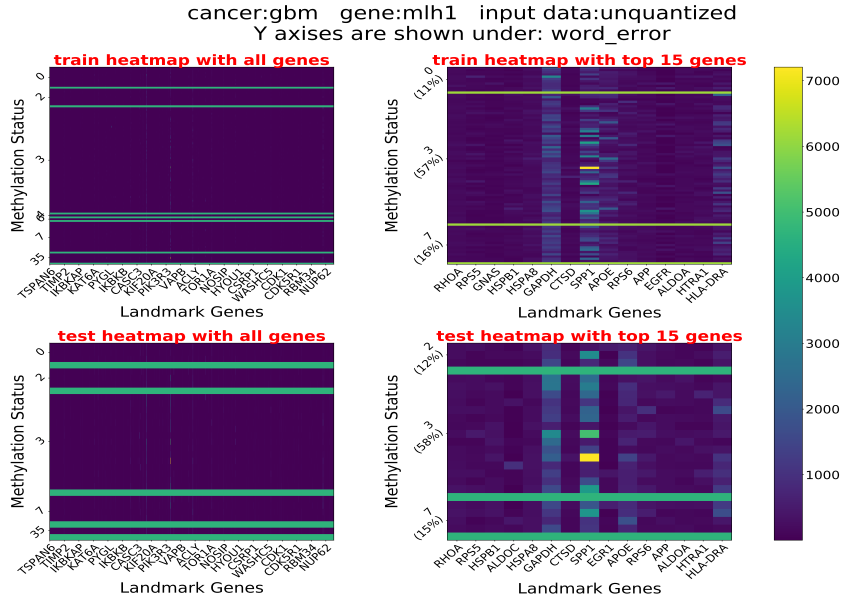

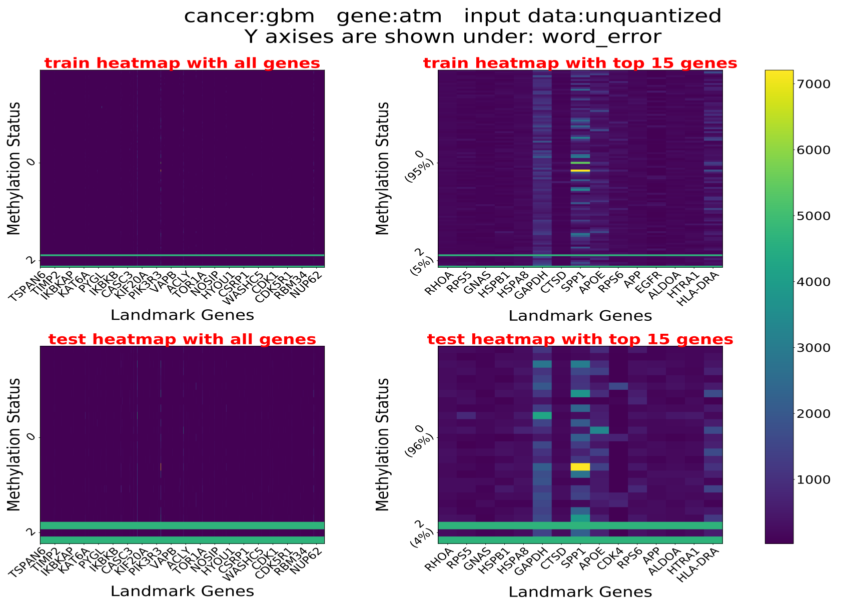
cancer:gbm gene:atm input data:unquantized
Y axises are shown under: word_error
train heatmap with all genes
train heatmap with top 15 genes
test heatmap with all genes
test heatmap with top 15 genes
Methylation Status
Landmark Genes
0
2
0 (95%)
2 (5%)
0 (96%)
2 (4%)
TSPAN6 TIMP2 IKBKAP KAT6A PYGL IKBKB CASC3 KIF20A PIK3R3 VAPB ACLY TOR1A NOSIP HYOU1 CSRP1 WASHC5 CDK1 CDK5R1 RBM34 NUP62
RHOA RPS5 GNAS HSPB1 HSPA8 GAPDH CTSD SPP1 APOE RPS6 APP EGFR ALDOA HTRA1 HLA-DRA
RHOA RPS5 GNAS HSPB1 HSPA8 GAPDH CTSD SPP1 APOE CDK4 RPS6 APP ALDOA HTRA1 HLA-DRA
1000
2000
3000
4000
5000
6000
7000

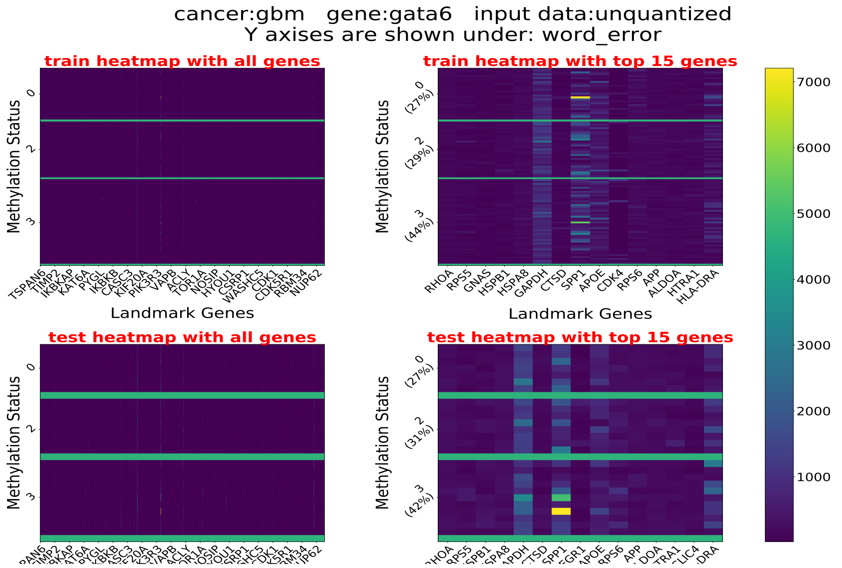
cancer:gbm gene:gata6 input data:unquantized
Y axises are shown under: word_error
train heatmap with all genes
train heatmap with top 15 genes
test heatmap with all genes
test heatmap with top 15 genes
Methylation Status
Landmark Genes
0
2
3
0 (27%)
2 (29%)
3 (44%)
0 (27%)
2 (31%)
3 (42%)
TSPAN6 TIMP2 IKBKAP KAT6A PYGL IKBKB CASC3 KIF20A PIK3R3 VAPB ACLY TOR1A NOSIP HYOU1 CSRP1 WASHC5 CDK1 CDK5R1 RBM34 NUP62
RHOA RPS5 GNAS HSPB1 HSPA8 GAPDH CTSD SPP1 APOE CDK4 RPS6 APP ALDOA HTRA1 HLA-DRA
1000
2000
3000
4000
5000
6000
7000

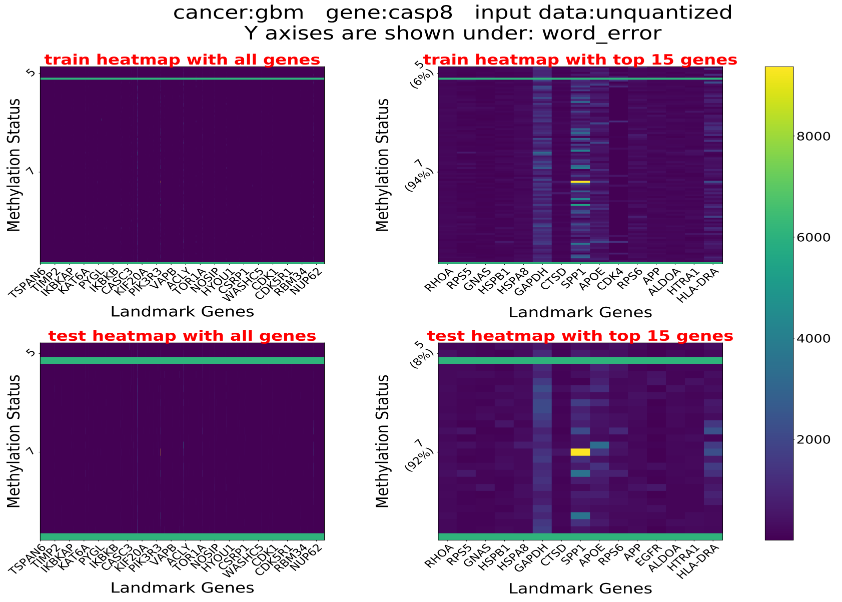
cancer:gbm gene:casp8 input data:unquantized
Y axises are shown under: word_error
train heatmap with all genes
train heatmap with top 15 genes
test heatmap with all genes
test heatmap with top 15 genes
Methylation Status
Landmark Genes
5
7
5 (6%)
7 (94%)
5 (8%)
7 (92%)
TSPAN6 TIMP2 IKBKAP KAT6A PYGL IKBKB CASC3 KIF20A PIK3R3 VAPB ACLY TOR1A NOSIP HYOU1 CSRP1 WASHC5 CDK1 CDK5R1 RBM34 NUP62
RHOA RPS5 GNAS HSPB1 HSPA8 GAPDH CTSD SPP1 APOE CDK4 RPS6 APP ALDOA HTRA1 HLA-DRA
RHOA RPS5 GNAS HSPB1 HSPA8 GAPDH CTSD SPP1 APOE RPS6 APP EGFR ALDOA HTRA1 HLA-DRA
2000
4000
6000
8000

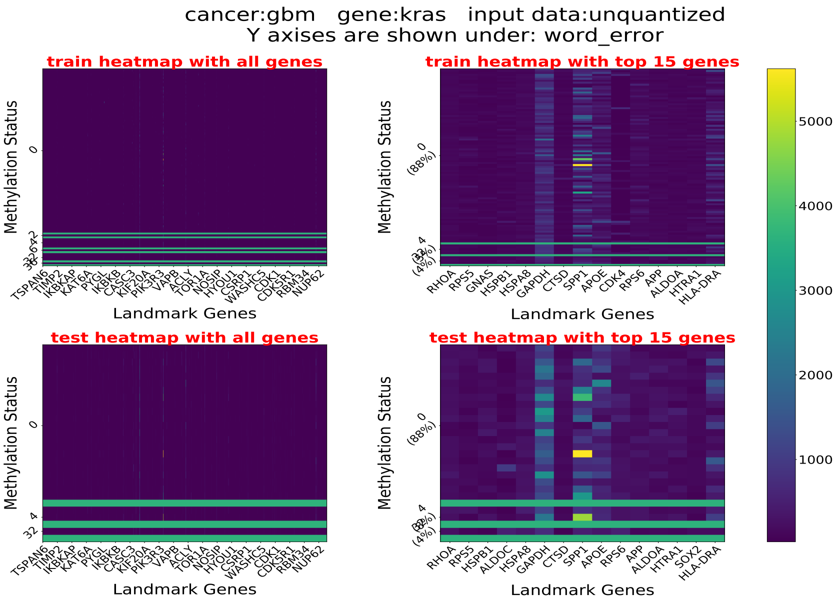
cancer:gbm gene:kras input data:unquantized
Y axises are shown under: word_error
train heatmap with all genes
train heatmap with top 15 genes
test heatmap with all genes
test heatmap with top 15 genes
Methylation Status
Landmark Genes
0
(88%)
4
(4%)
TSPAN6
TIMP2
IKBKAP
KAT6A
PYGL
IKBKB
CASC3
KIF20A
PIK3R3
VAPB
ACLY
TOR1A
NOSIP
HYOU1
CSRP1
WASHC5
CDK1
CDK5R1
RBM34
NUP62
RHOA
RPS5
GNAS
HSPB1
HSPA8
GAPDH
CTSD
SPP1
APOE
CDK4
RPS6
APP
ALDOA
HTRA1
HLA-DRA
ALDOC
SOX2
5000
4000
3000
2000
1000

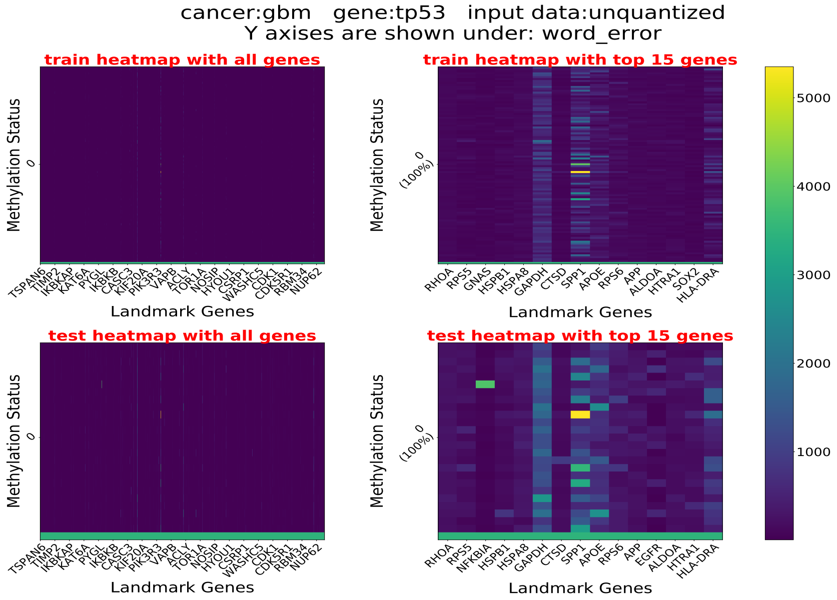
cancer:gbm gene:tp53 input data:unquantized
Y axises are shown under: word_error
train heatmap with all genes
train heatmap with top 15 genes
test heatmap with all genes
test heatmap with top 15 genes
Methylation Status
Landmark Genes
0
(100%)
TSPAN6
TIMP2
IKBKAP
KAT6A
PYGL
IKBKB
CASC3
KIF20A
PIK3R3
VAPB
ACLY
TOR1A
NOSIP
HYOU1
CSRP1
WASHC5
CDK1
CDK5R1
RBM34
NUP62
RHOA
RPS5
GNAS
HSPB1
HSPA8
GAPDH
CTSD
SPP1
APOE
RPS6
APP
ALDOA
HTRA1
SOX2
HLA-DRA
NFKBIA
EGFR
5000
4000
3000
2000
1000

#### 4.2 LGG

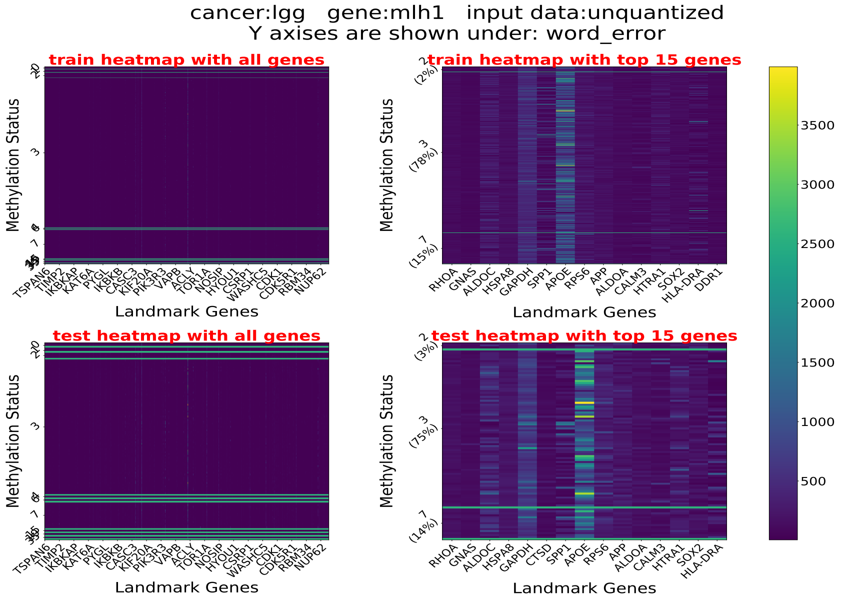
cancer:lgg gene:mlh1 input data:unquantized
Y axises are shown under: word_error
train heatmap with all genes
train heatmap with top 15 genes
test heatmap with all genes
test heatmap with top 15 genes
Methylation Status
Landmark Genes
2 (2%)
3 (78%)
7 (15%)
2 (3%)
3 (75%)
7 (14%)
TSPAN6
TIMP2
IKBKAP
KAT6A
PYGL
IKBKB
CASC3
KIF20A
PIK3R3
VAPB
ACLY
TOR1A
NOSIP
HYOU1
CSRP1
WASHC5
CDK1
CDK5R1
RBM34
NUP62
RHOA
GNAS
ALDOC
HSPA8
GAPDH
CTSD
SPP1
APOE
RPS6
APP
ALDOA
CALM3
HTRA1
SOX2
HLA-DRA
DDR1
3500
3000
2500
2000
1500
1000
500

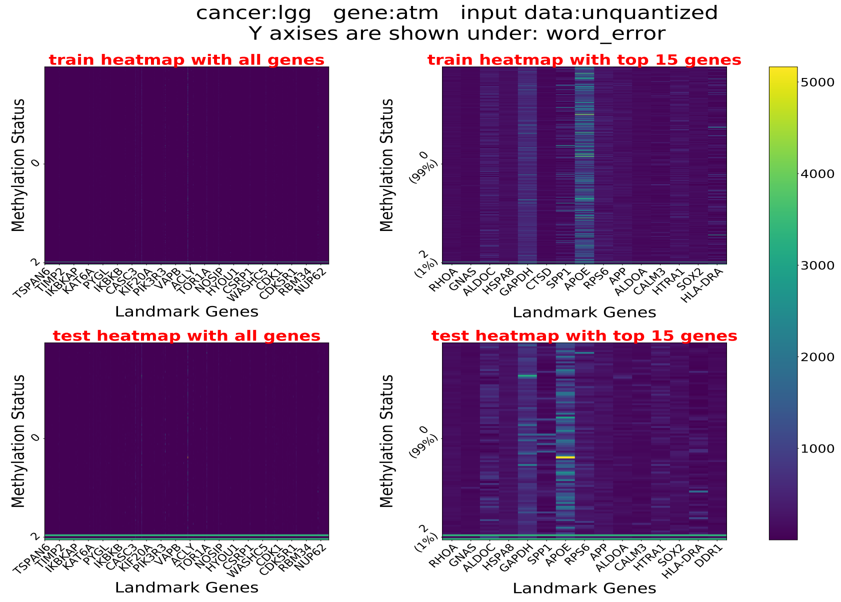
cancer:lgg gene:atm input data:unquantized
Y axises are shown under: word_error
train heatmap with all genes
train heatmap with top 15 genes
test heatmap with all genes
test heatmap with top 15 genes
Methylation Status
Landmark Genes
0 (99%)
2 (1%)
TSPAN6
TIMP2
IKBKAP
KAT6A
PYGL
IKBKB
CASC3
KIF20A
PIK3R3
VAPB
ACLY
TOR1A
NOSIP
HYOU1
CSRP1
WASHC5
CDK1
CDK5R1
RBM34
NUP62
RHOA
GNAS
ALDOC
HSPA8
GAPDH
CTSD
SPP1
APOE
RPS6
APP
ALDOA
CALM3
HTRA1
SOX2
HLA-DRA
DDR1
5000
4000
3000
2000
1000

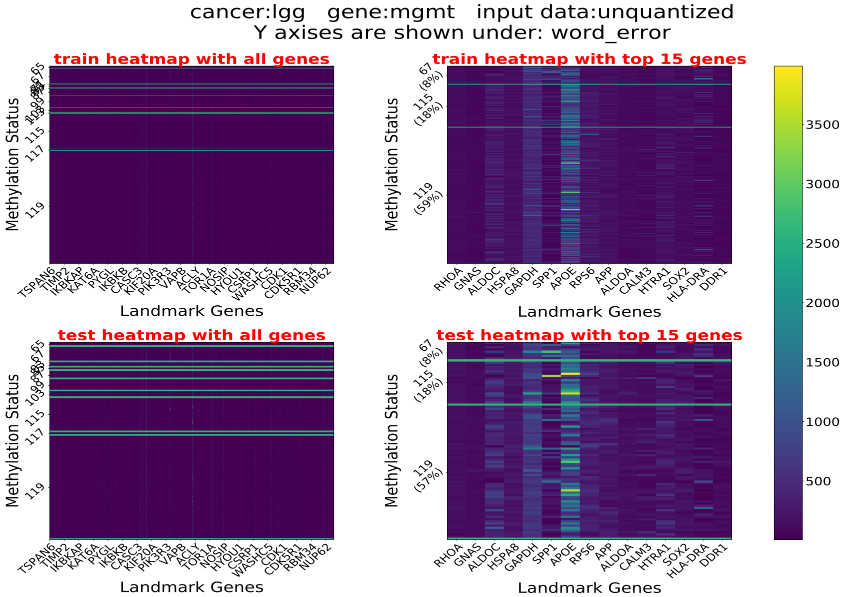
cancer:lgg gene:mgmt input data:unquantized
Y axises are shown under: word_error
train heatmap with all genes
train heatmap with top 15 genes
test heatmap with all genes
test heatmap with top 15 genes
Methylation Status
Landmark Genes
TSPAN6 TIMP2 IKBKAP KAT6A PTGL IKBKB CASC3 KIF20A PIK3R3 VAPB ACLY TOR1A NOSIP HYOU1 CSRP1 WASHC5 CDK1 CDK5R1 RBM34 NUP62
RHOA GNAS ALDOC HSPA8 GAPDH SPP1 APOE RPS6 APP ALDOA CALM3 HTRA1 SOX2 HLA-DRA DDR1
67 (8%)
115 (18%)
119 (59%)
119 (57%)
3500 3000 2500 2000 1500 1000 500

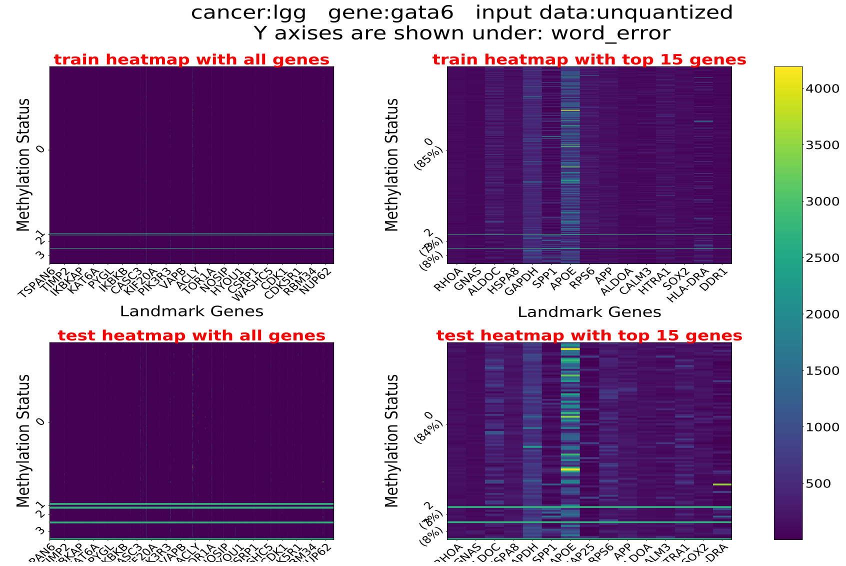
cancer:lgg gene:gata6 input data:unquantized
Y axises are shown under: word_error
train heatmap with all genes
train heatmap with top 15 genes
test heatmap with all genes
test heatmap with top 15 genes
Methylation Status
Landmark Genes
0 (85%)
0 (84%)
4000 3500 3000 2500 2000 1500 1000 500

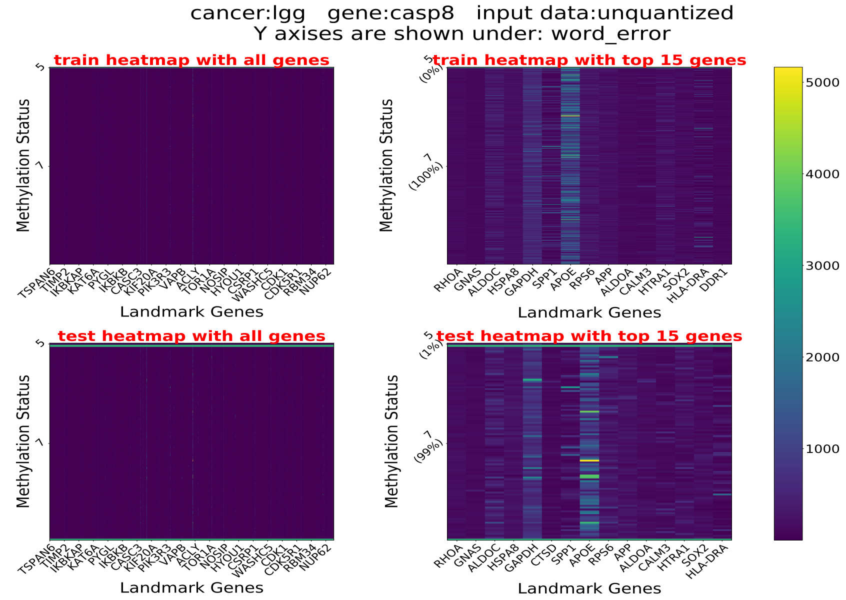
cancer:lgg gene:casp8 input data:unquantized
Y axises are shown under: word_error
train heatmap with all genes
train heatmap with top 15 genes
test heatmap with all genes
test heatmap with top 15 genes
Methylation Status
Landmark Genes
5 (0%)
7 (100%)
5 (1%)
7 (99%)
RHOA GNAS ALDOC HSPA8 GAPDH CTSD SPP1 APOE RPS6 APP ALDOA CALM3 HTRA1 SOX2 HLA-DRA
5000 4000 3000 2000 1000

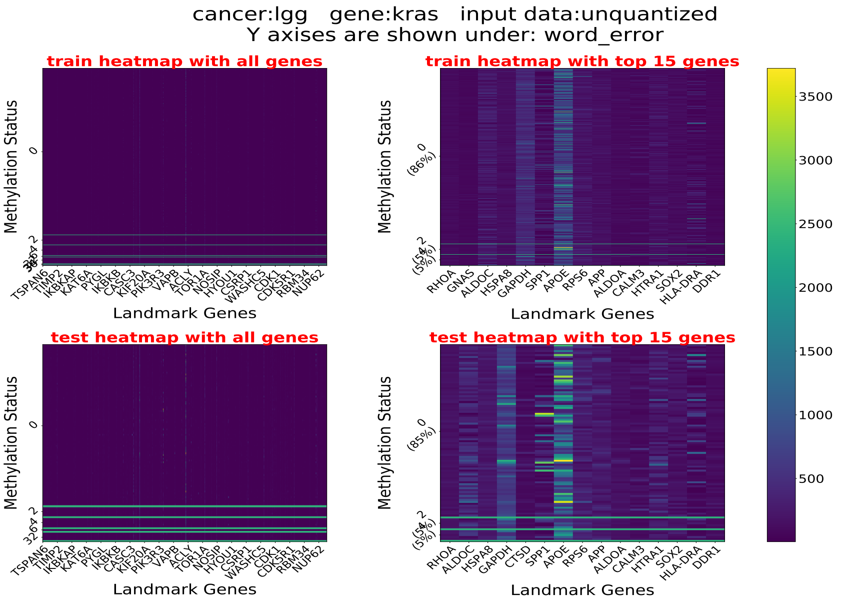
cancer:lgg gene:kras input data:unquantized
Y axises are shown under: word_error
train heatmap with all genes
train heatmap with top 15 genes
test heatmap with all genes
test heatmap with top 15 genes
Methylation Status
Landmark Genes
TSPAN6
TIMP2
IKBKAP
KAT6A
PYGL
IKBKB
CASC3
KIF20A
PIK3R3
VAPB
ACLY
TOR1A
NOSIP
HYOU1
CSRP1
WASHC5
CDK1
CDK5R1
RBM34
NUP62
RHOA
GNAS
ALDOC
HSPA8
GAPDH
CTSD
SPP1
APOE
RPS6
APP
ALDOA
CALM3
HTRA1
SOX2
HLA-DRA
DDR1
(86%)
(85%)
3500
3000
2500
2000
1500
1000
500

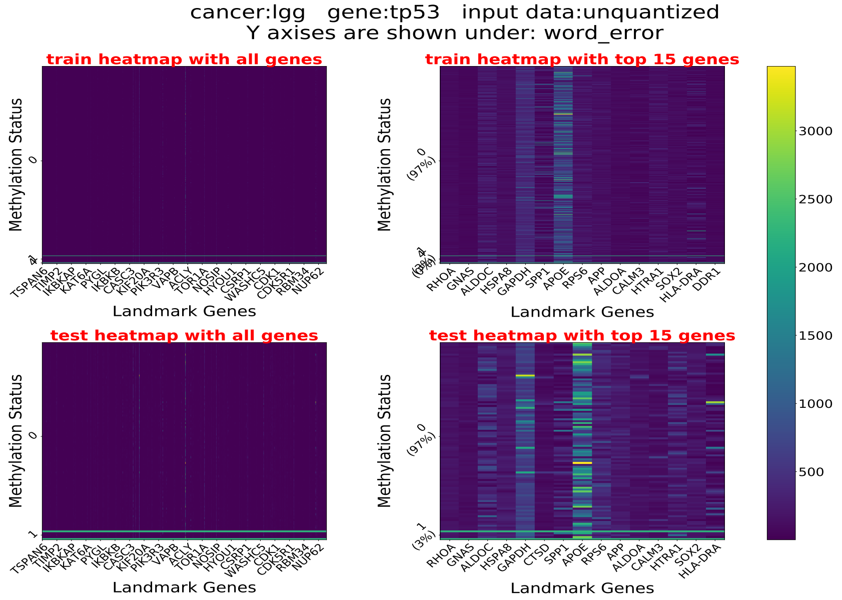
cancer:lgg gene:tp53 input data:unquantized
Y axises are shown under: word_error
train heatmap with all genes
train heatmap with top 15 genes
test heatmap with all genes
test heatmap with top 15 genes
Methylation Status
Landmark Genes
(97%)
(3%)
3000
2500
2000
1500
1000
500

#### 4.3 Brain Cancer

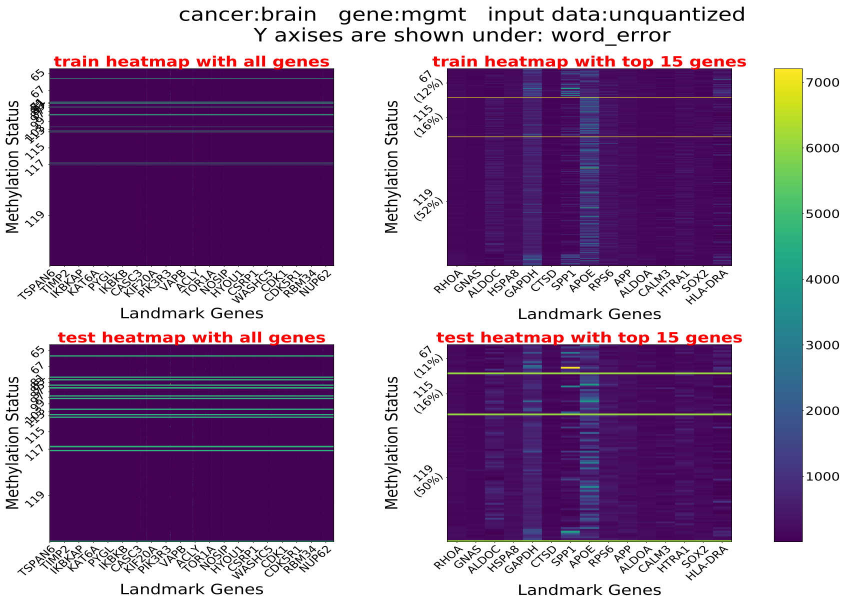

vspace-.15in

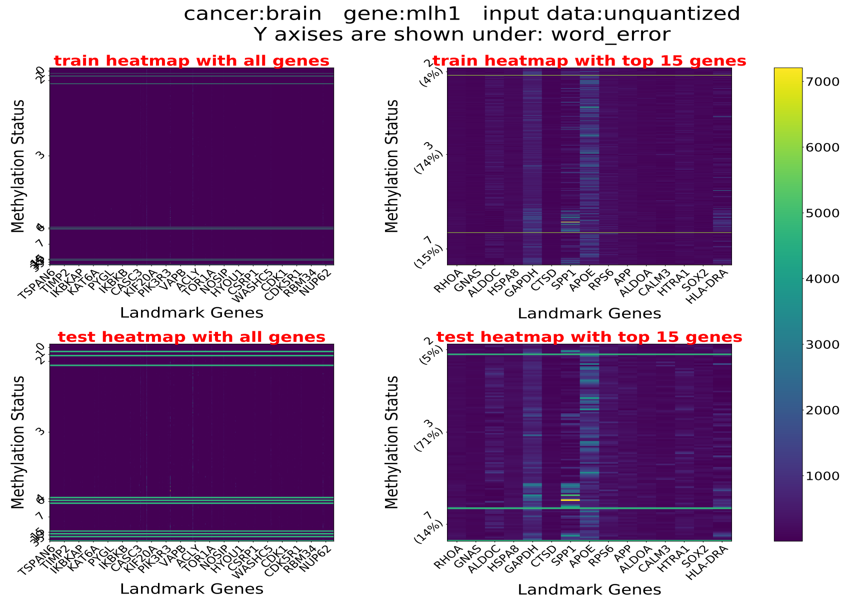

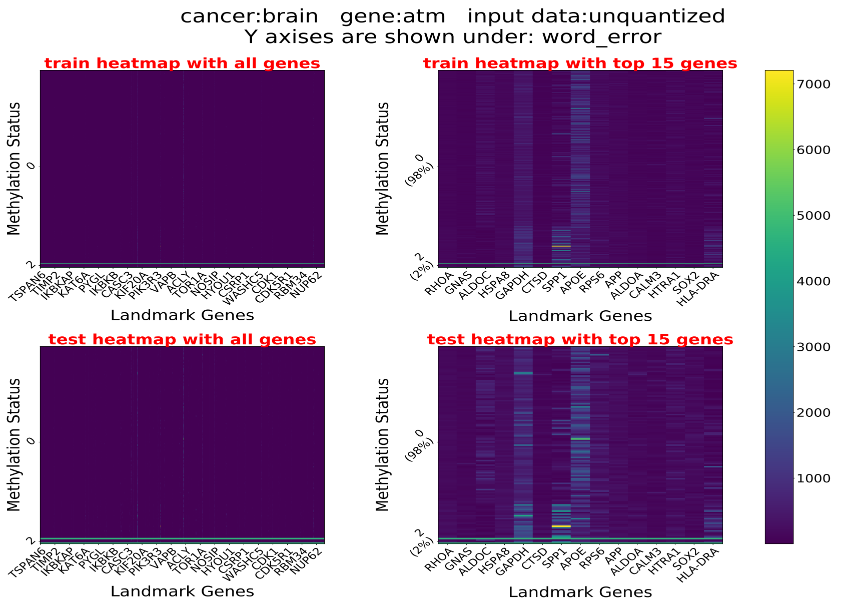
cancer:brain gene:atm input data:unquantized
Y axises are shown under: word_error
train heatmap with all genes
train heatmap with top 15 genes
test heatmap with all genes
test heatmap with top 15 genes
Methylation Status
Landmark Genes
0 (98%)
2 (2%)
TSPAN6 TIMP2 IKBKAP KAT6A PYGL IKBKB CASC3 KIF20A PIK3R3 VAPB ACLY TOR1A NOSIP HYOU1 CSRP1 WASHC5 CDK1 CDK5R1 RBM34 NUP62
RHOA GNAS ALDOC HSPA8 GAPDH CTSD SPP1 APOE RPS6 APP ALDOA CALM3 HTRA1 SOX2 HLA-DRA
1000 2000 3000 4000 5000 6000 7000

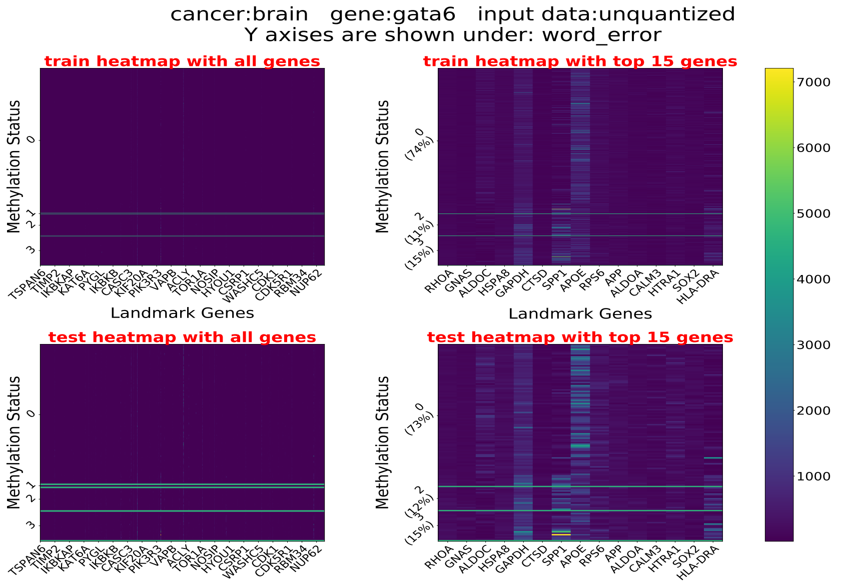
cancer:brain gene:gata6 input data:unquantized
Y axises are shown under: word_error
train heatmap with all genes
train heatmap with top 15 genes
test heatmap with all genes
test heatmap with top 15 genes
Methylation Status
Landmark Genes
0 (74%)
2 (11%)
3 (15%)
0 (73%)
2 (12%)
3 (15%)
TSPAN6 TIMP2 IKBKAP KAT6A PYGL IKBKB CASC3 KIF20A PIK3R3 VAPB ACLY TOR1A NOSIP HYOU1 CSRP1 WASHC5 CDK1 CDK5R1 RBM34 NUP62
RHOA GNAS ALDOC HSPA8 GAPDH CTSD SPP1 APOE RPS6 APP ALDOA CALM3 HTRA1 SOX2 HLA-DRA
1000 2000 3000 4000 5000 6000 7000

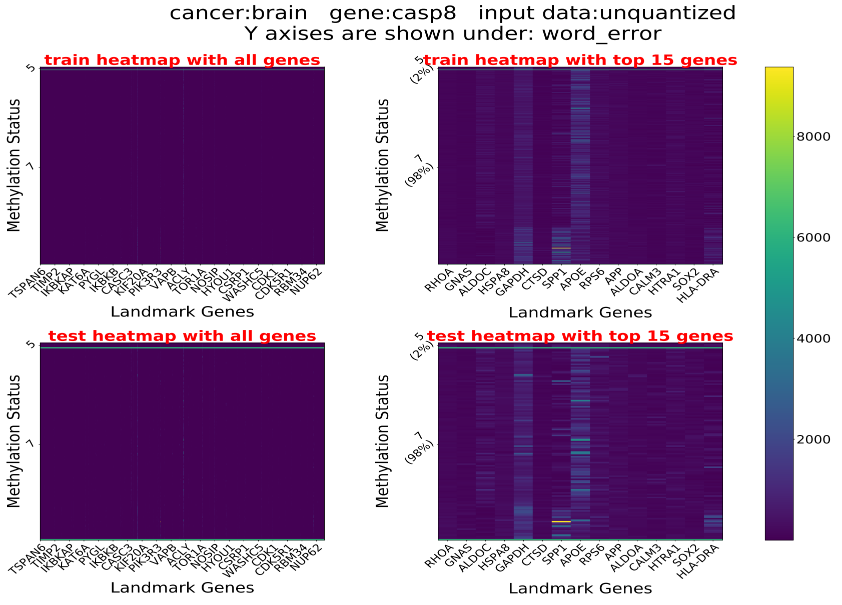
cancer:brain gene:casp8 input data:unquantized
Y axises are shown under: word_error
train heatmap with all genes
train heatmap with top 15 genes
test heatmap with all genes
test heatmap with top 15 genes
Methylation Status
Landmark Genes
5 (2%)
7 (98%)
TSPAN6 TIMP2 IKBKAP KAT6A PYGL IKBKB CASC3 KIF20A PIK3R3 VAPB ACLY TOR1A NOSIP HYOU1 CSRP1 WASHC5 CDK1 CDK5R1 RBM34 NUP62
RHOA GNAS ALDOC HSPA8 GAPDH CTSD SPP1 APOE RPS6 APP ALDOA CALM3 HTRA1 SOX2 HLA-DRA
2000 4000 6000 8000

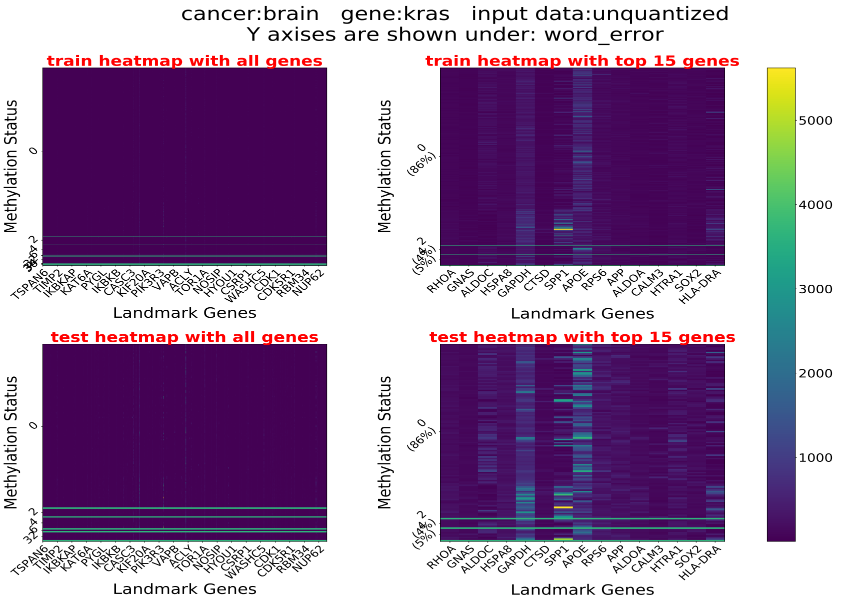
cancer:brain gene:kras input data:unquantized
Y axises are shown under: word_error
train heatmap with all genes
train heatmap with top 15 genes
test heatmap with all genes
test heatmap with top 15 genes
Methylation Status
Landmark Genes
TSPAN6
TIMP2
IKBKAP
KAT6A
PTGL
IKBKB
CASC3
KIF20A
PIK3R3
VAPB
ACLY
TOR1A
NOSIP
HYOU1
CSRP1
WASHC5
CDK1
CDK5R1
RBM34
NUP62
RHOA
GNAS
ALDOC
HSPA8
GAPDH
CTSD
SPP1
APOE
RPS6
APP
ALDOA
CALM3
HTRA1
SOX2
HLA-DRA
0
(86%)
5000
4000
3000
2000
1000

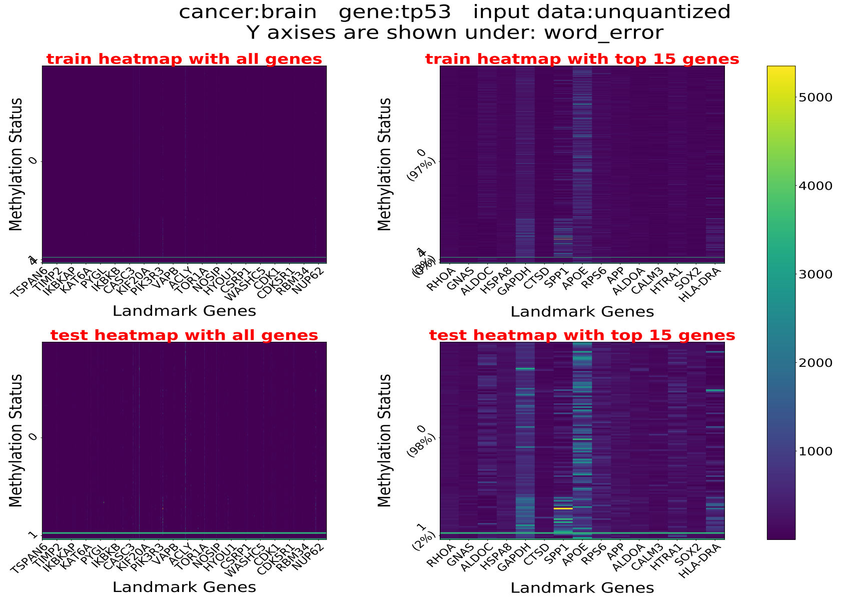
cancer:brain gene:tp53 input data:unquantized
Y axises are shown under: word_error
train heatmap with all genes
train heatmap with top 15 genes
test heatmap with all genes
test heatmap with top 15 genes
Methylation Status
Landmark Genes
TSPAN6
TIMP2
IKBKAP
KAT6A
PTGL
IKBKB
CASC3
KIF20A
PIK3R3
VAPB
ACLY
TOR1A
NOSIP
HYOU1
CSRP1
WASHC5
CDK1
CDK5R1
RBM34
NUP62
RHOA
GNAS
ALDOC
HSPA8
GAPDH
CTSD
SPP1
APOE
RPS6
APP
ALDOA
CALM3
HTRA1
SOX2
HLA-DRA
0
(97%)
(98%)
1
(2%)
5000
4000
3000
2000
1000

#### 4.4 LUAD

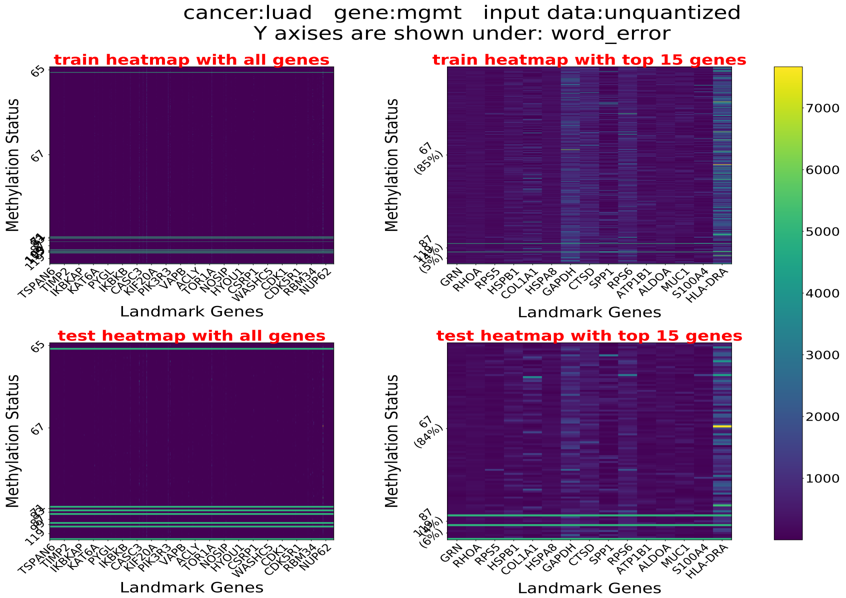
cancer:luad gene:mgmt input data:unquantized
Y axises are shown under: word_error
train heatmap with all genes
train heatmap with top 15 genes
test heatmap with all genes
test heatmap with top 15 genes
Methylation Status
Landmark Genes

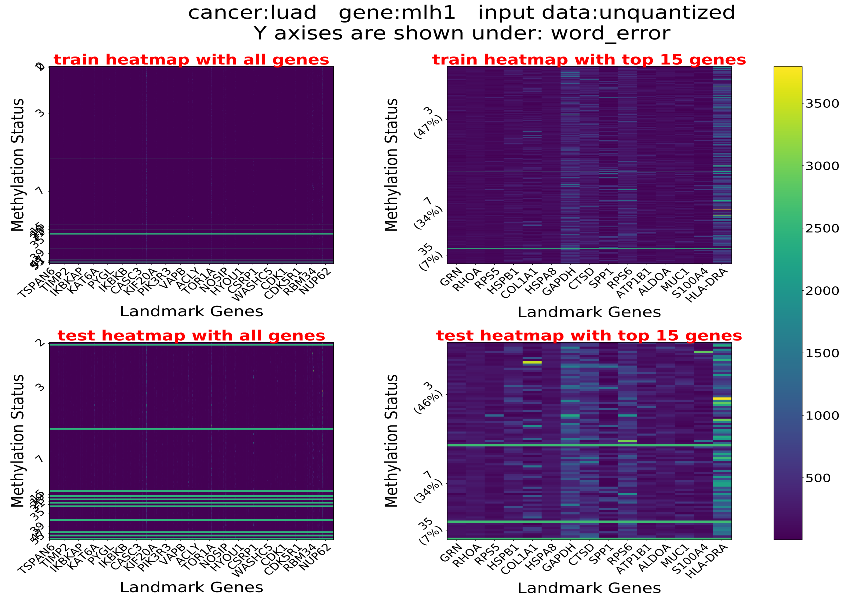
cancer:luad gene:mlh1 input data:unquantized
Y axises are shown under: word_error
train heatmap with all genes
train heatmap with top 15 genes
test heatmap with all genes
test heatmap with top 15 genes
Methylation Status
Landmark Genes

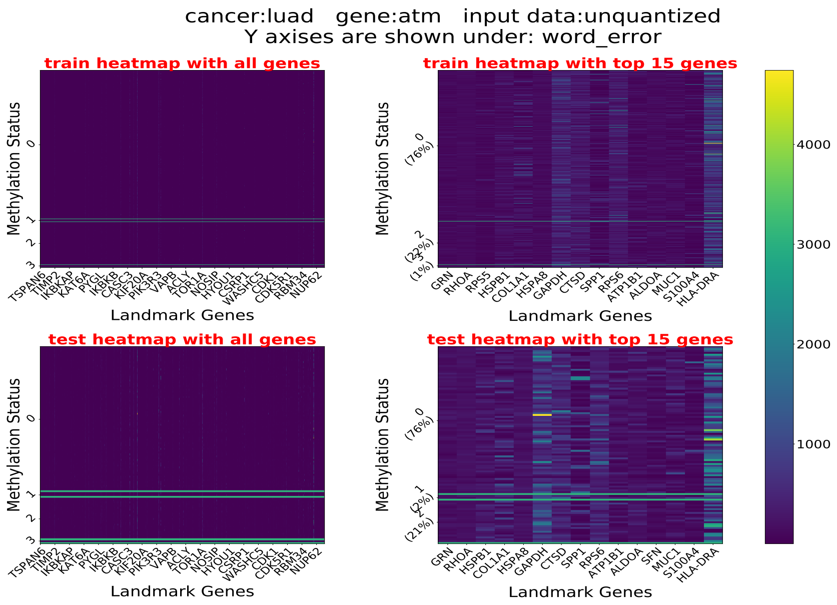
cancer:luad gene:atm input data:unquantized
Y axises are shown under: word_error
train heatmap with all genes
train heatmap with top 15 genes
test heatmap with all genes
test heatmap with top 15 genes
Methylation Status
Landmark Genes
TSPAN6 TIMP2 IKBKAP KAT6A PYGL IKBKB CASC3 KIF20A PIK3R3 VAPB ACLY TOR1A NOSIP HYOU1 CSRP1 WASHC5 CDK1 CDK5R1 RBM34 NUP62
GRN RHOA RPS5 HSPB1 COL1A1 HSPA8 GAPDH CTSD SPP1 RPS6 ATP1B1 ALDOA MUC1 S100A4 HLA-DRA
GRN RHOA HSPB1 COL1A1 HSPA8 GAPDH CTSD SPP1 RPS6 ATP1B1 ALDOA SFN MUC1 S100A4 HLA-DRA
0 (76%)
2 (22%)
3 (1%)
1 (2%)
3 (21%)
4000
3000
2000
1000

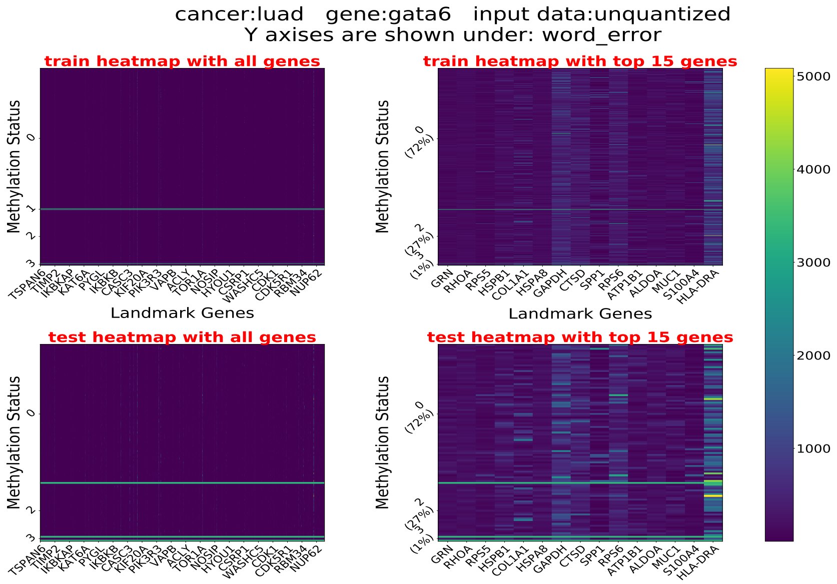
cancer:luad gene:gata6 input data:unquantized
Y axises are shown under: word_error
train heatmap with all genes
train heatmap with top 15 genes
test heatmap with all genes
test heatmap with top 15 genes
Methylation Status
Landmark Genes
TSPAN6 TIMP2 IKBKAP KAT6A PYGL IKBKB CASC3 KIF20A PIK3R3 VAPB ACLY TOR1A NOSIP HYOU1 CSRP1 WASHC5 CDK1 CDK5R1 RBM34 NUP62
GRN RHOA RPS5 HSPB1 COL1A1 HSPA8 GAPDH CTSD SPP1 RPS6 ATP1B1 ALDOA MUC1 S100A4 HLA-DRA
0 (72%)
2 (27%)
3 (1%)
5000
4000
3000
2000
1000

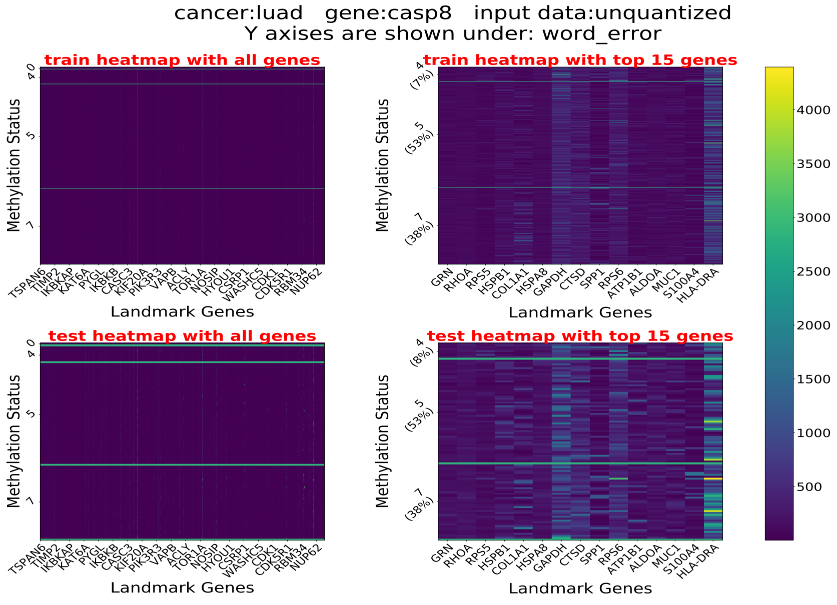
cancer:luad gene:casp8 input data:unquantized
Y axises are shown under: word_error
train heatmap with all genes
train heatmap with top 15 genes
test heatmap with all genes
test heatmap with top 15 genes
Methylation Status
Landmark Genes
TSPAN6 TIMP2 IKBKAP KAT6A PYGL IKBKB CASC3 KIF20A PIK3R3 VAPB ACLY TOR1A NOSIP HYOU1 CSRP1 WASHC5 CDK1 CDK5R1 RBM34 NUP62
GRN RHOA RPS5 HSPB1 COL1A1 HSPA8 GAPDH CTSD SPP1 RPS6 ATP1B1 ALDOA MUC1 S100A4 HLA-DRA
4 (7%)
5 (53%)
7 (38%)
4 (8%)
4000
3500
3000
2500
2000
1500
1000
500

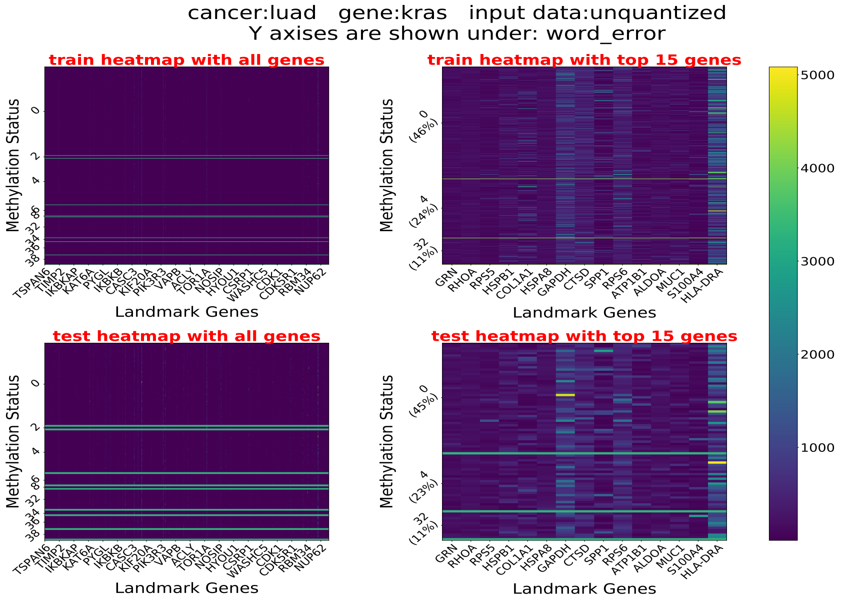
cancer:luad gene:kras input data:unquantized
Y axises are shown under: word_error
train heatmap with all genes
train heatmap with top 15 genes
test heatmap with all genes
test heatmap with top 15 genes
Methylation Status
Landmark Genes
TSPAN6 TIMP2 IKBKAP KAT6A PYGL IKBKB CASC3 KIF20A PIK3R3 VAPB ACLY TOR1A NOSIP HYOU1 CSRP1 WASHC5 CDK1 CDK5R1 RBM34 NUP62
GRN RHOA RPS5 HSPB1 COL1A1 HSPA8 GAPDH CTSD SPP1 RPS6 ATP1B1 ALDOA MUC1 S100A4 HLA-DRA
0 (46%)
4 (24%)
32 (11%)
0 (45%)
4 (23%)
32 (11%)
1000
2000
3000
4000
5000

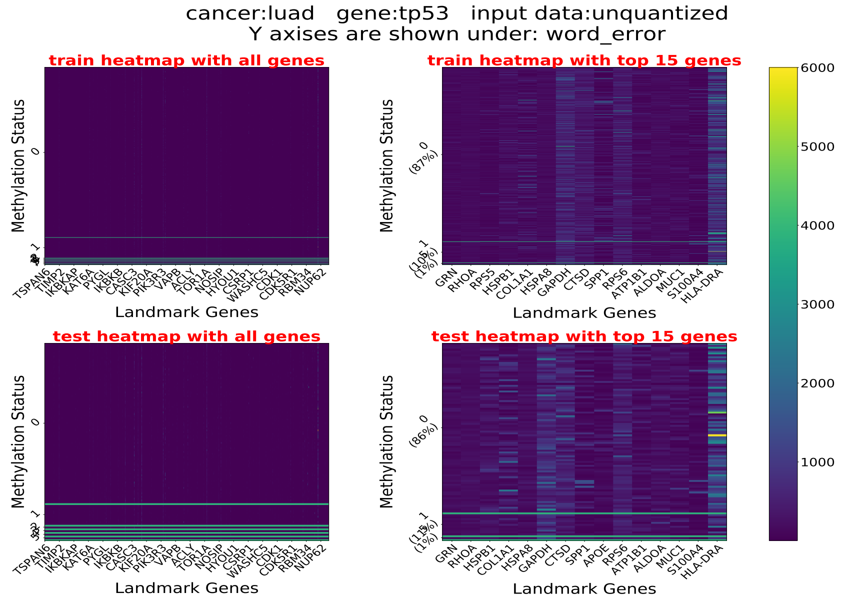
cancer:luad gene:tp53 input data:unquantized
Y axises are shown under: word_error
train heatmap with all genes
train heatmap with top 15 genes
test heatmap with all genes
test heatmap with top 15 genes
Methylation Status
Landmark Genes
TSPAN6 TIMP2 IKBKAP KAT6A PYGL IKBKB CASC3 KIF20A PIK3R3 VAPB ACLY TOR1A NOSIP HYOU1 CSRP1 WASHC5 CDK1 CDK5R1 RBM34 NUP62
GRN RHOA RPS5 HSPB1 COL1A1 HSPA8 GAPDH CTSD SPP1 RPS6 ATP1B1 ALDOA MUC1 S100A4 HLA-DRA
GRN RHOA HSPB1 COL1A1 HSPA8 GAPDH CTSD SPP1 APOE RPS6 ATP1B1 ALDOA MUC1 S100A4 HLA-DRA
0 (87%)
0 (86%)
1000
2000
3000
4000
5000
6000

#### 4.5 LUSC

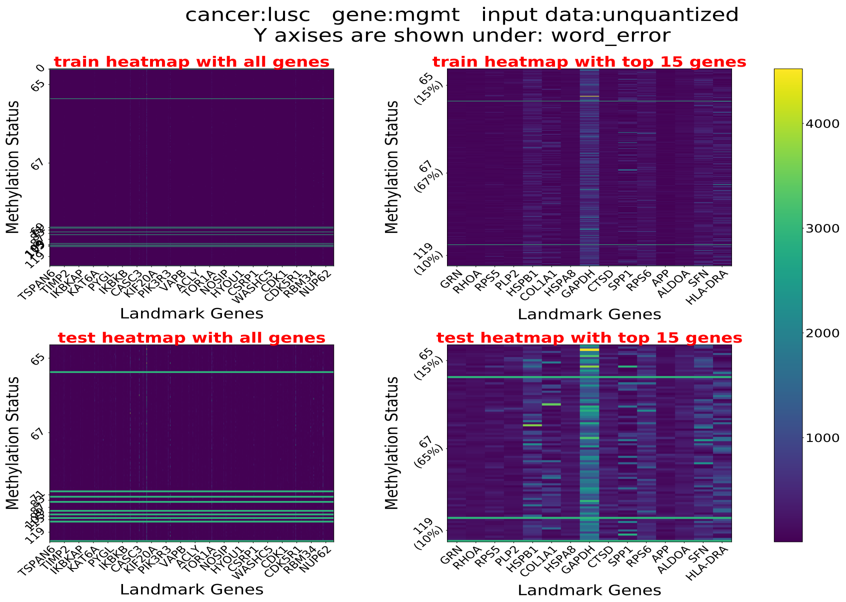
cancer:lusc gene:mgmt input data:unquantized
Y axises are shown under: word_error
train heatmap with all genes
train heatmap with top 15 genes
test heatmap with all genes
test heatmap with top 15 genes
Methylation Status
Landmark Genes
65 (15%)
67 (67%)
119 (10%)
67 (65%)
TSPAN6 TIMP2 IKBKAP KAT6A PYGL IKBKB CASC3 KIF20A PIK3R3 VAPB ACLY TOR1A NOSIP HYOU1 CSRP1 WASHC5 CDK1 CDK5R1 RBM34 NUP62
GRN RHOA RPS5 PLP2 HSPB1 COL1A1 HSPA8 GAPDH CTSD SPP1 RPS6 APP ALDOA SFN HLA-DRA
1000
2000
3000
4000

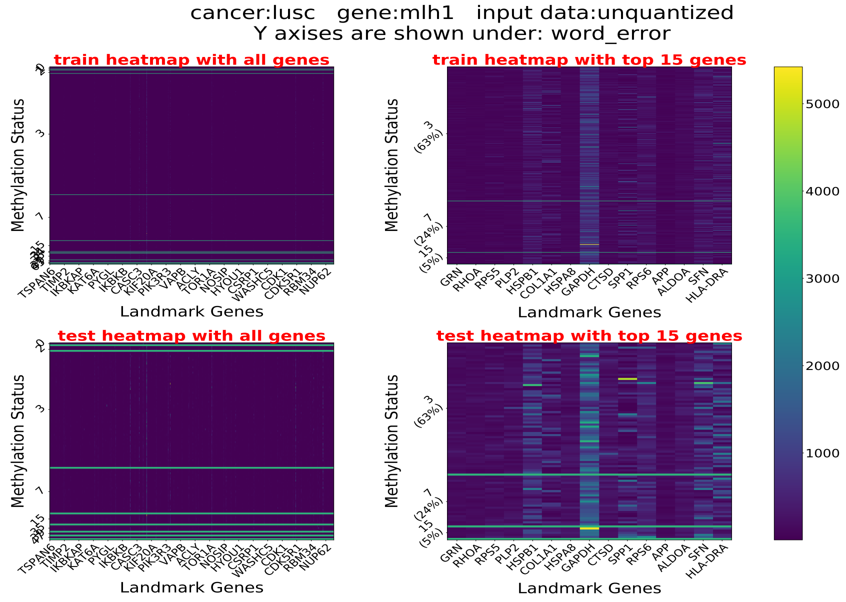
cancer:lusc gene:mlh1 input data:unquantized
Y axises are shown under: word_error
train heatmap with all genes
train heatmap with top 15 genes
test heatmap with all genes
test heatmap with top 15 genes
Methylation Status
Landmark Genes
3 (63%)
7 (24%)
15 (5%)
TSPAN6 TIMP2 IKBKAP KAT6A PYGL IKBKB CASC3 KIF20A PIK3R3 VAPB ACLY TOR1A NOSIP HYOU1 CSRP1 WASHC5 CDK1 CDK5R1 RBM34 NUP62
GRN RHOA RPS5 PLP2 HSPB1 COL1A1 HSPA8 GAPDH CTSD SPP1 RPS6 APP ALDOA SFN HLA-DRA
1000
2000
3000
4000
5000

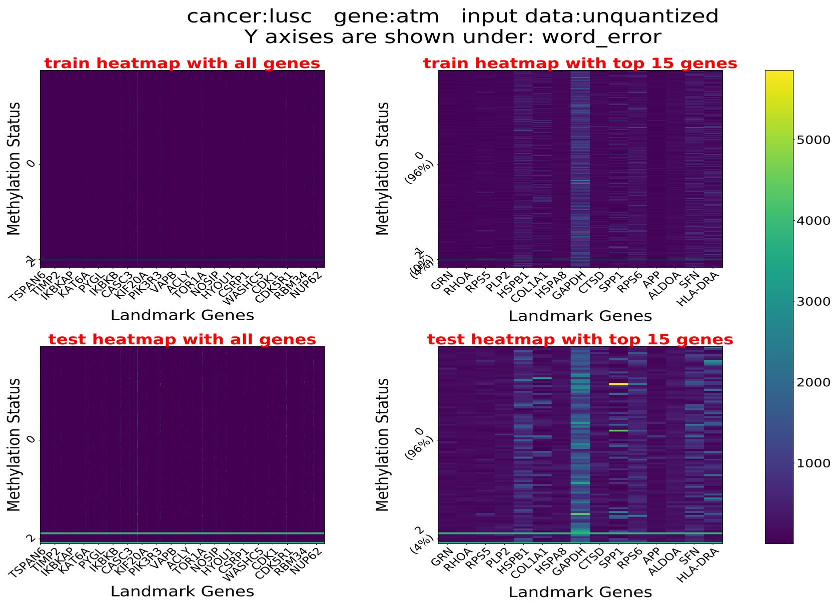
cancer:lusc gene:atm input data:unquantized
Y axises are shown under: word_error
train heatmap with all genes
train heatmap with top 15 genes
test heatmap with all genes
test heatmap with top 15 genes
Methylation Status
Landmark Genes
TSPAN6 TIMP2 IKBKAP KAT6A PYGL IKBKB CASC3 KIF20A PIK3R3 VAPB ACLY TOR1A NOSIP HYOU1 CSRP1 WASHC5 CDK1 CDK5R1 RBM34 NUP62
GRN RHOA RPS5 PLP2 HSPB1 COL1A1 HSPA8 GAPDH CTSD SPP1 RPS6 APP ALDOA SFN HLA-DRA
0 (96%)
2 (4%)
1000 2000 3000 4000 5000

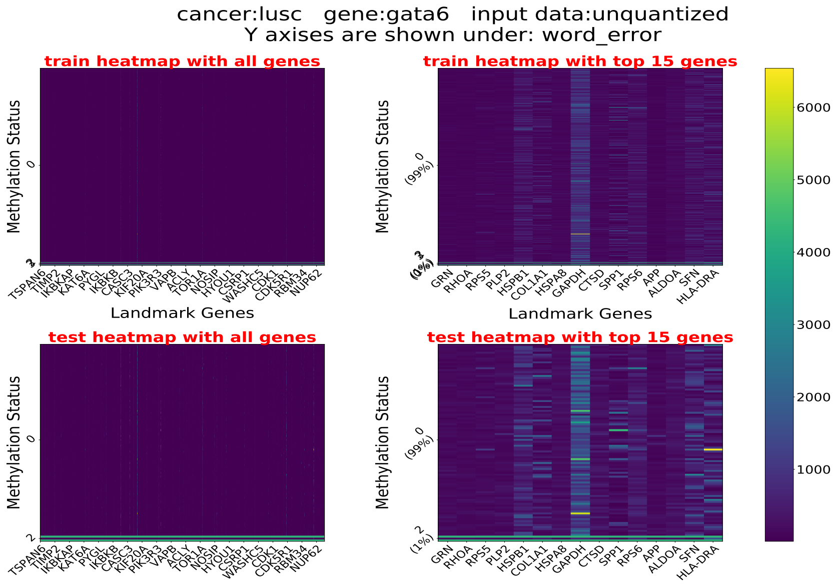
cancer:lusc gene:gata6 input data:unquantized
Y axises are shown under: word_error
train heatmap with all genes
train heatmap with top 15 genes
test heatmap with all genes
test heatmap with top 15 genes
Methylation Status
Landmark Genes
TSPAN6 TIMP2 IKBKAP KAT6A PYGL IKBKB CASC3 KIF20A PIK3R3 VAPB ACLY TOR1A NOSIP HYOU1 CSRP1 WASHC5 CDK1 CDK5R1 RBM34 NUP62
GRN RHOA RPS5 PLP2 HSPB1 COL1A1 HSPA8 GAPDH CTSD SPP1 RPS6 APP ALDOA SFN HLA-DRA
0 (99%)
2 (1%)
1000 2000 3000 4000 5000 6000

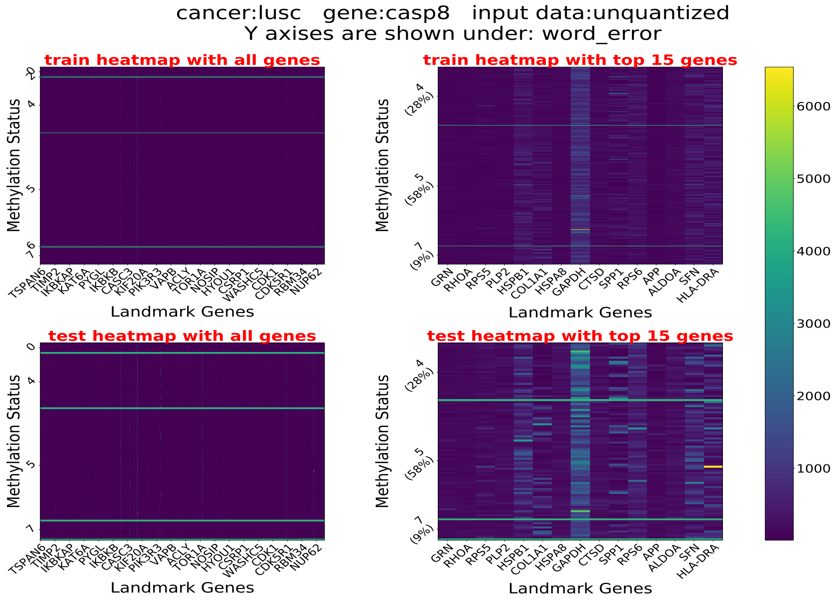
cancer:lusc gene:casp8 input data:unquantized
Y axises are shown under: word_error
train heatmap with all genes
train heatmap with top 15 genes
test heatmap with all genes
test heatmap with top 15 genes
Methylation Status
Landmark Genes
TSPAN6 TIMP2 IKBKAP KAT6A PYGL IKBKB CASC3 KIF20A PIK3R3 VAPB ACLY TOR1A NOSIP HYOU1 CSRP1 WASHC5 CDK1 CDK5R1 RBM34 NUP62
GRN RHOA RPS5 PLP2 HSPB1 COL1A1 HSPA8 GAPDH CTSD SPP1 RPS6 APP ALDOA SFN HLA-DRA
4 (28%)
5 (58%)
7 (9%)
1000 2000 3000 4000 5000 6000

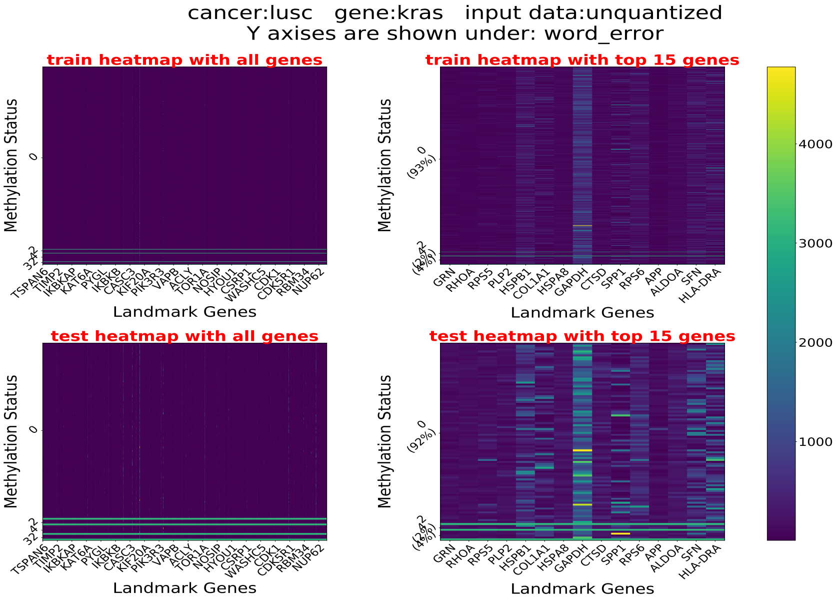
cancer:lusc gene:kras input data:unquantized
Y axises are shown under: word_error
train heatmap with all genes
train heatmap with top 15 genes
test heatmap with all genes
test heatmap with top 15 genes
Methylation Status
Landmark Genes
TSPAN6
TIMP2
IKBKAP
KAT6A
PYGL
IKBKB
CASC3
KIF20A
PIK3R3
VAPB
ACLY
TOR1A
NOSIP
HYOU1
CSRP1
WASHC5
CDK1
CDK5R1
RBM34
NUP62
GRN
RHOA
RPS5
PLP2
HSPB1
COL1A1
HSPA8
GAPDH
CTSD
SPP1
RPS6
APP
ALDOA
SFN
HLA-DRA
0
(93%)
(92%)
4000
3000
2000
1000

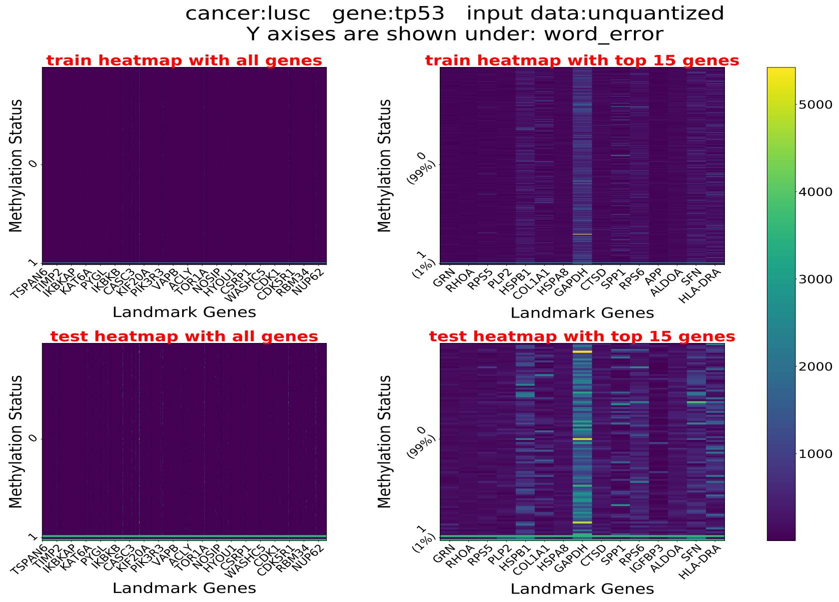
cancer:lusc gene:tp53 input data:unquantized
Y axises are shown under: word_error
train heatmap with all genes
train heatmap with top 15 genes
test heatmap with all genes
test heatmap with top 15 genes
Methylation Status
Landmark Genes
TSPAN6
TIMP2
IKBKAP
KAT6A
PYGL
IKBKB
CASC3
KIF20A
PIK3R3
VAPB
ACLY
TOR1A
NOSIP
HYOU1
CSRP1
WASHC5
CDK1
CDK5R1
RBM34
NUP62
GRN
RHOA
RPS5
PLP2
HSPB1
COL1A1
HSPA8
GAPDH
CTSD
SPP1
RPS6
APP
IGFBP3
ALDOA
SFN
HLA-DRA
0
(99%)
1
(1%)
5000
4000
3000
2000
1000

#### 4.6 Lung Cancer

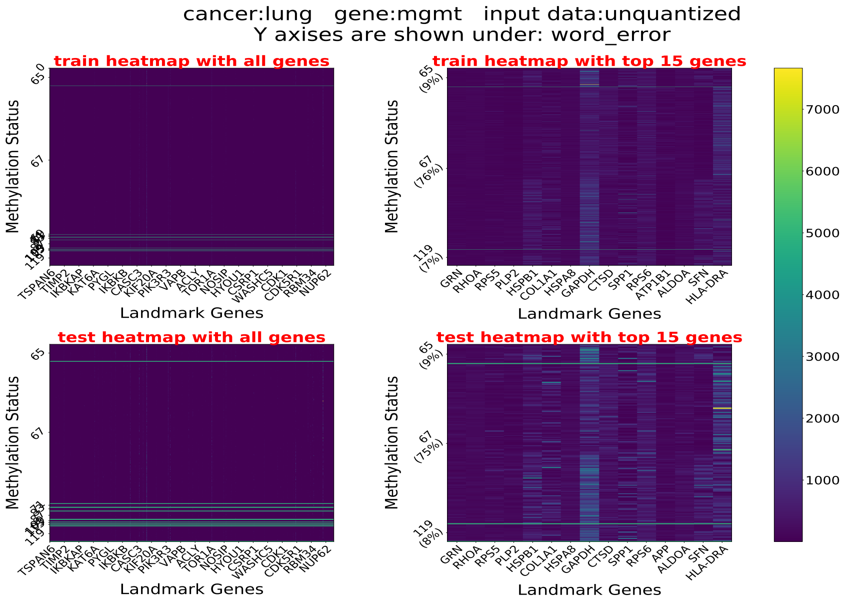

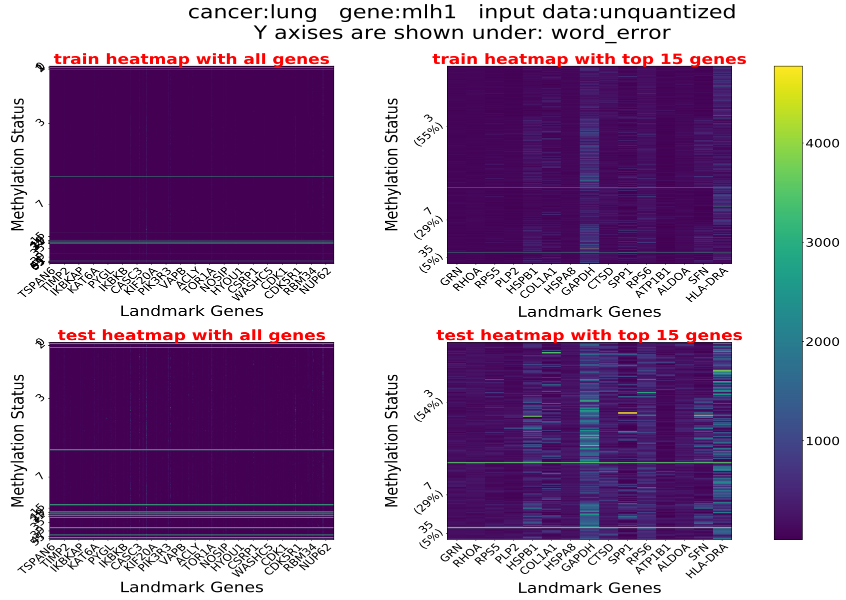

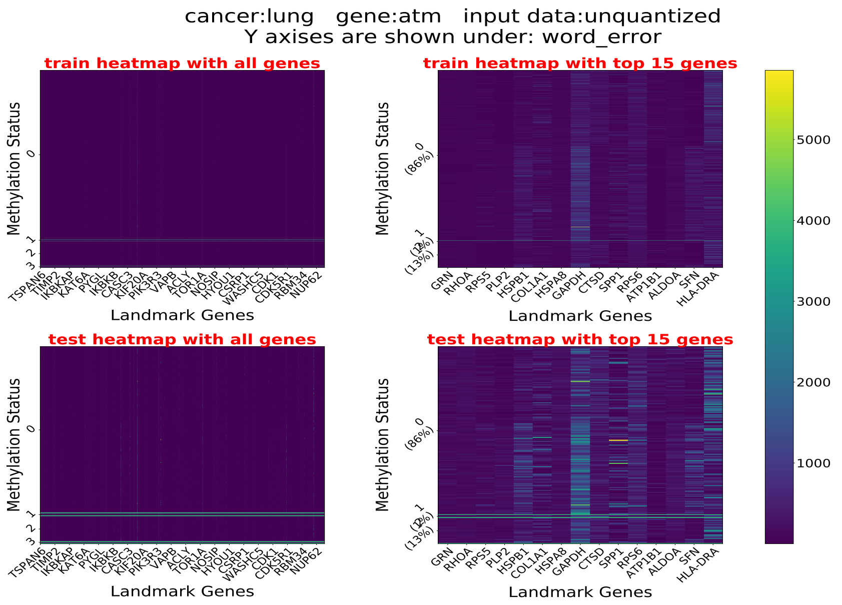
cancer:lung gene:atm input data:unquantized
Y axises are shown under: word_error
train heatmap with all genes
train heatmap with top 15 genes
test heatmap with all genes
test heatmap with top 15 genes
Methylation Status
Landmark Genes
TSPAN6 TIMP2 IKBKAP KAT6A PYGL IKBKB CASC3 KIF20A PIK3R3 VAPB ACLY TOR1A NOSIP HYOU1 CSRP1 WASHC5 CDK1 CDK5R1 RBM34 NUP62
GRN RHOA RPS5 PLP2 HSPB1 COL1A1 HSPA8 GAPDH CTSD SPP1 RPS6 ATP1B1 ALDOA SFN HLA-DRA
0 (86%)
1000 2000 3000 4000 5000

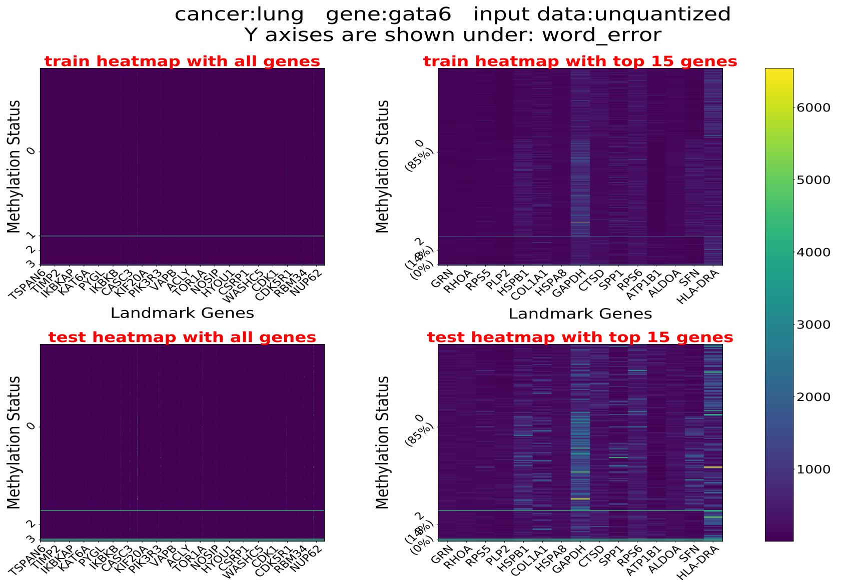
cancer:lung gene:gata6 input data:unquantized
Y axises are shown under: word_error
train heatmap with all genes
train heatmap with top 15 genes
test heatmap with all genes
test heatmap with top 15 genes
Methylation Status
Landmark Genes
TSPAN6 TIMP2 IKBKAP KAT6A PYGL IKBKB CASC3 KIF20A PIK3R3 VAPB ACLY TOR1A NOSIP HYOU1 CSRP1 WASHC5 CDK1 CDK5R1 RBM34 NUP62
GRN RHOA RPS5 PLP2 HSPB1 COL1A1 HSPA8 GAPDH CTSD SPP1 RPS6 ATP1B1 ALDOA SFN HLA-DRA
0 (85%)
1000 2000 3000 4000 5000 6000

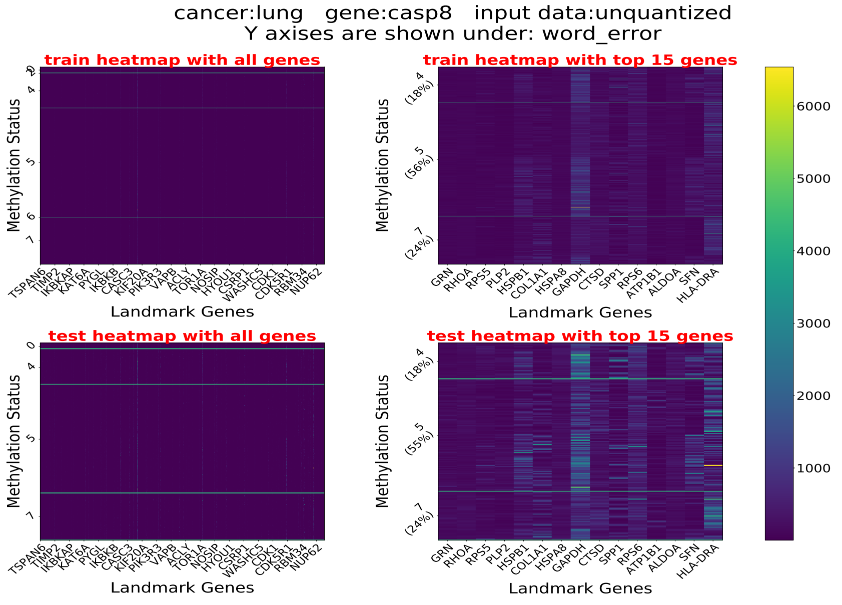
cancer:lung gene:casp8 input data:unquantized
Y axises are shown under: word_error
train heatmap with all genes
train heatmap with top 15 genes
test heatmap with all genes
test heatmap with top 15 genes
Methylation Status
Landmark Genes
TSPAN6 TIMP2 IKBKAP KAT6A PYGL IKBKB CASC3 KIF20A PIK3R3 VAPB ACLY TOR1A NOSIP HYOU1 CSRP1 WASHC5 CDK1 CDK5R1 RBM34 NUP62
GRN RHOA RPS5 PLP2 HSPB1 COL1A1 HSPA8 GAPDH CTSD SPP1 RPS6 ATP1B1 ALDOA SFN HLA-DRA
4 (18%)
5 (56%)
5 (55%)
7 (24%)
1000 2000 3000 4000 5000 6000

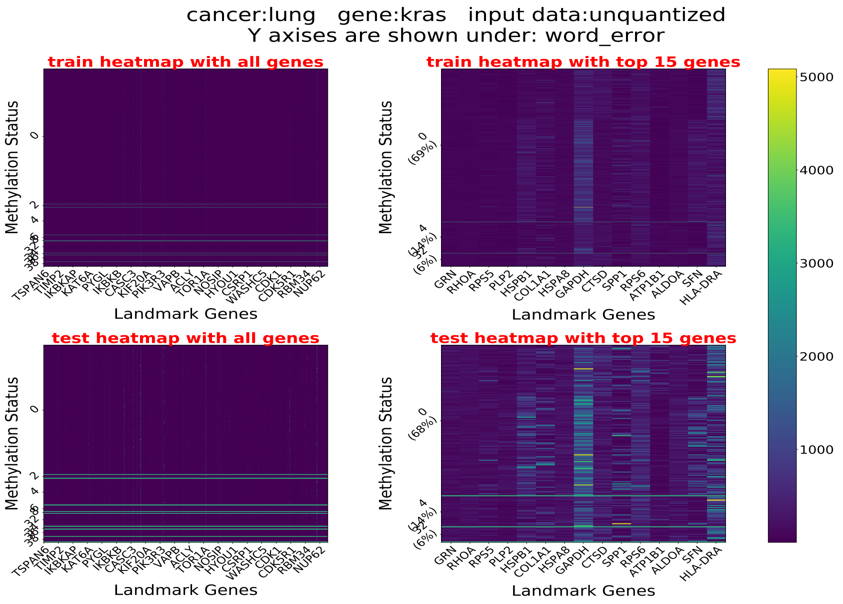
cancer:lung gene:kras input data:unquantized
Y axises are shown under: word_error
train heatmap with all genes
train heatmap with top 15 genes
test heatmap with all genes
test heatmap with top 15 genes
Methylation Status
Landmark Genes
TSPAN6 TIMP2 IKBKAP KAT6A PYGL IKBKB CASC3 KIF20A PIK3R3 VAPB ACLY TOR1A NOSIP HYOU1 CSRP1 WASHC5 CDK1 CDK5R1 RBM34 NUP62
GRN RHOA RPS5 PLP2 HSPB1 COL1A1 HSPA8 GAPDH CTSD SPP1 RPS6 ATP1B1 ALDOA SFN HLA-DRA
0 (69%)
4 (14%)
32 (6%)
0 (68%)
4 (14%)
32 (6%)
5000
4000
3000
2000
1000

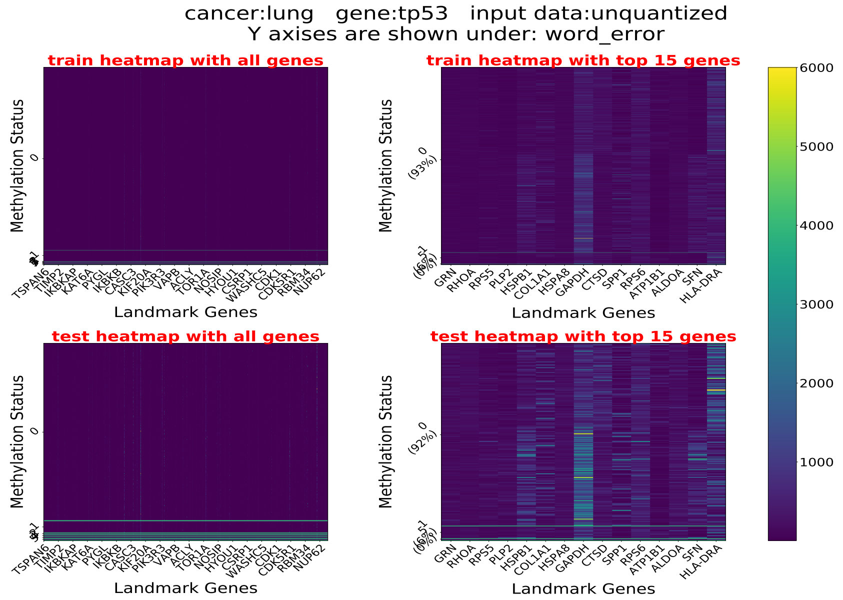
cancer:lung gene:tp53 input data:unquantized
Y axises are shown under: word_error
train heatmap with all genes
train heatmap with top 15 genes
test heatmap with all genes
test heatmap with top 15 genes
Methylation Status
Landmark Genes
TSPAN6 TIMP2 IKBKAP KAT6A PYGL IKBKB CASC3 KIF20A PIK3R3 VAPB ACLY TOR1A NOSIP HYOU1 CSRP1 WASHC5 CDK1 CDK5R1 RBM34 NUP62
GRN RHOA RPS5 PLP2 HSPB1 COL1A1 HSPA8 GAPDH CTSD SPP1 RPS6 ATP1B1 ALDOA SFN HLA-DRA
0 (93%)
0 (92%)
6000
5000
4000
3000
2000
1000

#### 4.7 STAD

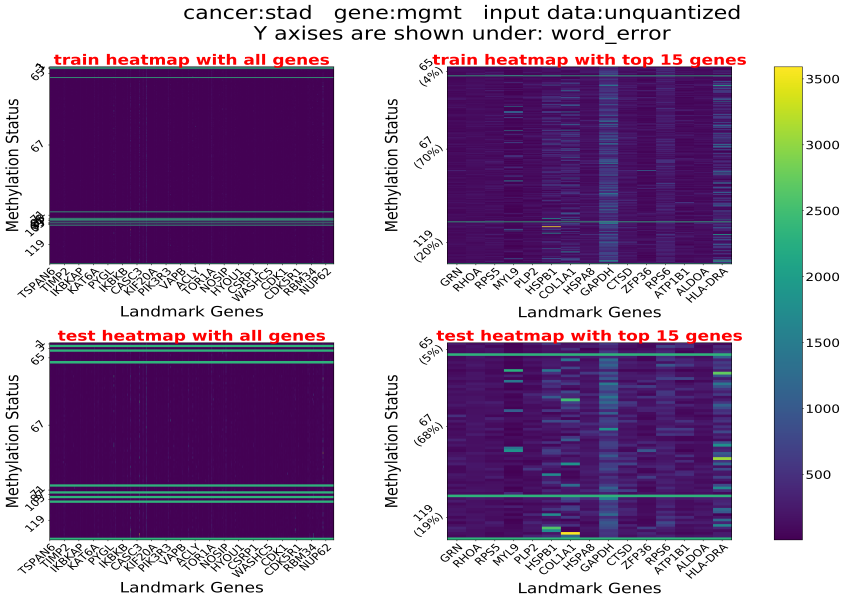
cancer:stad gene:mgmt input data:unquantized
Y axises are shown under: word_error
train heatmap with all genes
train heatmap with top 15 genes
test heatmap with all genes
test heatmap with top 15 genes
Methylation Status
Landmark Genes
TSPAN6 TIMP2 IKBKAP KAT6A PYGL IKBKB CASC3 KIF20A PIK3R3 VAPB ACLY TOR1A NOSIP HYOU1 CSRP1 WASHC5 CDK1 CDK5R1 RBM34 NUP62
GRN RHOA RPS5 MYL9 PLP2 HSPB1 COL1A1 HSPA8 GAPDH CTSD ZFP36 RPS6 ATP1B1 ALDOA HLA-DRA
65 (4%)
67 (70%)
119 (20%)
65 (5%)
67 (68%)
119 (19%)
3500
3000
2500
2000
1500
1000
500

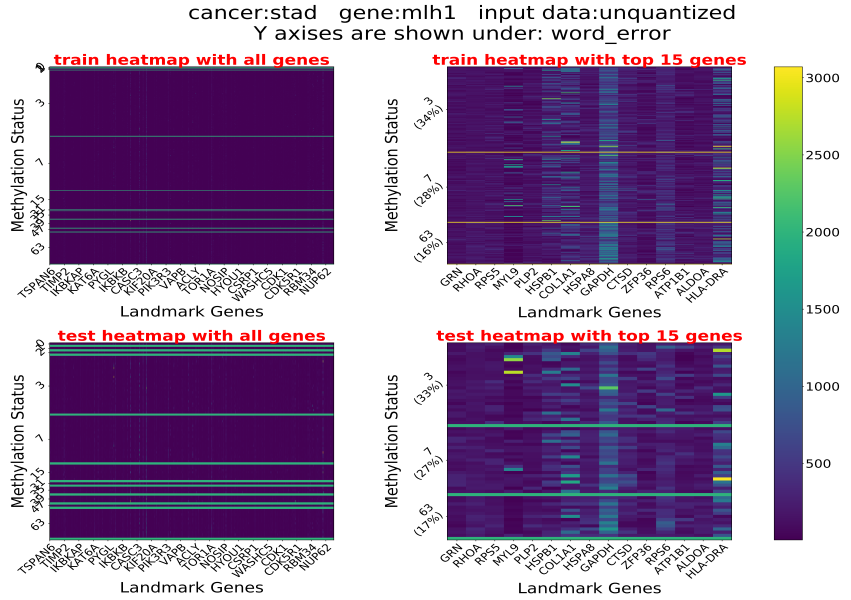
cancer:stad gene:mlh1 input data:unquantized
Y axises are shown under: word_error
train heatmap with all genes
train heatmap with top 15 genes
test heatmap with all genes
test heatmap with top 15 genes
Methylation Status
Landmark Genes
TSPAN6 TIMP2 IKBKAP KAT6A PYGL IKBKB CASC3 KIF20A PIK3R3 VAPB ACLY TOR1A NOSIP HYOU1 CSRP1 WASHC5 CDK1 CDK5R1 RBM34 NUP62
GRN RHOA RPS5 MYL9 PLP2 HSPB1 COL1A1 HSPA8 GAPDH CTSD ZFP36 RPS6 ATP1B1 ALDOA HLA-DRA
3 (34%)
7 (28%)
63 (16%)
3 (33%)
7 (27%)
63 (17%)
3000
2500
2000
1500
1000
500

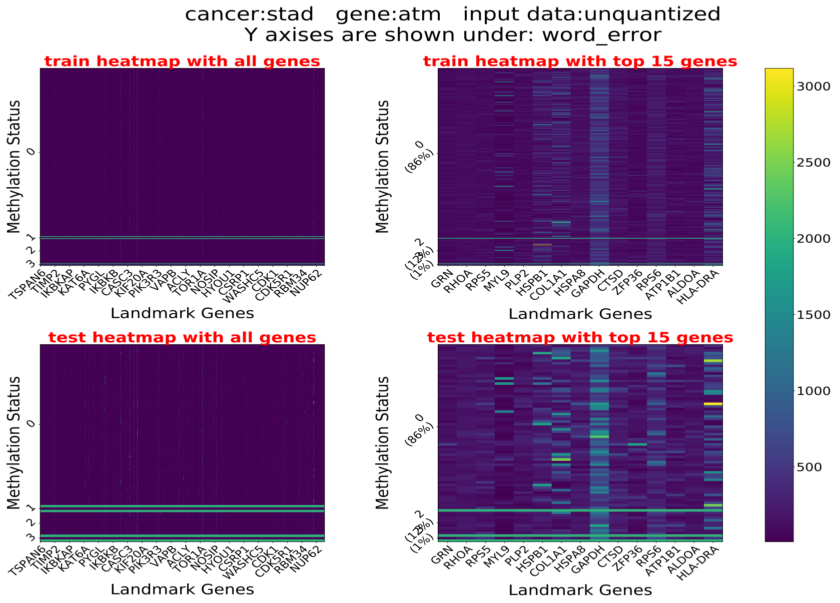
cancer:stad gene:atm input data:unquantized
Y axises are shown under: word_error
train heatmap with all genes
train heatmap with top 15 genes
test heatmap with all genes
test heatmap with top 15 genes
Methylation Status
Landmark Genes
TSPAN6 TIMP2 IKBKAP KAT6A PYGL IKBKB CASC3 KIF20A PIK3R3 VAPB ACLY TOR1A NOSIP HYOU1 CSRP1 WASHC5 CDK1 CDK5R1 RBM34 NUP62
GRN RHOA RPS5 MYL9 PLP2 HSPB1 COL1A1 HSPA8 GAPDH CTSD ZFP36 RPS6 ATP1B1 ALDOA HLA-DRA
0 (86%)
3000
2500
2000
1500
1000
500

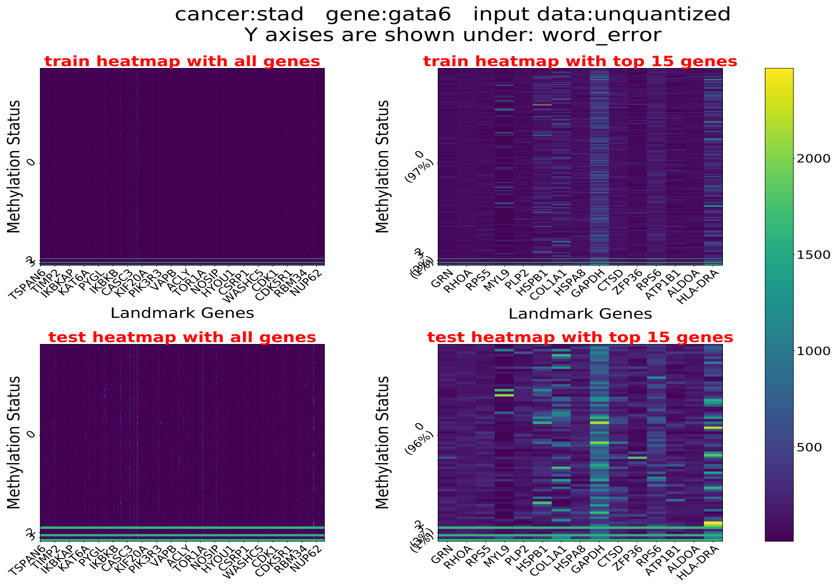
cancer:stad gene:gata6 input data:unquantized
Y axises are shown under: word_error
train heatmap with all genes
train heatmap with top 15 genes
test heatmap with all genes
test heatmap with top 15 genes
Methylation Status
Landmark Genes
TSPAN6 TIMP2 IKBKAP KAT6A PYGL IKBKB CASC3 KIF20A PIK3R3 VAPB ACLY TOR1A NOSIP HYOU1 CSRP1 WASHC5 CDK1 CDK5R1 RBM34 NUP62
GRN RHOA RPS5 MYL9 PLP2 HSPB1 COL1A1 HSPA8 GAPDH CTSD ZFP36 RPS6 ATP1B1 ALDOA HLA-DRA
0 (97%)
0 (96%)
2000
1500
1000
500

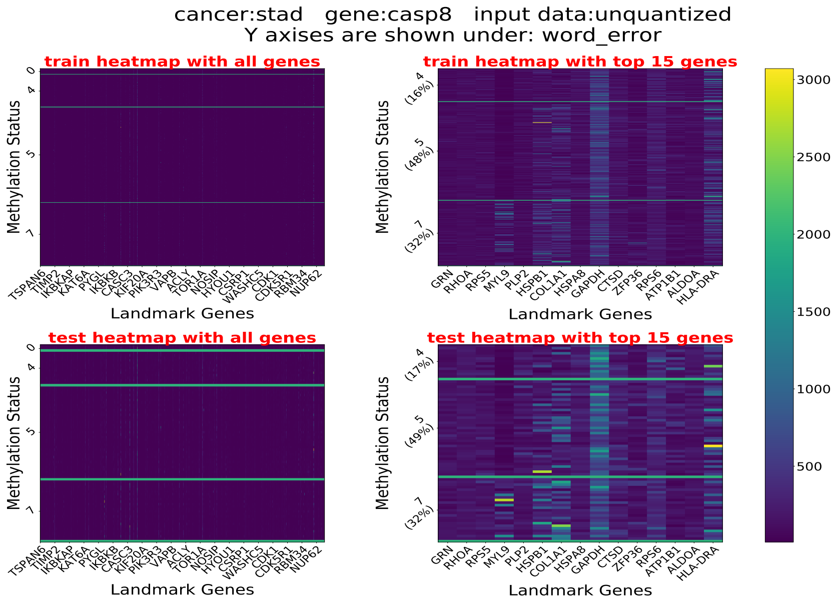
cancer:stad gene:casp8 input data:unquantized
Y axises are shown under: word_error
train heatmap with all genes
train heatmap with top 15 genes
test heatmap with all genes
test heatmap with top 15 genes
Methylation Status
Landmark Genes
TSPAN6 TIMP2 IKBKAP KAT6A PYGL IKBKB CASC3 KIF20A PIK3R3 VAPB ACLY TOR1A NOSIP HYOU1 CSRP1 WASHC5 CDK1 CDK5R1 RBM34 NUP62
GRN RHOA RPS5 MYL9 PLP2 HSPB1 COL1A1 HSPA8 GAPDH CTSD ZFP36 RPS6 ATP1B1 ALDOA HLA-DRA
4 (16%)
5 (48%)
7 (32%)
4 (17%)
5 (49%)
7 (32%)
3000
2500
2000
1500
1000
500

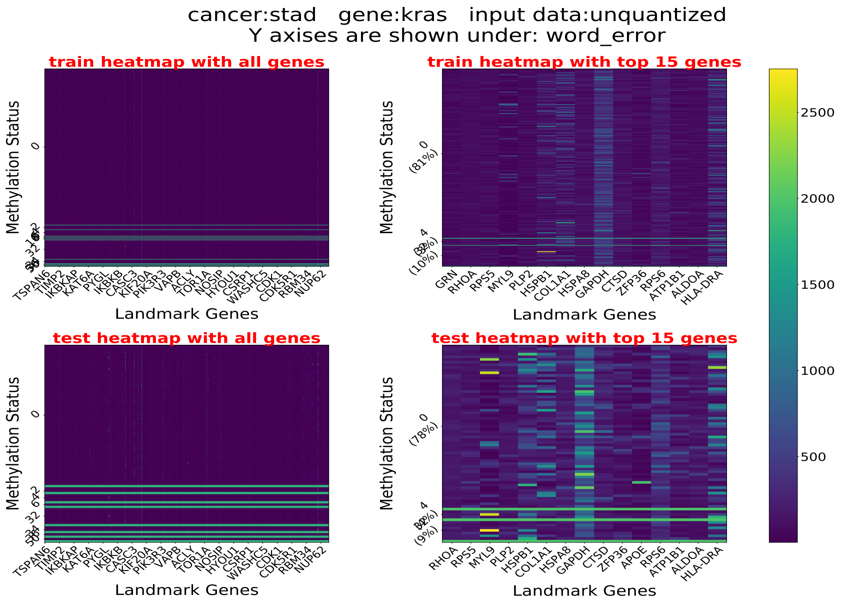
cancer:stad gene:kras input data:unquantized
Y axises are shown under: word_error
train heatmap with all genes
train heatmap with top 15 genes
test heatmap with all genes
test heatmap with top 15 genes
Methylation Status
Landmark Genes
TSPAN6 TIMP2 IKBKAP KAT6A PYGL IKBKB CASC3 KIF20A PIK3R3 VAPB ACLY TOR1A NOSIP HYOU1 CSRP1 WASHC5 CDK1 CDK5R1 RBM34 NUP62
GRN RHOA RPS5 MYL9 PLP2 HSPB1 COL1A1 HSPA8 GAPDH CTSD ZFP36 RPS6 ATP1B1 ALDOA HLA-DRA
RHOA RPS5 MYL9 PLP2 HSPB1 COL1A1 HSPA8 GAPDH CTSD ZFP36 APOE RPS6 ATP1B1 ALDOA HLA-DRA
(81%)
(10%)
(78%)
(9%)
2500
2000
1500
1000
500

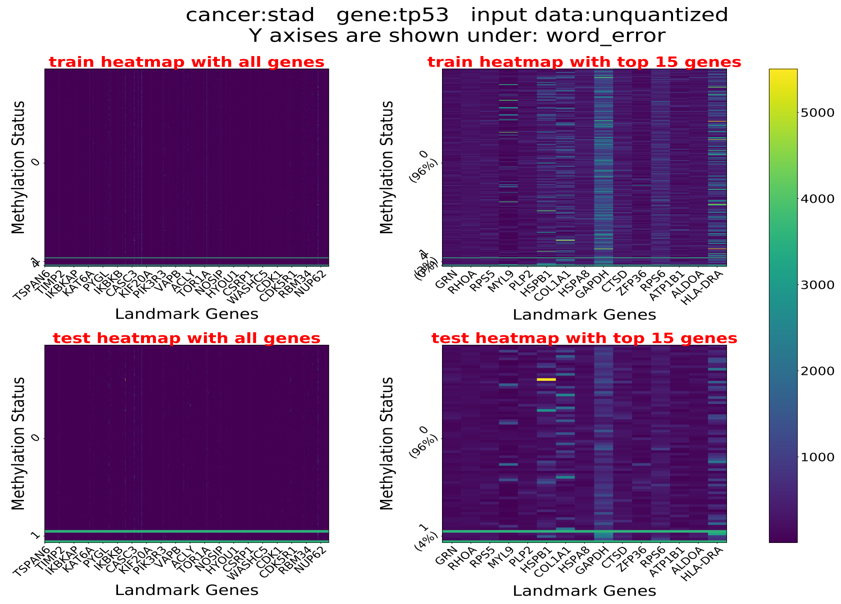
cancer:stad gene:tp53 input data:unquantized
Y axises are shown under: word_error
train heatmap with all genes
train heatmap with top 15 genes
test heatmap with all genes
test heatmap with top 15 genes
Methylation Status
Landmark Genes
TSPAN6 TIMP2 IKBKAP KAT6A PYGL IKBKB CASC3 KIF20A PIK3R3 VAPB ACLY TOR1A NOSIP HYOU1 CSRP1 WASHC5 CDK1 CDK5R1 RBM34 NUP62
GRN RHOA RPS5 MYL9 PLP2 HSPB1 COL1A1 HSPA8 GAPDH CTSD ZFP36 RPS6 ATP1B1 ALDOA HLA-DRA
(96%)
(4%)
5000
4000
3000
2000
1000

#### 5 Prediction Accuracy of MR and FCNN

The presented results include the performance of MR (Multiclass Regression) and 3-layer FCNN (Fully Connected Neural Network) for methylation pattern prediction. In the tables provided, each column represents one gene and each row represents one cancer type. On average, FCNN has a better performance than MR across all genes, as discussed in the main text.

| | MGMT | MLH1 | ATM | GATA6 | CASP8 | KRAS | TP53 |
|---|---|---|---|---|---|---|---|
| LGG | 0.57 | 0.75 | 0.99 | 0.83 | 0.99 | 0.84 | 0.93 |
| GBM | 0.17 | 0.50 | 0.84 | 0.38 | 0.88 | 0.80 | 1.00 |
| BRAIN | 0.47 | 0.69 | 0.97 | 0.79 | 0.96 | 0.85 | 0.94 |
| LUAD | 0.72 | 0.37 | 0.68 | 0.71 | 0.58 | 0.42 | 0.73 |
| LUSC | 0.64 | 0.55 | 0.90 | 0.98 | 0.58 | 0.77 | 0.99 |
| LUNG | 0.71 | 0.53 | 0.75 | 0.69 | 0.59 | 0.61 | 0.75 |
| STAD | 0.61 | 0.43 | 0.76 | 0.96 | 0.59 | 0.70 | 0.93 |
| average | 0.56 | 0.55 | 0.84 | 0.76 | 0.74 | 0.71 | 0.90 |

Table 3: Prediction accuracy of MR for all considered cancer types and genes.

| | MGMT | MLH1 | ATM | GATA6 | CASP8 | KRAS | TP53 |
|---|---|---|---|---|---|---|---|
| LGG | 0.56 | 0.74 | 0.99 | 0.84 | 0.99 | 0.82 | 0.97 |
| GBM | 0.24 | 0.65 | 0.96 | 0.35 | 0.84 | 0.85 | 1.00 |
| BRAIN | 0.46 | 0.71 | 0.98 | 0.82 | 0.96 | 0.85 | 0.97 |
| LUAD | 0.84 | 0.34 | 0.72 | 0.72 | 0.61 | 0.47 | 0.84 |
| LUSC | 0.60 | 0.59 | 0.95 | 0.99 | 0.55 | 0.91 | 0.99 |
| LUNG | 0.72 | 0.50 | 0.85 | 0.85 | 0.56 | 0.66 | 0.91 |
| STAD | 0.64 | 0.40 | 0.84 | 0.96 | 0.64 | 0.78 | 0.96 |
| average | 0.58 | 0.56 | 0.90 | 0.79 | 0.74 | 0.76 | 0.95 |

Table 4: Prediction accuracy of FCNN for all considered cancer types and genes.

#### 6 Prediction Accuracy of Quantized-$E^2M$

The presented results include the performance of Quantized-$E^2M$, with 4-bit and 8-bit uniformly quantized input expressions, and 8-bit and 16-bit uniformly quantized network weights, for all chosen genes and all cancer types.

#### 6.1 Results for Unquantized Expression Data

| | MGMT | MLH1 | ATM | GATA6 | CASP8 | KRAS | TP53 |
|---|---|---|---|---|---|---|---|
| LGG | 0.56 | 0.75 | 0.99 | 0.90 | 0.99 | 0.84 | 0.96 |
| GBM | 0.31 | 0.61 | 0.92 | 0.58 | 0.88 | 0.80 | 1.00 |
| BRAIN | 0.53 | 0.69 | 0.98 | 0.81 | 0.98 | 0.84 | 0.98 |
| LUAD | 0.83 | 0.42 | 0.73 | 0.69 | 0.62 | 0.38 | 0.84 |
| LUSC | 0.64 | 0.65 | 0.94 | 0.99 | 0.63 | 0.92 | 0.99 |
| LUNG | 0.71 | 0.53 | 0.85 | 0.84 | 0.58 | 0.68 | 0.90 |
| STAD | 0.65 | 0.44 | 0.84 | 0.96 | 0.63 | 0.79 | 0.95 |
| average | 0.60 | 0.58 | 0.89 | 0.82 | 0.76 | 0.75 | 0.95 |

Table 5: Prediction accuracy of $E^2M$ for unquantized input expression data and full-precision weights.

| | MGMT | MLH1 | ATM | GATA6 | CASP8 | KRAS | TP53 |
|---|---|---|---|---|---|---|---|
| LGG | 0.55 | 0.73 | 0.99 | 0.90 | 0.99 | 0.82 | 0.96 |
| GBM | 0.27 | 0.56 | 0.91 | 0.53 | 0.88 | 0.77 | 1.00 |
| BRAIN | 0.49 | 0.69 | 0.98 | 0.81 | 0.98 | 0.83 | 0.98 |
| LUAD | 0.82 | 0.42 | 0.71 | 0.68 | 0.59 | 0.36 | 0.84 |
| LUSC | 0.62 | 0.62 | 0.94 | 0.99 | 0.59 | 0.92 | 0.99 |
| LUNG | 0.71 | 0.50 | 0.84 | 0.83 | 0.57 | 0.67 | 0.90 |
| STAD | 0.65 | 0.36 | 0.84 | 0.96 | 0.63 | 0.78 | 0.94 |
| average | 0.59 | 0.55 | 0.89 | 0.81 | 0.75 | 0.74 | 0.94 |

Table 6: Prediction accuracy of $E^2M$ for unquantized input expression data and 16-bit uniformly quantized weights.

| | MGMT | MLH1 | ATM | GATA6 | CASP8 | KRAS | TP53 |
|---|---|---|---|---|---|---|---|
| LGG | 0.37 | 0.64 | 0.99 | 0.86 | 0.99 | 0.84 | 0.96 |
| GBM | 0.24 | 0.52 | 0.96 | 0.40 | 0.15 | 0.87 | 1.00 |
| BRAIN | 0.11 | 0.66 | 0.28 | 0.73 | 0.98 | 0.83 | 0.96 |
| LUAD | 0.79 | 0.39 | 0.76 | 0.66 | 0.48 | 0.38 | 0.73 |
| LUSC | 0.55 | 0.59 | 0.47 | 0.99 | 0.39 | 0.92 | 0.99 |
| LUNG | 0.43 | 0.47 | 0.24 | 0.81 | 0.51 | 0.63 | 0.90 |
| STAD | 0.62 | 0.28 | 0.85 | 0.95 | 0.29 | 0.14 | 0.93 |
| average | 0.44 | 0.51 | 0.65 | 0.77 | 0.54 | 0.66 | 0.92 |

Table 7: Prediction accuracy of $E^2M$ for unquantized input expression data and 8-bit uniformly quantized weights.

#### 6.2 Results for 8-bit Uniformly Quantized Expression Data

| | MGMT | MLH1 | ATM | GATA6 | CASP8 | KRAS | TP53 |
|---|---|---|---|---|---|---|---|
| LGG | 0.57 | 0.73 | 0.99 | 0.89 | 0.99 | 0.83 | 0.97 |
| GBM | 0.24 | 0.65 | 0.96 | 0.31 | 0.92 | 0.88 | 1.00 |
| BRAIN | 0.49 | 0.71 | 0.98 | 0.79 | 0.98 | 0.85 | 0.97 |
| LUAD | 0.84 | 0.43 | 0.61 | 0.68 | 0.55 | 0.46 | 0.83 |
| LUSC | 0.64 | 0.65 | 0.96 | 0.99 | 0.64 | 0.92 | 0.99 |
| LUNG | 0.76 | 0.59 | 0.84 | 0.85 | 0.54 | 0.70 | 0.91 |
| STAD | 0.64 | 0.41 | 0.86 | 0.95 | 0.63 | 0.77 | 0.96 |
| average | 0.60 | 0.60 | 0.89 | 0.78 | 0.75 | 0.77 | 0.95 |

Table 8: Prediction accuracy of $E^2M$ with 8-bit uniformly quantized input data and full-precision weights.

| | MGMT | MLH1 | ATM | GATA6 | CASP8 | KRAS | TP53 |
|---|---|---|---|---|---|---|---|
| LGG | 0.55 | 0.71 | 0.99 | 0.90 | 0.99 | 0.80 | 0.97 |
| GBM | 0.24 | 0.52 | 0.96 | 0.36 | 0.92 | 0.86 | 1.00 |
| BRAIN | 0.47 | 0.69 | 0.98 | 0.76 | 0.98 | 0.85 | 0.97 |
| LUAD | 0.82 | 0.43 | 0.64 | 0.69 | 0.55 | 0.41 | 0.81 |
| LUSC | 0.61 | 0.61 | 0.96 | 0.99 | 0.61 | 0.92 | 0.99 |
| LUNG | 0.73 | 0.54 | 0.84 | 0.81 | 0.44 | 0.67 | 0.91 |
| STAD | 0.61 | 0.37 | 0.85 | 0.95 | 0.59 | 0.76 | 0.96 |
| average | 0.57 | 0.55 | 0.89 | 0.78 | 0.73 | 0.75 | 0.94 |

Table 9: Prediction accuracy of $E^2M$ with 8-bit uniformly quantized input data and 16-bit uniformly quantized weights.

| | MGMT | MLH1 | ATM | GATA6 | CASP8 | KRAS | TP53 |
|---|---|---|---|---|---|---|---|
| LGG | 0.53 | 0.71 | 0.99 | 0.89 | 0.99 | 0.80 | 0.97 |
| GBM | 0.24 | 0.53 | 0.96 | 0.37 | 0.92 | 0.87 | 1.00 |
| BRAIN | 0.41 | 0.70 | 0.98 | 0.77 | 0.98 | 0.85 | 0.97 |
| LUAD | 0.82 | 0.45 | 0.64 | 0.68 | 0.54 | 0.45 | 0.83 |
| LUSC | 0.61 | 0.63 | 0.96 | 0.99 | 0.58 | 0.92 | 0.99 |
| LUNG | 0.73 | 0.53 | 0.84 | 0.82 | 0.44 | 0.69 | 0.91 |
| STAD | 0.61 | 0.36 | 0.85 | 0.96 | 0.60 | 0.77 | 0.96 |
| average | 0.57 | 0.56 | 0.89 | 0.78 | 0.72 | 0.76 | 0.95 |

Table 10: Prediction accuracy of $E^2M$ with 8-bit uniformly quantized input data and 8-bit uniformly quantized weights.

#### 6.3 Results for 4-bit Uniformly Quantized Expression Data

| | MGMT | MLH1 | ATM | GATA6 | CASP8 | KRAS | TP53 |
|---|---|---|---|---|---|---|---|
| LGG | 0.56 | 0.72 | 0.99 | 0.89 | 0.99 | 0.80 | 0.97 |
| GBM | 0.28 | 0.65 | 0.96 | 0.42 | 0.92 | 0.88 | 1.00 |
| BRAIN | 0.54 | 0.68 | 0.98 | 0.80 | 0.98 | 0.84 | 0.98 |
| LUAD | 0.84 | 0.42 | 0.76 | 0.62 | 0.51 | 0.40 | 0.84 |
| LUSC | 0.64 | 0.65 | 0.95 | 0.99 | 0.64 | 0.92 | 0.99 |
| LUNG | 0.71 | 0.60 | 0.84 | 0.82 | 0.55 | 0.64 | 0.99 |
| STAD | 0.68 | 0.46 | 0.86 | 0.96 | 0.61 | 0.78 | 0.96 |
| average | 0.60 | 0.60 | 0.91 | 0.79 | 0.74 | 0.75 | 0.96 |

Table 11: Prediction accuracy of $E^2M$ with 4-bit uniformly quantized input data and full-precision weights.

| | MGMT | MLH1 | ATM | GATA6 | CASP8 | KRAS | TP53 |
|---|---|---|---|---|---|---|---|
| LGG | 0.54 | 0.70 | 0.99 | 0.90 | 0.99 | 0.78 | 0.96 |
| GBM | 0.24 | 0.60 | 0.96 | 0.41 | 0.92 | 0.86 | 1.00 |
| BRAIN | 0.48 | 0.67 | 0.98 | 0.79 | 0.98 | 0.83 | 0.98 |
| LUAD | 0.80 | 0.42 | 0.75 | 0.61 | 0.50 | 0.36 | 0.83 |
| LUSC | 0.59 | 0.63 | 0.95 | 0.99 | 0.59 | 0.91 | 0.99 |
| LUNG | 0.71 | 0.58 | 0.83 | 0.82 | 0.51 | 0.63 | 0.92 |
| STAD | 0.64 | 0.46 | 0.85 | 0.96 | 0.60 | 0.76 | 0.96 |
| average | 0.57 | 0.58 | 0.90 | 0.78 | 0.73 | 0.73 | 0.95 |

Table 12: Prediction accuracy of $E^2M$ with 4-bit uniformly quantized input data and 16-bit uniformly quantized weights.

| | MGMT | MLH1 | ATM | GATA6 | CASP8 | KRAS | TP53 |
|---|---|---|---|---|---|---|---|
| LGG | 0.54 | 0.70 | 0.99 | 0.90 | 0.99 | 0.79 | 0.96 |
| GBM | 0.25 | 0.60 | 0.96 | 0.42 | 0.92 | 0.86 | 1.00 |
| BRAIN | 0.49 | 0.66 | 0.98 | 0.80 | 0.98 | 0.83 | 0.98 |
| LUAD | 0.81 | 0.42 | 0.74 | 0.61 | 0.50 | 0.36 | 0.83 |
| LUSC | 0.60 | 0.65 | 0.95 | 0.99 | 0.59 | 0.92 | 0.99 |
| LUNG | 0.71 | 0.58 | 0.84 | 0.82 | 0.51 | 0.63 | 0.92 |
| STAD | 0.65 | 0.46 | 0.86 | 0.96 | 0.60 | 0.76 | 0.96 |
| average | 0.58 | 0.58 | 0.90 | 0.78 | 0.73 | 0.74 | 0.95 |

Table 13: Prediction accuracy of $E^2M$ with 4-bit uniformly quantized input data and 8-bit uniformly quantized weights.
